# TamL is a key player of the outer membrane homeostasis in Bacteroidetes

**DOI:** 10.1101/2024.10.17.618404

**Authors:** Fabio Giovannercole, Tom de Smet, Miguel Angel Vences-Guzmán, Frédéric Lauber, Rémy Dugauquier, Marc Dieu, Laura Lizen, Jonas Dehairs, Gipsi Lima-Mendez, Ziqiang Guan, Christian Sohlenkamp, Francesco Renzi

## Abstract

In Proteobacteria, the outer membrane protein TamA and the inner membrane-anchored protein TamB form the Translocation and Assembly Module (TAM) complex, which facilitates the transport of autotransporters, virulence factors, and likely lipids across the two membranes.

In Bacteroidetes TamA is replaced by TamL, a TamA-like lipoprotein with a lipid modification at its N-terminus that likely anchors it to the outer membrane. This structural difference suggests that TamL may have a distinct function compared to TamA. However, the role of TAM in bacterial phyla other than Proteobacteria remains unexplored.

Our study aimed to elucidate the functional importance of TamL in *Flavobacterium johnsoniae*, an environmental Bacteroidetes. Unlike its homologues in Proteobacteria, we found that TamL and TamB are essential in *F. johnsoniae*. Through genetic, phenotypic, proteomic, and lipidomic analyses, we discovered that TamL depletion severely compromises outer membrane integrity, as evidenced by reduced cell viability, altered cell shape, increased susceptibility to membrane-disrupting agents, and elevated levels of outer membrane lipoproteins. Notably, we did not observe any impact on outer membrane lipid composition.

Via pull-down protein assays, we confirmed that TamL interacts with TamB in *F. johnsoniae*, likely forming the TAM complex. Furthermore, our *in silico* analysis revealed that the presence of TamL and TamB monocistronic genes is a shared genetic feature among Bacteroidetes members, including the human pathogen *Capnocytophaga canimorsus* where we also confirmed the essentiality of the TamL and TamB homologs.

To our knowledge, this study is the first to provide functional insights into a TAM subunit beyond Proteobacteria.

**Significance:** In Proteobacteria, the outer membrane (OM) protein TamA forms with the inner membrane (IM)-anchored protein TamB the Translocation and Assembly Module Complex (TAM). which contributes to efficient biogenesis of the OM. In Bacteroidetes TamA is replaced by TamL, a TamA-like lipoprotein of unknown role. In this work, we studied TamL in the Bacteroidetes *Flavobacterium johnsoniae*. We found that TamL and TamB are essential for cell viability, and that TamL depletion disrupts outer membrane stability, increases outer membrane vesicle size, and lead to higher sensitivity to OM stressors. These findings highlight TamL critical role in maintaining OM structure in Bacteroidetes. To our surprise, we also identified multiple TamL, TamB and TamA homologs in Bacteroidetes. Altogether, our findings extend the current knowledge on TAM and provide novel insights into a field of research barely investigated outside Proteobacteria.

## Introduction

All gram-negative (diderm) bacteria possess an outer membrane (OM) surrounding the periplasm, the aqueous compartment that separates the OM from the inner membrane (IM) [1,2]. In most diderm bacteria, the OM is made of phospholipids (PLs) and lipopolysaccharide (LPS), whose distribution is asymmetrical along the OM, with PLs and LPS located in the inner and outer leaflets, respectively [2]. Additionally, the OM is enriched with proteins, such as integral OM proteins (OMPs) that adopt a characteristic β-barrel fold, and lipoproteins, which are globular proteins anchored to the OM via a lipid moiety [3]. Both OMPs and OM lipoproteins are synthesized in the cytoplasm as precursors and are then translocated across the periplasm towards the OM thanks to dedicated shuttle systems, such as the periplasmic protein LolA in the case of OM lipoproteins or the periplasmic chaperones Skp, SurA and DegP for OMPs [4–6]. After reaching the OM, the insertion and folding of OMPs into the OM is catalyzed by members of the Omp85 family [7,8], whereas OM lipoproteins are inserted in the OM by LolB [3,5].

The OMP BamA is the best characterized member of the Omp85 family and the main subunit of the BAM (β-Barrel Assembly Machinery) complex, which assembles OMPs in all diderm bacteria [8,9]. However, components of a family of outer membrane/secreted proteins known as autotransporters seem to rely on an auxiliary complex, called Translocation and Assembly Module (TAM) [10–12]. TAM is composed of TamA, a BamA homologue, which is the OM foldase and insertase of the complex, and of an IM-anchored periplasmic protein TamB, which likely conveys nascent and unfolded OMPs from the IM to TamA [13,14]. The full-length structure of TamA has been determined [15,16] and, likewise members of the Omp85 family, TamA displays a C-terminal β-barrel domain made of 16 antiparallel β-sheets, and 3 highly rigid N-terminal POlypeptide TRansport Associated (POTRA) domains extending towards the periplasm and adopting a C-shaped arrangement [14,15]. As for TamB, only the structure of a limited portion (residues 977-1136) of the *Escherichia coli* TamB has been experimentally solved, which folds into a hydrophobic β-taco motif consisting of β-sheets and random coils [17]. Structural predictions indicate that the whole periplasmic portion of TamB likely consists of repetitions of this fold, which would create a periplasmic tube-like structure, whose interior cavity would be large and hydrophobic enough to accommodate an unfolded polypeptide chain to chaperone to TamA [10,11]. Furthermore, the fold adopted by the last six β-strands of the C-terminal end of TamB mirrors that of OMP β-barrels, suggesting a pseudosubstrate function. This pseudosubstrate domain likely interacts with the β-barrel domain of TamA until high affinity substrates bind [10,14].

TamB belongs to a family known as AsmA-like proteins, which in *E. coli* includes five additional members, AsmA, YicH, YhjG, YhdP and YdbH, all anchored to the IM [18]. These six AsmA-like proteins have in common the periplasmic β-taco fold, and thereby their tube-like structures resemble that of the eukaryotic repeated β-groove (RBG) lipid-transfer proteins [19–21]. In fact, recent experimental evidence has shown that YhdP, YdbH and TamB could be implicated in the anterograde (IM to OM) transports of PLs [22–27]. To date, the structure and function of TAM have been investigated in depth exclusively in Proteobacteria, where the genes coding for TamA and TamB are part of the same operon [28,29]. Nevertheless, members of the Bacteroidetes and Chlorobi phyla encode TamL, a TamA-like lipoprotein. In fact, TamL homologs all possess a characteristic lipoprotein signal peptide harboring a conserved cysteine to which lipid moieties are attached during maturation of the protein. This suggests that the TamL periplasmic POTRA domain are anchored to the OM, rather than freely facing the periplasm [29]. Furthermore, it must be pointed out that in general the OM lipid and protein composition in Bacteroidetes significantly differs from that of Proteobacteria [30–34]. These biochemical features raise therefore numerous questions about TamL function and its ability to interact with TamB in Bacteroidetes.

In this work we aimed to investigate the biological role of TamL in OM homeostasis and biogenesis of the Bacteroidetes *Flavobacterium johnsoniae*, an environmental and model bacterium for gliding and Type 9 secretion system (T9SS) [35–40]. Our data show that in *F. johnsoniae* the protein TamL is essential, and that its depletion drastically affects OM integrity, induces blebs with release of larger outer membrane vesicles (OMVs), and drastically perturbs the OM proteome composition, while having no significant impact on the OM lipidome. Additionally, we show that TamL interacts with TamB and thereby likely forms a TAM complex. We also confirmed the essentiality of TamL in *Capnocytophaga canimorsus,* an oral commensal in cats and dogs but also a human pathogen [31,41,42]. Finally, we found that, unlike *tamA* and *tamB* in Proteobacteria, the genes *tamL* and *tamB* in Bacteroidetes are not part of the same operon, which seems to be a genetic feature exclusive to members of this phylum.

In this study we provide the first insights into the biological role played by a TamA-like protein in OM homeostasis and extend our understanding of the function of an Omp85 family member outside Proteobacteria.

## Results

### *F. johnsoniae* possesses several TamL and TamB proteins

We first searched for a putative TamL homologue in *F. johnsoniae* by DELTA-BLAST [43] using TamA from *E. coli* (*Ec*TamA, UniProt code: P0ADE4) as a query. In addition to BamA (*fjoh_1640*), which was annotated as “Surface antigen (D15)” and had the highest score, we identified two proteins of interest annotated as “BamA/TamA family outer membrane protein”, encoded by the genes *fjoh_1900* and *fjoh_1464*, and one protein annotated as “Membrane protein” encoded by the gene *fjoh_0402* (Table 1). A closer inspection of their AlphaFold2-predicted structures revealed the presence of three N-terminal POTRA domains and of a C-terminal β-barrel made of 16 β-sheets for all the homologues, alike *Ec*TamA (Figure 1A). Interestingly, when checked for the presence of a signal peptide (SP) by SignalP 6.0 [44], we found that the *fjoh_1900* and *fjoh_1464* encoded proteins were predicted with high likelihood (0.99 and >0.83, respectively) to have a lipoprotein signal peptide (SPII), with a cysteine, Cys27 (Fjoh_1900) and Cys21 (Fjoh_1464), preceding the POTRA1 domains being the predicted lipidated residue. As a result, we concluded that both *fjoh_1900* and *fjoh_1464* likely code for a TamL protein. In contrast, a SPI signal peptide (likelihood >0.99) was observed for the protein encoded by *fjoh_0402*, which suggested that this gene may code for a putative TamA protein. This was surprising since, to our knowledge, *tamA* is nearly exclusively confined to Proteobacteria, where it is in operon with *tamB* [28,29]. Conversely, we did not identify any TamB-encoding gene flanking *fjoh_0402*.

**Table 1.** DELTA-BLAST [43] results using *Ec*TamA, *Ec*TamB or *Ec*AsmA as queries.

| Annotation | Total score | Query cover | E-value | % identity | Accession length | Accession | Protein number |
| --- | --- | --- | --- | --- | --- | --- | --- |
| ***EcTamA*** |  |  |  |  |  |  |  |
| Surface antigen (D15) | 271 | 96% | 5e-40 | 13.34 | 900 | ABQ04722.1 | Fjoh_1640 |
| BamA/TamA family outer membrane protein | 174 | 88% | 8e-30 | 12.52 | 772 | WP_012023976.1 | Fjoh_1900 |
| Membrane protein | 88.2 | 88% | 5e-19 | 13.78 | 566 | WP_012022494.1 | Fjoh_0402 |
| BamA/TamA family outer membrane protein | 185 | 87% | 6e-16 | 12.44 | 851 | WP_012023543.1 | Fjoh_1464 |
| ***EcTamB*** |  |  |  |  |  |  |  |
| Translocation/assembly module TamB | 166 | 88% | 7e-42 | 13.13% | 1699 | WP_012023975.1 | Fjoh_1899 |
| Translocation/assembly module TamB domain-containing protein | 152 | 85% | 9e-38 | 13.48% | 1494 | WP_012026557.1 | Fjoh_4592 |
| ***EcAsmA*** |  |  |  |  |  |  |  |
| AsmA family protein | 147 | 99% | 2e-20 | 13.84% | 867 | WP_012023792.1 | Fjoh_1716 |
| AsmA family protein | 72.4 | 84% | 7e-14 | 13.92% | 919 | WP_012023624.1 | Fjoh_1548 |
| Hypothetical lipoprotein | 115 | 81% | 4e-13 | 13.09% | 820 | WP_012023422.1 | Fjoh_1342 |
| AsmA family protein | 67.8 | 82% | 2e-12 | 13.30% | 1090 | WP_012025171.1 | Fjoh_3185 |
| AsmA family protein | 114 | 79% | 6e-12 | 10.78% | 939 | WP_012026291.1 | Fjoh_4317 |
| Translocation/assembly module TamB | 55.5 | 76% | 1e-08 | 15.73% | 1699 | WP_012023975.1 | Fjoh_1899 |

**Figure 1.**
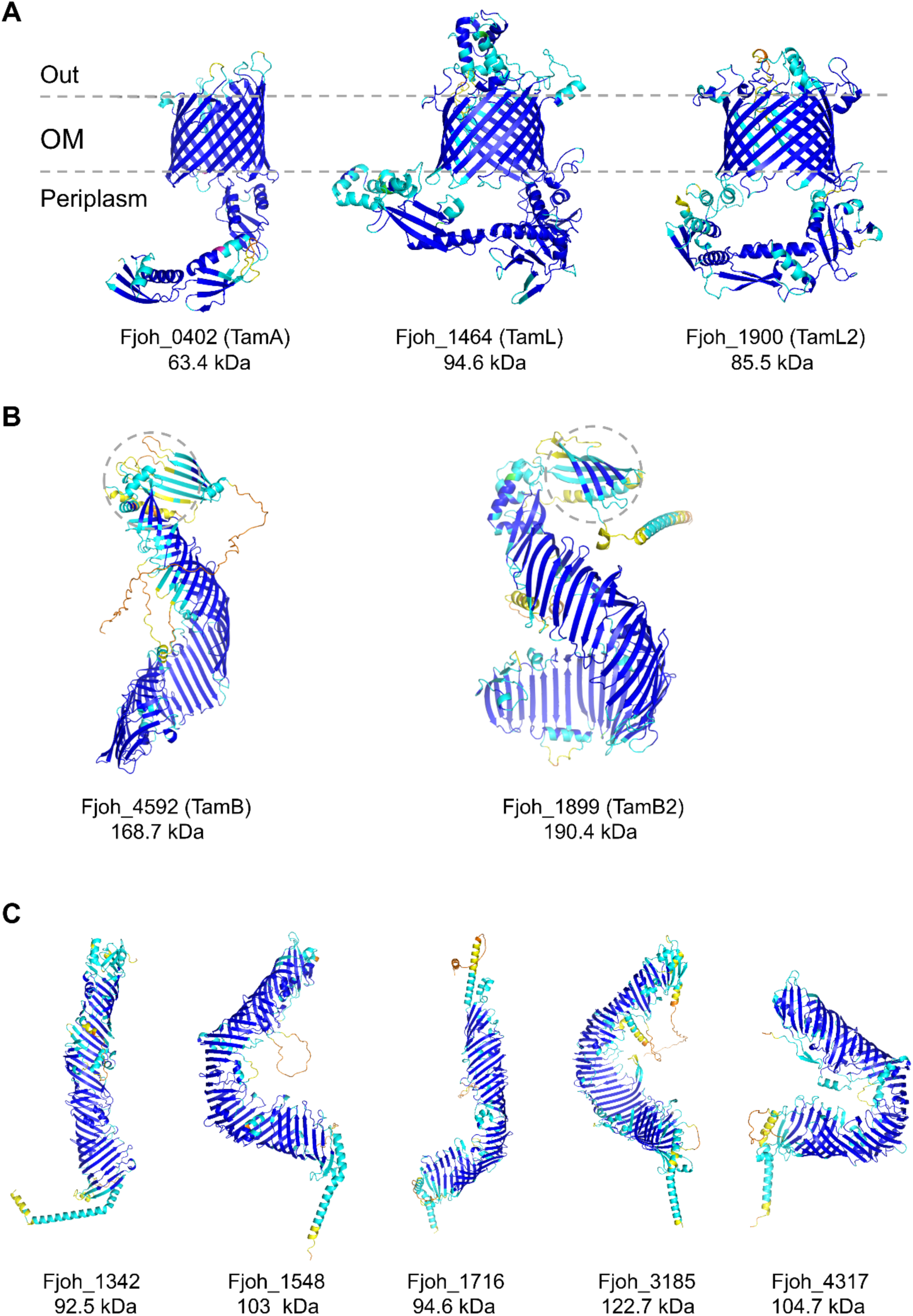
Three-dimensional structures of TamA (A), TamB (B) and AsmA-like (C) homologs in *F. johnsoniae* as predicted by AlphaFold2 (https://alphafold.ebi.ac.uk) [98]. Protein names and their predicted molecular weights are reported below each structure. Structures are colored based on the per-residue confidence score (pLDDT) between 0 and 100: dark blue (pLDDT > 90), cyan (90 > pLDDT > 70), yellow (70 > pLDDT > 50), and orange (pLDDT < 50). In B, only the last 1000 C-terminal residues are shown. The predicted C-terminal pseudosubstrate domain [10] is highlighted via a dashed circle.

We then sought putative TamB homologues using *Ec*TamB as a query. We identified two proteins encoded by *fjoh_1899* and *fjoh_4592* (Table 1). A closer inspection of their AlphaFold2-based predicted structures confirmed the presence of repeated units of the β-taco fold, typical of the AsmA-like proteins, as well as of the pseudosubstrate domain (residues 1321-1420 in Fjoh_4592; residues 1538-1643 in Fjoh_1899), a short sector of amphipathic β-barrel located at the C-terminus [17], akin to what was already observed in *Ec*TamB (Figure 1B) [10,28]. Interestingly, *fjoh_1899* may be in operon with *fjoh_1900*, likewise what already observed in several members of Bacteroidetes/Chlorobi phyla [28], since the start codon of the latter precedes the end codon of the former.

In Proteobacteria, TamB is a well-known member of the AsmA-like family, which in *E. coli* includes six members, while seven AsmA-like orthologs were recently identified in *Pseudomonas aeruginosa* [45]. As such, we sought additional AsmA-like proteins in *F. johnsoniae* using the *E. coli* AsmA protein as a query. Interestingly, in addition to *fjoh_1899*, five proteins encoded by *fjoh_1342*, *fjoh_1548*, *fjoh_1716*, *fjoh_3185* and *fjoh_4317* were retrieved (Table 1). No additional homologs could be identified using the other *E. coli* AsmA-like proteins (including *Ec*TamB) as queries.

Their AlphaFold2-based predictions strongly suggest their identity as AsmA-like proteins (Figure 1C). As a result, we concluded that *F. johnsoniae* possesses in total seven AsmA-like proteins.

### Genes *fjoh_1464* and *fjoh_4592* encode essential TamL- and TamB-homologs in *F. johnsoniae*

To investigate the role of the TamL and TamB homologs in *F. johnsoniae*, we sought to delete their encoding genes. Surprisingly, while we could delete *fjoh_0402*, *fjoh_1899* and *fjoh_1900*, as well as co-delete *fjoh_1899* and *fjoh_1900* in the wild-type background, no chromosomal deletion mutants could be obtained for *fjoh_1464* and *fjoh_4592* unless an exogenous copy was provided on a plasmid under the control of an IPTG-tunable promoter. Furthermore, when the plasmid carried a mutant of *fjoh_1464* in which the predicted site of lipidation was mutated (*fjoh_1464_21cys-gly_)*, no mutant could be obtained. These results suggest that both *fjoh_1464* and *fjoh_4592* are essential genes and that the lipidation and anchorage of the N-terminus of the protein encoded by *fjoh_1464* is probably important for its function. Accordingly, we named TamL and TamB the proteins encoded by *fjoh_1464* and *fjoh_4592*, while TamL2 and TamB2 those by *fjoh_1900* and *fjoh_1899*, respectively.

When the single mutants for *tamA*, *tamB2, tamL2,* and the double *tamB2*-*tamL2* mutant were cultured in rich (CYE) or motility medium (MM), they all grew to the same extent as the wild-type strain (Supplementary Figure S1), indicating that under standard growth conditions TamA, TamB2 and TamL2 do not have a major impact on fitness. Therefore, we decided to restrict our study to TamB and TamL owing to their essentiality.

### TamB and TamL exhibit distinct regulations within the cell

In Proteobacteria *tamA* and *tamB* are part of the same operon, suggesting that the two proteins likely undergo the same spatial-temporal regulation within the cell in order to form an active TAM complex [28,29]. Hence, we next investigated TamB and TamL regulation in *F. johnsoniae*. To this aim, we generated strains expressing tagged variants of TamL and TamB, where a tag sequence was inserted into the native genomic locus of *tamL* and *tamB*. In the case of TamL, we inserted a 3xFLAG tag, flanked by four glycine residues on both extremities, in a non-conserved extracellular loop of TamL (^574^TNQV^577^), as predicted by AlphaFold2 (Figure 2A), which in our construct was followed by the repetition of the same loop sequence. We thus generated the (3xFLAG)TamL variant (^574^TNQV-GGGG-3XFLAG-GGGG^−^TNQV^611^). As for TamB, we added a Twin-strep (2xStrep) tag at the C-terminal end, thus creating the (2xStrep)TamB variant (Figure 2B). We additionally generated a 3xFLAG-TamL/2xStrep-TamB double-tagged strain, which exhibited no significant growth difference compared to the wild-type strain when grown in CYE medium (Figure 2C), thus indicating that the two tags do not alter the function of the essential proteins TamL and TamB. We then monitored the expression of both proteins over time. Our data show that, while TamB signal was detectable throughout all time points with a slight increase during the transition from exponential to stationary phase of growth, TamL detection was limited to the initial eight hours of growth spanning from early to late-exponential phase (Figures 2C and 2D). This may indicate that TamL, but not TamB, likely undergoes strict regulation. As a result, TamL and TamB are co-present within the cell only when cells are dividing.

**Figure 2:**
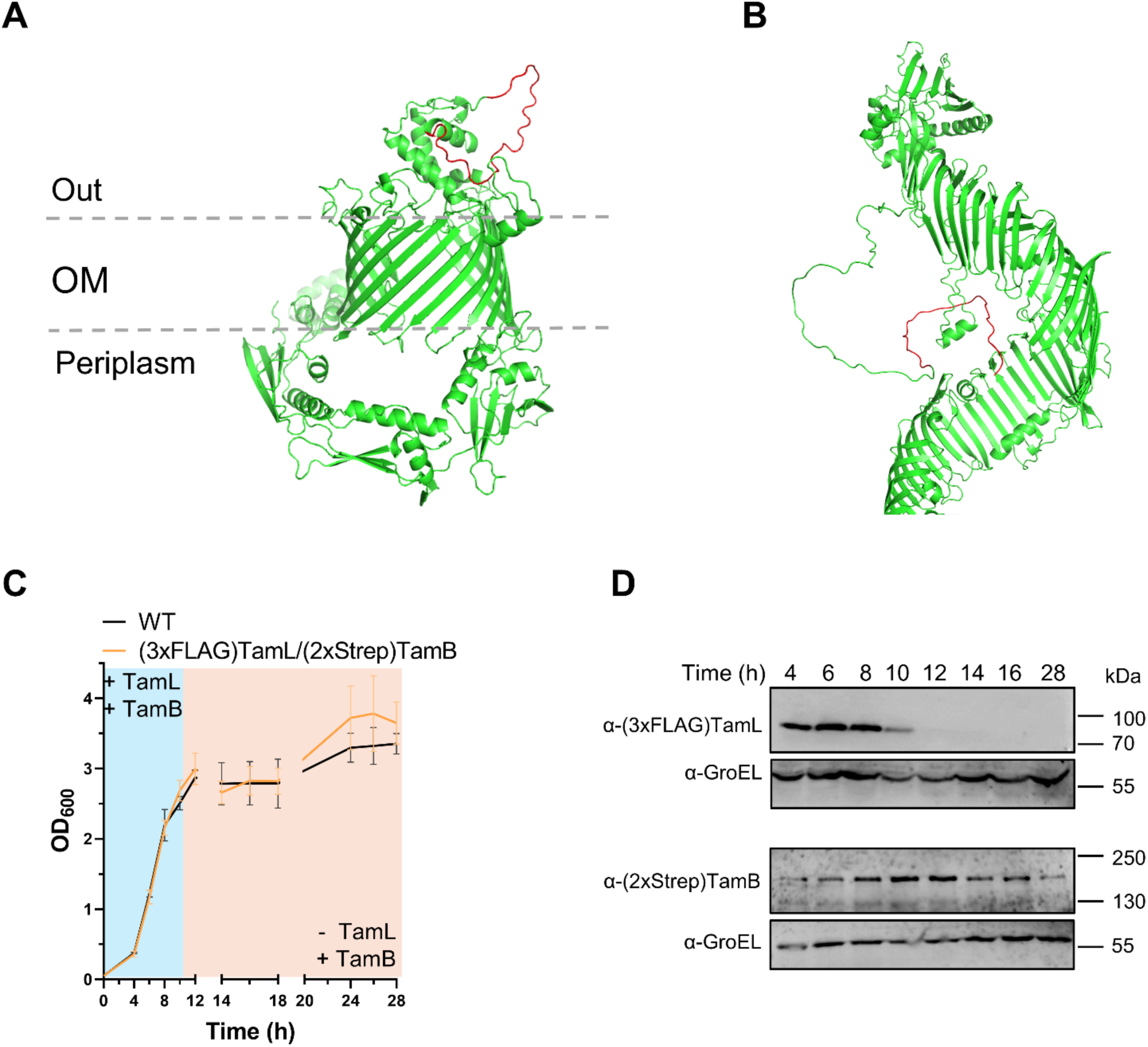
TamL and TamB expression pattern throughout cell growth. Structural models of the tagged variants of TamL **(A)** and TamB **(B)** as predicted by AlphaFold2 [96,98]. The 3xFLAG-tag sequence in TamL **(A)**, and the 2xStrep-tag sequence in TamB **(B)**, inserted downstream of the predicted extracellular loop ^574^TNQV^577^ in TamL and at the C-terminus in TamB, are highlighted in red. In **(B),** the first N-terminal 346 residues are not shown. **(C)** Growth curve in CYE medium of *F. johnsoniae* WT (black) and (3xFLAG)TamL/(2xStrep)TamB (orange) strains showing no significant effect of the two tags on cell growth. Data are displayed as mean ± standard deviation from at least three biological replicates. **(D)** TamL and TamB immunodetection throughout cell growth. GroEL detection was used as a loading control.

### TamL depletion causes loss of cell viability and leads to shape abnormalities

Given the essentiality of *tamL* and *tamB* in *F. johnsoniae*, we aimed to knock-down their expression profiles by generating TamL and TamB depletion strains. To do so, in the (3xFLAG)*tamL*/(2xStrep)*tamB* background we first generated a strain expressing the transcriptional repressor LacI under control of the *F. johnsoniae* OmpA promoter (P_ompA_::*lacI*), and then we replaced the native promoter of *tamL* with the IPTG-tunable promoter P_cfxA-lacO_, thus generating the *Fjoh* P_ompA_::*lacI*-P_cfxA-lacO_::*tamL* strain. Unfortunately, this turned out to be impossible for *tamB* as no mutant colonies could be obtained, even with the employment of other inducible promoters, such as an arabinose-inducible one. Therefore, we restricted our study exclusively to *tamL*. We cultured the *Fjoh* P_ompA_::*lacI*-P_cfxA-lacO_::*tamL* strain in CYE medium with or without IPTG (+ or - IPTG), corresponding to permissive or non-permissive conditions of growth, respectively. We noticed that, while in permissive conditions its growth curve mirrored that of the wild-type (WT) strain, in non-permissive condition its growth curve exhibited a drastic decline starting approximately at ten hours of growth (Figure 3A). A similar phenotype was previously observed in *E. coli* upon depletion of BamA [46,47]. Likewise, we observed a significant growth reduction on solid medium compared to the control strain (Figure 3B). To confirm that the observed growth defects were caused by TamL depletion, we monitored the TamL profile over time within cells grown with or without IPTG. Notably, even if under the expression of a constitutive promoter (P_cfxA-lacO_), in permissive conditions TamL-profile mirrored that of the wild-type strain (Figures 3C and 2D), which may suggest a post-translational regulation, whereas we could not detect TamL in cells grown in non-permissive conditions (Figure 3C). Furthermore, by phase-contrast microscopy we noticed that cells grown in non-permissive conditions displayed severe morphological defects unlike those grown with IPTG. Already after six hours of growth (6h), approximately 15-20% of the cells exhibited a lollipop-like shape, which were then replaced by more severe shape irregularities that became prominent over time, as indicated by the accumulation of a substantial cluster of cells and debris at later time intervals (12h and 28h, Figure 3D). These results clearly confirm that the growth of *Fjoh* P_ompA_::*lacI*-P_cfxA-lacO_::*tamL* is IPTG-dependent, and that TamL depletion is the cause of the observed phenotypes in non-permissive conditions.

**Figure 3:**
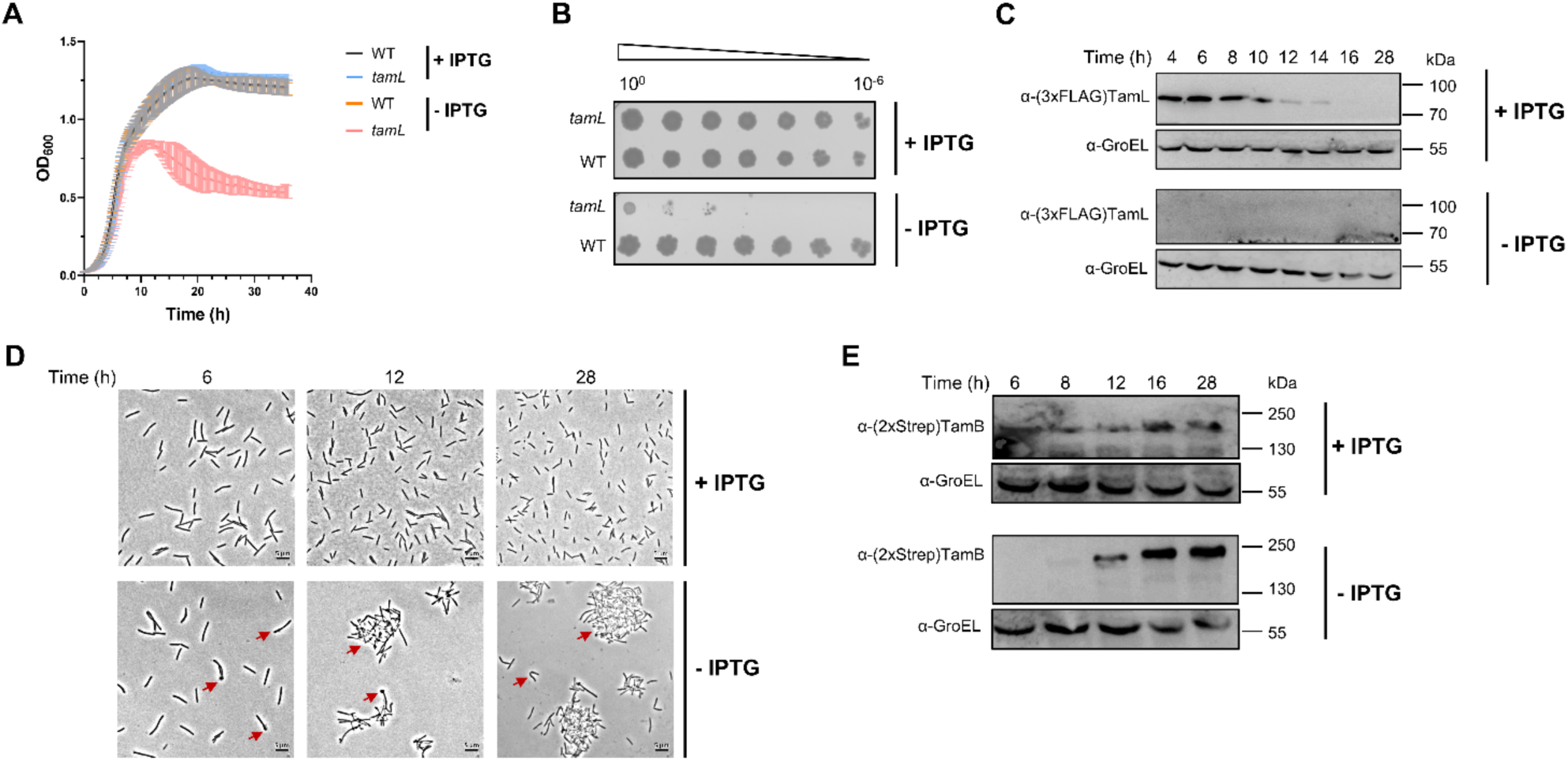
TamL depletion causes loss of cell viability and shape abnormalities. (A) Growth curves of *Fjoh* WT and *Fjoh* P_ompA_::*lacI-*P_cfxA-lacO_::*tamL* (*tamL* in figure) in CYE medium in permissive (+ IPTG) and non-permissive (- IPTG) conditions. Cells from overnight cultures (in CYE with IPTG) were OD_600_-adjusted and freshly inoculated in the same medium (+ or - IPTG) onto a 96-well plate and grown at 30°C for 36 hours. **(B) Spot assay of cells grown in permissive and non-permissive conditions.** Overnight cultures of *Fjoh* WT and *Fjoh* P_ompA_::*lacI-*P_cfxA-lacO_::*tamL* (*tamL* in figure) were adjusted for OD_600_ and then serial diluted in 1x PBS. Drops from serial dilutions were deposited onto CYE-Agar plates (+ or – IPTG). Plates were incubated at 30°C and photographed after 48 hours. **(C) Immunoblotting of (3xFLAG)TamL.** OD_600_-normalized cells were harvested at specific time points, resuspended in 1x Laemmli buffer and boiled. Proteins from total cell lysates were resolved in a SDS-polyacrylamide gel and then transferred onto a nitrocellulose membrane for (3xFLAG)TamL immunodetection. GroEL signal was used as loading control. **(D) Imaging of cells grown in permissive and non-permissive conditions.** OD_600_-normalized cells were collected at specific time points, fixed in 4% paraformaldehyde, and then visualized by phase-contrast microscopy. Red arrows indicate the observed shape abnormalities and aggregation. Scale bars represent 5 µm. **(E) Immunoblotting of (2xStrep)TamB**. Image obtained as described in (C). Data in A are displayed as mean ± standard deviation from three independent experiments. In B, C, D and E, images were from one of three independent replicates.

Interestingly, we noticed a sharp increase in TamB protein levels over time in non-permissive conditions (Figure 3E). Conversely, we found that the deletion of *tamA*, *tamL2* or both did not significantly aggravate cell fitness in TamL-depleted cells, suggesting that neither of them can substitute for TamL (Supplementary Figure S2).

### TamL depletion disrupts outer membrane integrity, with increased release of outer membrane vesicles (OMVs)

Intrigued by the morphological defects observed in TamL-depleted cells, we visualized cells grown in permissive or non-permissive conditions by transmission electron microscopy (TEM) to detect shape abnormalities at higher resolution. While in both growth conditions we could detect the presence of spherical structures resembling outer membrane vesicles (OMVs), in non-permissive conditions the OMVs appeared bigger (Figure 4A, see also Figure 6B for diameter quantification). Prompted by this finding, we sought to determine if TamL played a key role on outer membrane (OM) integrity. Therefore, we grew cells in the presence of several OM perturbing agents to assess if the depletion of TamL determined an increased susceptibility to them. We tested two antibiotics, vancomycin and polymyxin B, which target the peptidoglycan (PG) and the lipid A of lipopolysaccharide (LPS), respectively, and sodium dodecyl sulfate (SDS), a strong anionic detergent [48–50]. As shown in Figure 4B, cells incubated in the presence of the three envelope perturbing agents displayed a strong reduction in cell viability compared to the control condition (untreated cells) only when grown in non-permissive conditions, whereas a negligible effect on cell growth was observed in permissive conditions at the tested concentrations.

**Figure 4:**
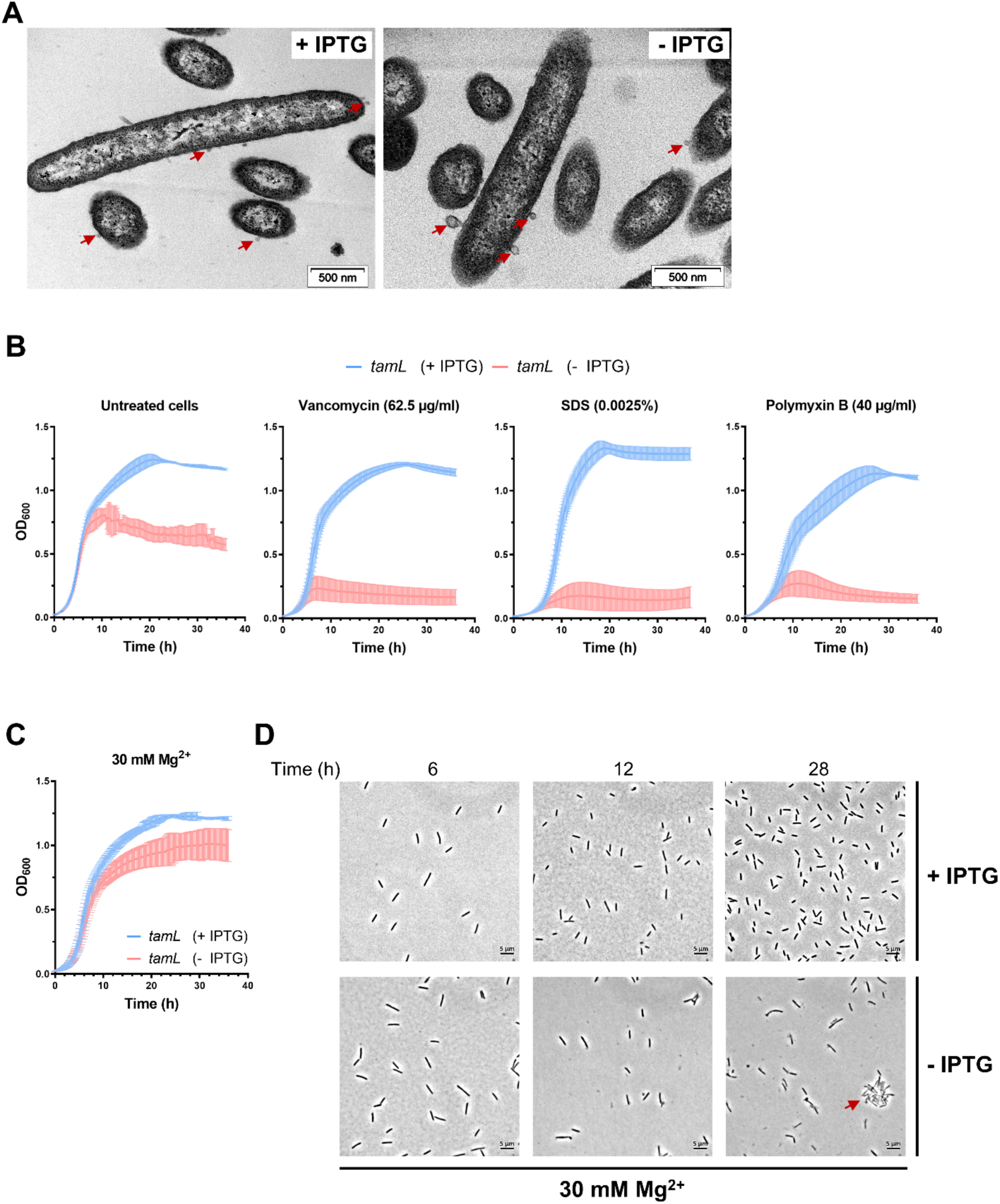
TamL depletion results in increased outer membrane permeability and release of bigger outer membrane vesicles (OMVs). (A) Transmission electron microscopy (TEM) imaging of cells grown in permissive (+ IPTG) and non-permissive conditions (- IPTG). Three biological replicates of samples from each condition were fixed onto 630 grids in triplicate. Representative OMVs are indicated by red arrows. (B and C) Growth curves of *Fjoh* P_ompA_::*lacI-* P_cfxA-lacO_::*tamL* (*tamL* in figure) in CYE medium in permissive (+ IPTG) and non-permissive (- IPTG) conditions. Cells from overnight cultures (in CYE with IPTG) were OD_600_-adjusted and freshly inoculated in the same medium in a 96-well plate and grown at 30°C for 36 hours. (B) Cells were incubated without (untreated cells) or with the envelope perturbing agents at the indicated concentrations. (C) Cells were grown in CYE medium with increased concentration of Mg^2+^. **(D) The increment in Mg^2+^ partially rescues the envelope defects in TamL-depleted cells**. Cells were grown in permissive and non-permissive conditions in CYE medium supplemented with 30 mM Mg^2+^. At specific time intervals, OD_600_-normalized cells were collected, fixed in 4% paraformaldehyde, and then visualized by phase-contrast microscopy. The red arrow at the bottom right panel indicates the observed cell aggregation. Scale bars represent 5 µm.

In *E. coli* it was shown that loss of OM material in *mlaA*\* cells (i.e., cells where the transport of phospholipid is increased from OM to the inner membrane (IM)), and the lysis observed in the Δ*yhdP*Δ*tamB* mutant could be both partially reduced by adding 1 mM MgCl_2_ in the medium [22,51,52]. In this light, when we increased the concentration of Mg^2+^ in the medium, we observed that, while having no toxic effect on cells grown in permissive conditions, the increment in Mg^2+^ in non-permissive conditions partially ameliorated cell growth (Figure 4C) and reduced but not abrogated shape abnormalities (Figure 4D). Taken together, we concluded that TamL depletion in *F. johnsoniae* is detrimental for cell viability and is accompanied by severe membrane and permeability defects, suggesting that TamL is required for OM integrity and homeostasis.

### TamL depletion alters outer membrane (lipo)protein composition

The above results clearly indicate the key function of TamL in maintaining OM integrity. Hence, we wondered if the observed phenotypes could be attributed to an alteration of the OM proteome caused by TamL depletion. Therefore, we isolated the membrane fractions from cells grown in permissive and non-permissive conditions. While the total protein content in each fraction was similar between conditions (Supplementary Figure S3), SDS-PAGE analysis indicated a clear difference in protein composition (Figure 5A).

**Figure 5:**
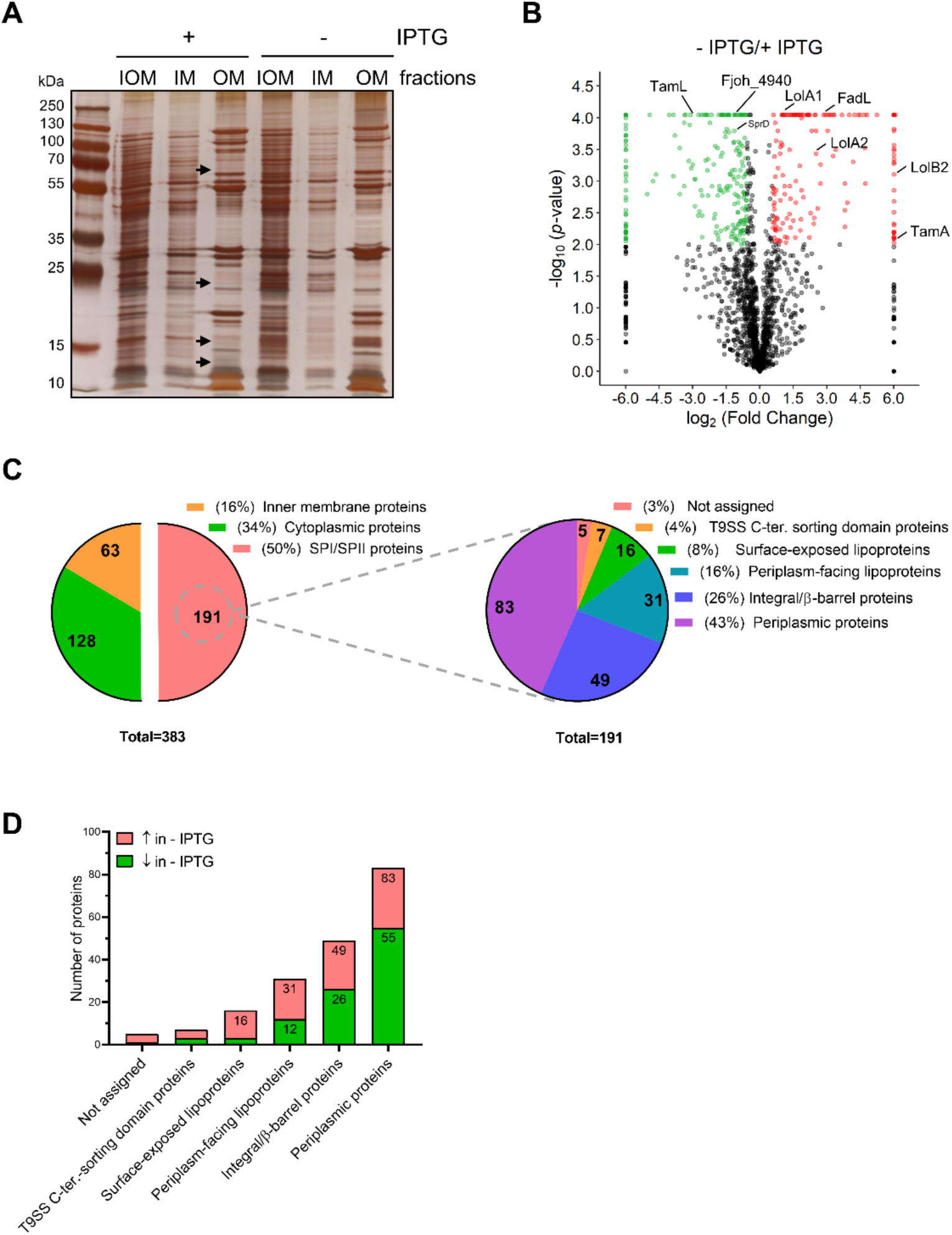
TamL depletion significantly affects the outer membrane protein composition. **(A)** AgNO_3_–stained polyacrylamide gel (12%) of the membrane fractions isolated from cells grown in permissive (+ IPTG) or non-permissive (- IPTG) conditions. Prior to gel electrophoresis, the total protein content of each fraction was quantified by Bradford protein assay (Bio-Rad) following the manufacturès instructions, and then 2 µg of each fraction was used. Black arrows indicate protein bands of the OM fractions showing different intensities in the two conditions (+/− IPTG). **(B)** Volcano plot depicting the fold change (FC) and the statistical significance (*p*-value) of the proteome of the outer membrane fractions from cells grown with/without (+/−) IPTG. The FC corresponds to - IPTG/+ IPTG ratio. Green and red dots indicate significantly decreased or increased proteins (FC ≥ |1.5|; p < 0.01), respectively, in cells grown – IPTG compared to those grown + IPTG. Black dots represent those proteins either not significantly affected (FC < |1.5|) or with a *p-*value ≥ 0.01. When a protein was not identified in one of the two experimental conditions, a representative FC value of |64| was used, while for those proteins whose *p*-value was < 0.0001, a representative value of 0.00009 was used. The projection of some proteins described in the main text is shown. **(C)** Left: pie chart showing the number of inner membrane proteins, cytoplasmic proteins and proteins with a signal peptide (SPI/SPII) and their relative abundance (%) over the total number of significantly affected proteins (i.e., 383). Right: same as left but depicting the number and the relative abundance of the different classes of SPI/SPII proteins over the total number of significantly affected SPI/SPII proteins (i.e., 191). **(D)** Bar chart showing the number of different classes of SPI/SPII proteins found significantly increased (red) or decreased (green) upon TamL depletion (- IPTG).

Subsequent mass spectrometry (MS) analysis identified a total of 1834 and 1716 proteins in the OM fractions from permissive and non-permissive conditions, respectively, 1565 of which were shared by both conditions (Supplementary Table S1). Upon sorting the identified proteins according to their cellular localization, we found a total of 383 proteins to be significantly affected (FC ≥ |1.5|; p < 0.01) (Figure 5B), of which half (191) were predicted to have a signal peptide (SPI or SPII). 91 proteins out of 191 (48%) were increased, while the remaining 100 (52%) were decreased upon TamL depletion (Supplementary Table S1). Amongst the 191 proteins with either SPI or SPII, the periplasmic proteins were the most abundant (83), followed by integral/β-barrel proteins (49) (i.e., proteins anchored to the OM as part of a complex or embedded into the OM due to the β-barrel domain), OM lipoproteins (47), and proteins annotated or exhibiting a Type 9 secretion system (T9SS) C-terminal sorting domain (7) [1,39,40,53]. For only 5 proteins, the proper localization could not be assigned. Notably, 16 out of the 47 OM lipoproteins were predicted to be surface-exposed (i.e., anchored on the outer leaflet of the OM), owing to the presence of the lipoprotein export signal (LES) typical of members of the Bacteroidetes phylum [32]; in contrast, the remaining 31 were predicted to be anchored to the inner leaflet, and thereby facing the periplasm.

A closer inspection of the impacted protein classes revealed significant changes in integral/β-barrel proteins (Supplementary Table S1). Among those increased, three were predicted to be TonB-dependent receptors involved in substrate uptake [54], while nine were predicted to be implicated in cell wall/membrane/envelope biogenesis. Of these, three were annotated as Outer Membrane Efflux Proteins (OEPs), suggesting their involvement in the Type 1 Secretion System (T1SS) [55]. Additionally, we identified Fjoh_4941 (FC: 5.6). Based on the AlphaFold2-predicted structure, the PFAM annotation and the sequence alignment by DELTA-BLAST (E-value: 8e-85), Fjoh_4941 is a potential homologue of the long-chain fatty acid (LCFA) transport protein FadL, which in *E. coli* plays a crucial role in maintaining OM integrity by importing LCFAs for membrane lipid synthesis [56]. Strikingly, we noticed that the putative TamA in *F. johnsoniae* (Fjoh_0402) stood out as one of the most increased proteins (Supplementary Table S1). In agreement with the depletion experiments, TamL (Fjoh_1464) was instead one of the most decreased proteins (FC: 0.1). Moreover, TamL depletion significantly decreased several TonB-dependent receptors (eleven in total), eight of which are encoded by genes that are likely part of polysaccharide utilization loci (PULs) that in Bacteroidetes play a fundamental role for the assimilation and metabolism of complex sugars [57]. Additionally, the SprD (Fjoh_0980), part of the T9SS, was significantly downregulated (FC: 0.4) upon TamL depletion [38,53].

Regarding the impact of TamL depletion on OM lipoproteins, we observed an increase in most of these proteins (31 out of 47), with surface-exposed lipoproteins being particularly affected (13 out of 16 displayed increased abundance) (Supplementary Table S1), though the function of most of these increased lipoproteins remains unclear.

Among the OM lipoproteins lacking LES, the LolB homologue Fjoh_1084 (LolB2) [58] showed a dramatic increase upon TamL depletion (FC: INF). Additionally, the two putative LolA homologs Fjoh_2111 (LolA1) and Fjoh_1085 (LolA2) were also elevated, suggesting a potential upregulation of the Lol system(s) in response to TamL depletion [58,59]. We also observed an increased abundance of the lipoprotein Fjoh_4940 (FC: 2.6) which, given the genomic proximity of its encoding gene (*fjoh_4940*) with that of the putative FadL protein (*fjoh_4941*), might indicate a functional link.

Not much information could be inferred from the OM lipoproteins decreased upon TamL depletion (16 out of 47).

Overall, from the above proteomic analysis we concluded that the depletion of TamL, while broadly affecting the OM protein content, significantly increases OM lipoprotein content.

### TamL depletion causes the release of bigger outer membrane vesicles (OMVs) with an altered protein content

As just described, TamL depletion is accompanied by the shedding of bigger OMVs. To better understand TamL role in membrane homeostasis, we then investigated whether and how OMV protein composition was affected by TamL depletion by purifying OMVs from cells grown in permissive and non-permissive conditions. We noticed that, at equal cell density (OD_600_) and growth volume, the dry weight of the OMVs isolated from TamL-depleted cells was ten times higher (60 vs 4-6 mg/L), and these OMVS exhibited an approximately twofold increase in diameter (median: 185 vs 87 nm, Figures 6A and 6B), as well as an approximative eightfold increase in the protein content (0.429 ± 0.030 vs 0.055 ± 0.020 mg, Figure 6B). The qualitative analysis of the OMVs proteome by AgNO_3_ staining revealed an even more noticeable difference in the protein band pattern between the OMVs isolated from the two conditions of growth compared to the that observed in the OM fractions (Figure 6C), which clearly indicates that TamL depletion has a strong impact on the OMVs protein composition. To further explore this, we then analyzed by MS the proteome of the OMVs isolated from cells grown in permissive and non-permissive conditions. A total of 1518 (+ IPTG) and 1339 (- IPTG) proteins were identified (Supplementary Table S2), 1105 of which were shared. 355 of these identified proteins were significantly affected (FC ≥ |1.5|; p < 0.01), 217 of which (61%) predicted to have a signal peptide (SPI or SPII) (Figures 6D and 6E).

**Figure 6:**
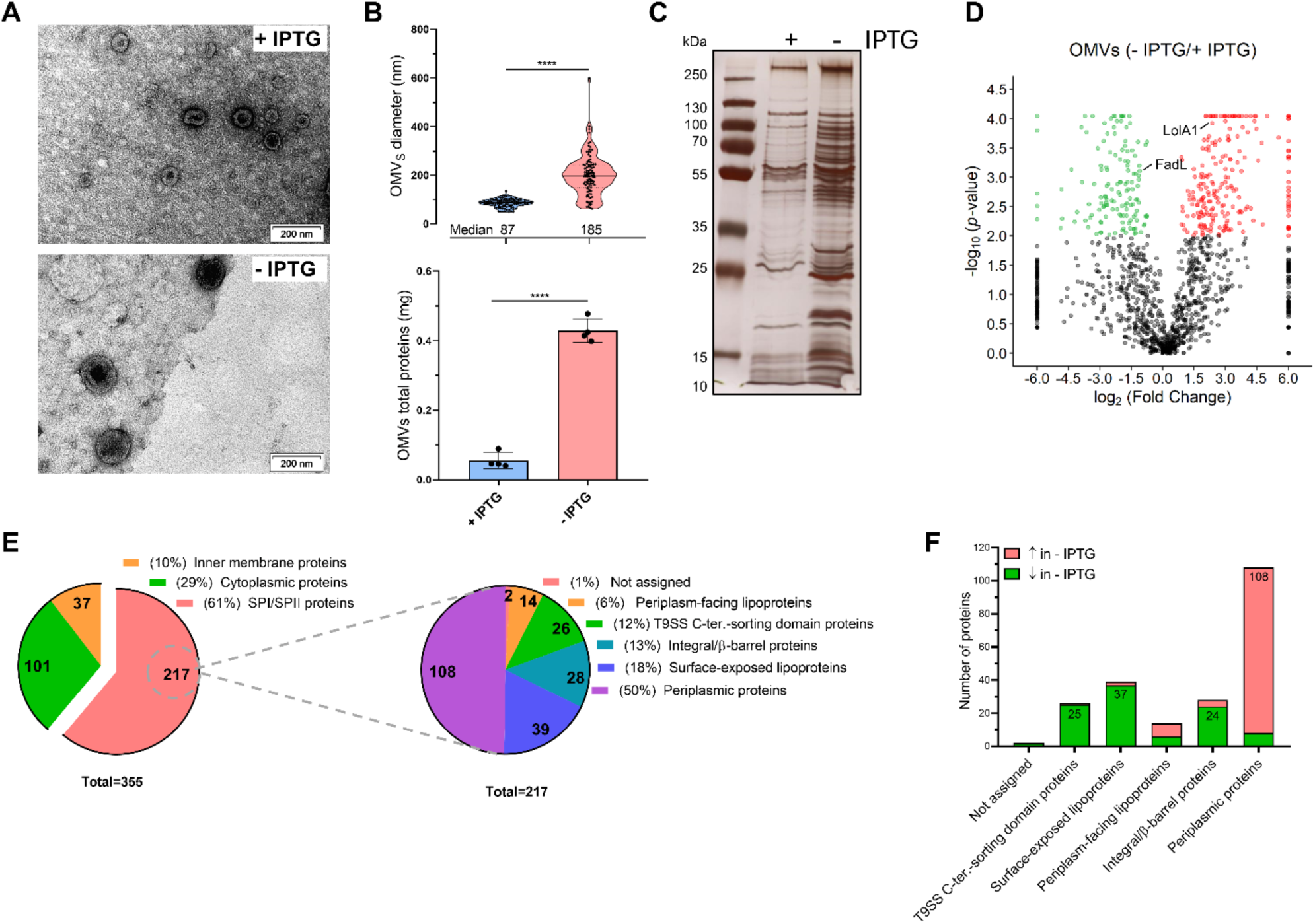
The OMVs shed from TamL-depleted cells are larger and enriched with periplasmic proteins. **(A)** Representative TEM images of OMVs isolated from cells grown in permissive (+ IPTG) and non-permissive (- IPTG) conditions. OMVs purified from biological replicates of each growth condition were fixed and stained as described in materials and methods before TEM visualization. **(B)** Top: violin plots quantifying the OMVs diameter (in nm). From each TEM image, the diameter of the OMVs (*n*=102 per condition) was measured by ImageJ using the scale bar as a reference. In the plots, the horizontal black line indicates the median, while each dot indicates each single value. Bottom: the total protein content of the OMVs was quantified using the Pierce™ BCA Protein Assay kit (ThermoFisher Scientific) following the manufacturer’s instructions. Data are displayed as mean ± standard deviation from four independent samples per condition. In both cases, a two-tailed *t*-test was performed for statistical analysis. Statistical significance is given as: ****(*p* < 0.001). **(C)** AgNO_3_–stained polyacrylamide gel (12%) of the OMVs isolated from cells grown in permissive (+ IPTG) or non-permissive (- IPTG) conditions. Prior to gel electrophoresis, the total protein content was quantified as described above. 2 µg was used. **(D)** Volcano plot depicting the fold change (FC) and the statistical significance (*p*-value) of the proteome of the OMVs collected from cells grown with/without (+/−) IPTG. The FC corresponds to - IPTG/+ IPTG ratio. Green and red dots indicate those proteins found significantly decreased or increased (FC ≥ |1.5|; p < 0.01), respectively, in the OMVs from cells grown – IPTG compared to the OMVs from cells grown + IPTG. Black dots represent those proteins either not significantly affected (FC < |1.5|) or with a *p-*value ≥ 0.01. When a protein was not identified in one of the two experimental conditions, a representative FC value of |64| was used, while for those proteins whose *p*-value was < 0.0001, a representative value of 0.00009 was used. The projection of some proteins described in the main text is shown. **(E)** Left: pie chart showing the number of inner membrane proteins, cytoplasmic proteins and proteins with a signal peptide (SPI/SPII) and their relative abundance (%) over the total number of significantly affected proteins (i.e., 355). Right: same as left but depicting the number and the relative abundance of the different classes of SPI/SPII proteins over the total number of significantly affected SPI/SPII proteins (i.e., 217). **(F)** Bar chart showing the number of different classes of SPI/SPII proteins found significantly increased (red) or decreased (green) in the OMVs upon TamL depletion (- IPTG).

A deeper inspection of the 217 SPI or SPII proteins showed that the predicted periplasmic proteins represented the most numerous class (108 out of 355), followed by OM lipoproteins (54 out of 355), 39 and 14 of which predicted to be surface-exposed or periplasm-facing lipoproteins, respectively; integral/β-barrel OMPs (28 out of 355), and proteins exhibiting a T9SS C-terminal sorting domain (26 out of 355). For only 2 proteins we could not assign a precise localization (Figure 6E).

Interestingly, we observed that the OMVs isolated in permissive conditions were almost exclusively enriched with cytoplasmic and periplasmic proteins. In particular, we observed that the vast majority (100 out of 108) of the predicted periplasmic proteins showed a significant increment upon TamL depletion (Figure 6F). Most of them were peptidases and glycosidases (in the latter case belonging to PULs), involved in amino acid and carbohydrate transport and metabolism, or in cell wall/membrane/envelope biogenesis. Notably, the periplasmic proteins also stood out as the category of SP proteins with the greatest fold change (FC), with approximately half (48 out of 100) exhibiting FC ≥ 10. Furthermore, the LolA homologue (Fjoh_2111) [58,59] was found to increase (FC: 3.9, Figure 6D), alike what was observed in the OM proteomics analysis.

In high contrast, the OMVs released upon TamL depletion were nearly completely devoid of integral/β-barrel OMPs, surface-exposed lipoproteins, and proteins with a T9SS C-terminal sorting domain (Figure 6F). Above all, surface-exposed lipoproteins turned out to be the SP proteins most negatively impacted, with 37 decreased proteins out of 39 (Figure 6F). Interestingly, although their precise role within the cell is unknown, we noticed that the majority of them was predicted to have one or more domains involved in carbohydrate/lipid/small molecule-protein binding, such as the SusD/RagB-, SusE-, PKD- or lipocaline-like domains, which probably work in concert with integral/β-barrel OMPs that participate in the transport and metabolism of these ligands. In fact, while up to 24 out of the 28 identified integral/β-barrel OMPs were strongly decreased in the OMVs upon TamL depletion (Figure 6F), more than half of them are annotated as TonB-dependent receptors. Furthermore, we once again identified the putative homologue of FadL (Fjoh_4941; FC: 0.4), but not the lipoprotein Fjoh_4940 previously observed to be enriched in our OM fractions upon TamL depletion (Figure 6D).

As for the proteins with a predicted T9SS C-terminal sorting domain, alike the surface-exposed lipoproteins, this class of SP proteins stood out as one of the most negatively affected by TamL depletion (25 out of 26 proteins strongly decreased) (Figure 6F). We noticed that most of them belonged to PULs and displayed one or more carbohydrate-binding modules (CBMs) and glycosyl hydrolase (GH) domains. This, combined with the predicted association of their encoding genes with PULs, suggests their involvement in carbohydrate transport and metabolism.

Taken together, our results reveal that, under physiological conditions of growth, *F. johnsoniae* produces and releases OMVs that are enriched of integral/β-barrel OMPs, OM lipoproteins, and of proteins predicted to pass through the T9SS, primarily involved in the transport and metabolism of complex sugars and metabolites, or in the homeostasis of cell-wall. In contrast, the loss of OM integrity due to TamL depletion is associated with the abnormal release of larger OMVs whose protein cargo is primarily composed of periplasmic proteins.

### TamL depletion does not alter the OMVs lipid composition

Lipidomic studies have shown that *F. johnsoniae* outer membrane (OM) is mainly composed of sulfonolipids (SLs), ornithine lipids (OLs) and the phospholipid phosphatidylethanolamine (PE), while other minor lipids can be detected depending on the growth conditions [33,34]. SLs and OLs represent the two dominant lipid classes of *F. johnsoniae* OM, and studies performed on a mutant deficient in *fjoh_2419*, encoding an 8-amino-7-oxononanoate synthase needed for SLs biosynthesis, suggest that the balance between SLs and OLs is crucial for the maintenance of the OM stability [34]. In the *E. coli* mutant lacking YhdP and TamB, the decrease in the phospholipid anterograde (IM to OM) transport causes an accumulation of LPS in the OM, which then leads to increased OMVs production and release to maintain the proper LPS/phospholipid balance [22,23]. As our data show that TamL depletion causes blebbing and severely compromises the OM integrity, we wondered if this phenotype was a consequence of lipid asymmetry in the OM. Hence, we first extracted the lipid content of the membrane fractions and OMVs isolated from cells grown in permissive and non-permissive conditions, and we then analyzed their lipid profile by thin layer chromatography (TLC). While in the membrane fractions isolated in the two conditions (+/− IPTG) we did not observe main changes in the OM lipid pattern (Supplementary Figure S4), we instead detected more PE in the OMVs from TamL-depleted cells, whereas no substantial changes in the level of OLs could be observed (Figure 7A). The SLs content could not be assessed. We then analyzed the OMVs lipid profile by LC-MS to gain more insights. Although our MS analysis did not provide an absolute quantification, we found that the PE/OLs ion intensities were higher in the OMVs from TamL-depleted cells, in support of the TLC results (Figure 7B). Additionally, the analysis detected elevated levels of SLs in both sample types (i.e., OMVs from cells grown with +/− IPTG), though no difference was observed. Therefore, we could conclude that TamL depletion affects the PE/OLs balance in the OMVs, likely determining an increase in the PE levels, whereas no major effects are seen on OLs and SLs.

**Figure 7.**
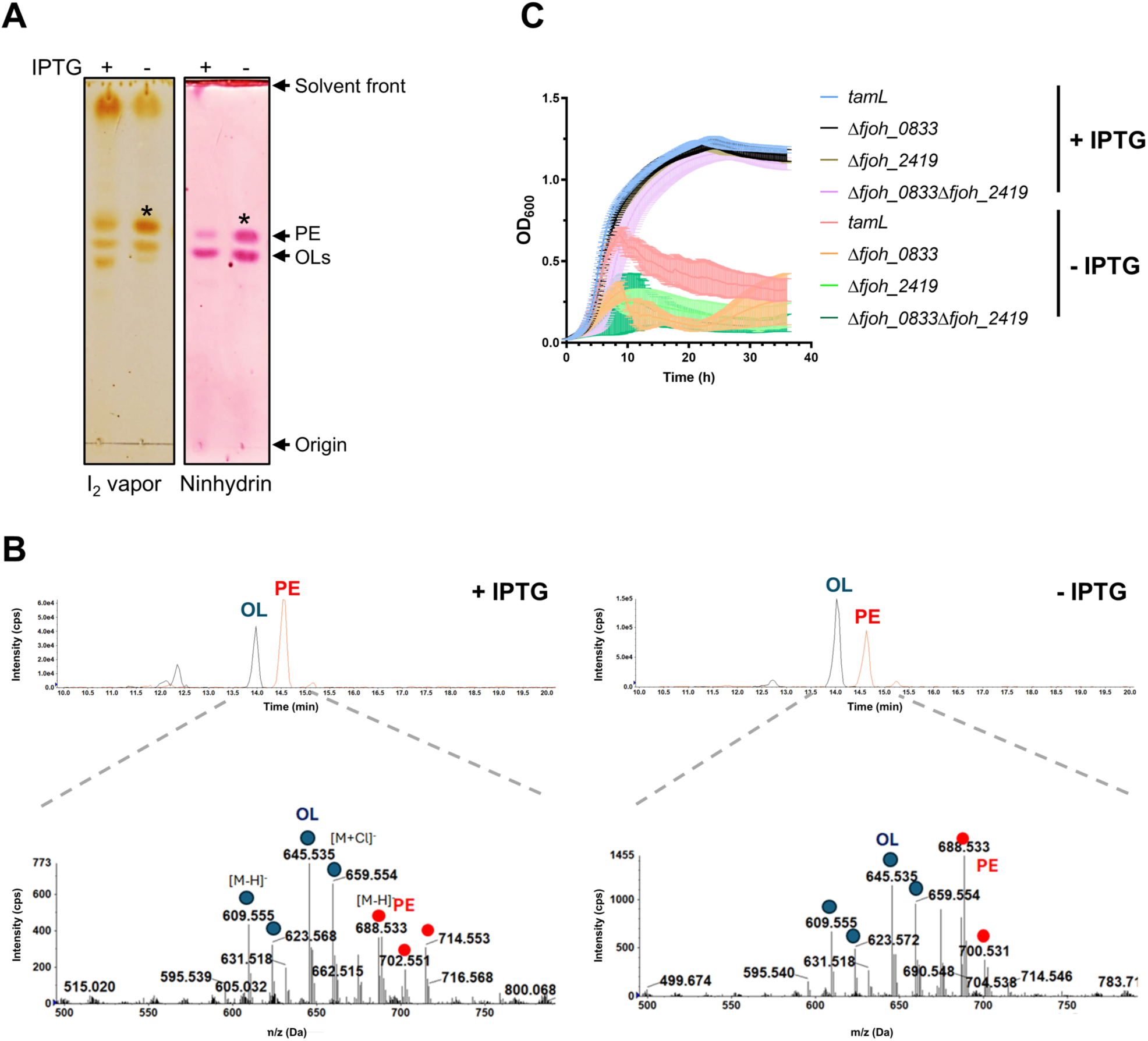
Depletion of TamL affects the lipid content of OMVs and the growth of *F. johnsoniae* mutants deficient in sulfonolipid and ornithine lipid formation. Lipids from OMVs isolated from cells grown in permissive (+ IPTG) or non-permissive conditions (- IPTG) using a protocol adapted from [90], and then analyzed by TLC (A) or LC-MS (B). **(A)** Lipids were resolved in a solvent mixture of chloroform/methanol/ammonium hydroxide with a ratio of 140:60:10 before being revealed by iodine (I_2_) vapor and ninhydrin staining. A significative phosphatidylethanolamine (PE) increment in the OMVs from – IPTG is observed (labelled * and quantified by ImageJ from three biological replicates). PE and ornithine lipids (OLs) were assigned to the respective spots based on standards. **(B)** Upper panels: ion chromatograms of the LC-MS analysis of total lipids extracted showing the elution profiles of PE and OLs. Bottom panels: negative ion mode mass spectra of the fractions of the upper panels with retention times between 13.5 and 15 min where PE and OLs were expected to be eluted off the column. Both lipid species were subjected to MS/MS analysis, and the fragmentation confirmed their identity. **(C)** Growth curves of the *F. johnsoniae* TamL depletion strain, *Fjoh* P_ompA_::*lacI-* P_cfxA-lacO_::*tamL* (*tamL* in figure), and of the same lipid mutants as in C but in the TamL-depletion parental strain in permissive (+ IPTG) and non-permissive conditions (- IPTG). In C and D, cells from overnight cultures (in CYE with IPTG) were OD_600_-adjusted and freshly inoculated in the same medium in a 96-well plate and grown at 30°C for 36 hours. Data are shown as mean ± standard deviations from at least three biological replicates.

As stated above, SLs and OLs represent the two most abundant lipid classes in the OM of *F. johnsoniae*, and a balance between these two classes exists to maintain the OM integrity. If TamL is the SLs and/or OLs transporter, mutants unable to synthesize either or both lipids would result in the loss of cell viability to same extent as that observed upon TamL depletion. To prove this, we generated single and double mutants deficient in *fjoh_*2419 (described above) and *fjoh_0833* (encoding a bifunctional acyltransferase responsible for OLs) [34,60]. To our surprise, despite the importance SLs and OLs have on OM lipid composition, both the single and double mutants (Δ*fjoh_0833*Δ*fjoh_2419*) could be generated. We first tested the growth performance of these mutants in rich (CYE) and motility (MM) media and we compared them to the wild-type strain grown in the same conditions. Interestingly, in CYE medium no substantial differences could be observed, which indicates that the co-deletion of *fjoh_0833* and *fjoh_2419* has no effect on cell fitness and does not replicate the loss of cell viability observed upon TamL depletion (Supplementary Figure S5). In contrast, in MM only the double mutant Δ*fjoh_0833*Δ*fjoh_2419* displayed a significative growth reduction, which may be attributed to the fact that in this strain no other major lipid classes can fully complement the lack of both OLs and SLs (Supplementary Figure S5).

We then performed the same experiment using the same mutants for *fjoh_*0833 and *fjoh_2419* but in the TamL depletion strain. If TamL is the OLs/SLs transporter, *fjoh_*0833 and/or *fjoh_2419* are in epistatic interaction with *tamL*, that is, their deletion should not worsen the reduced growth fitness observed upon TamL depletion. In contrast, we observed that both single mutants for *fjoh_0833* and *fjoh_2419* exhibited a drastic growth reduction upon TamL depletion compared to the control strain, and this phenotype was further exacerbated in the double mutant (Δ*fjoh_0833*Δ*fjoh_2419*) (Figure 7C). These findings clearly contradict the hypothesis that TamL may serve as OLs/SLs transporter. Therefore, based on these results and the lipidomic analyses of membrane fractions and OMVs, we conclude that TamL is most likely not implicated in the transport of the main OM lipid classes in *F. johnsoniae*.

### TamL physically interacts with TamB

It is known that in Proteobacteria the N-terminal POTRA domains of TamA contact the C-terminal end of TamB to form an active TAM complex [13,14]. This implies that TamA and TamB become functional within the cell only when they establish physical interaction. Since our data do not support the hypothesis that TamL is implicated in the anterograde transport of OM lipids, we then wondered if TamL truly establishes physical interaction with TamB *in vivo*. To prove this, we performed pull-down experiments after crosslinking with DTSSP using (3xFLAG)TamL as a bait. Although we could not detect any TamB signal in the TamL-elution fractions by immunoblot, likely due to the very low abundance of this protein within the cell as shown in Figure 2D, MS data analyses confirmed the presence of TamB only in the (3xFLAG)TamL-elutions, which stood out as the most abundant protein after TamL (Table 2). Our pull-downs assays did not allow us to identify with statistical significance OMPs that co-purified exclusively with TamL. At the current state, we could not rule out the possibility that TamL, upon interaction with TamB, may transport OMPs.

**Table 2.** TamL pull-down results. Data expressed as Total Spectra count from two biological replicates are shown. Statistical significance (*p*-value) was calculated via Fisher’s exact test comparing the data from mock (TamB single-tagged strain) and TamL/TamB double-tagged strain in each replicate.

| Identified protein | Accession ID | Molecular Weight | Replicate | *p*-value | Mock | Double-tagged strain |
| --- | --- | --- | --- | --- | --- | --- |
| TamL-3xFLAG | Fjoh_1464 | 98 kDa | 1 | < 0.00010 | 0 | 341 |
|  |  |  | 2 | < 0.00010 | 0 | 188 |
| 2xStrep-TamB | Fjoh_4592 | 169 kDa | 1 | < 0.00010 | 0 | 77 |
|  |  |  | 2 | < 0.00010 | 0 | 35 |

Once established that TamL and TamB interact *in vivo*, we then sought to visualize the putative interaction occurring between these two proteins using AlphaFold Multimer [61]. Interestingly, the C-terminal pseudosubstrate domain of TamB (residues 1321-1420) was predicted to interact via β-strand augmentation with the last β-strand of the β-barrel domain of TamL in a manner similar to what has been proposed to occur between TamA and TamB [26], and also between BamA and its outer membrane protein substrates [9,62–65]. Furthermore, a similar interaction has been recently proposed to occur between the C-terminus of YdbH, an AsmA-like protein, and the OM lipoprotein YnbE [27]. As a result of this interaction, the TamL β-barrel is wider, being now made of 22 β-strands, and internally crossed by a small fragment of TamB C-terminus (Supplementary Figure S6A).

### The monocistronic *tamL* and *tamB* genes are widely conserved in Bacteroidetes, whereas *tamA* and the *tamB-tamL* operon are not

Previous phylogenetic analyses have shown that TamB sequences can be found in all diderm bacteria, while TamA or TamL sequences are solely restricted to Proteobacteria and Bacteroidetes/Chlorobi, respectively, where they occur in an operon with TamB [28]. The surprising fact that *F. johnsoniae*, a member of the Bacteroidetes phylum, possesses multiple TamL- and TamB-encoding genes, as well as a gene coding for a putative TamA, led us to investigate whether this genetic feature could be attributed to other Bacteroidetes members. To this aim, we performed an *in silico* analysis searching for TamA, -L, and -B homologs in several Bacteroidetes species using the protein sequences of the homologs from *F. johnsoniae* as a query (Figure 8, and Supplementary Table S3). Strikingly, we found that all the species investigated possess putative TamL and TamB proteins, whose genes are not in operons, that display high sequence similarity with the essential TamL and TamB proteins of *F. johnsoniae*. In 20 out of 30 of these species, we also identified the TamB and TamL proteins, encoded by the operon *tamB-tamL*, exhibiting higher sequence similarity with the not essential proteins TamB2 and TamL2 of *F. johnsoniae*. Surprisingly, amongst those species lacking the operon, we found *F. columnare* and *F. psychrophilum* of the *Flavobacterium* genus, and all members of the *Capnocytophaga* genus (Figure 8).

**Figure 8.**
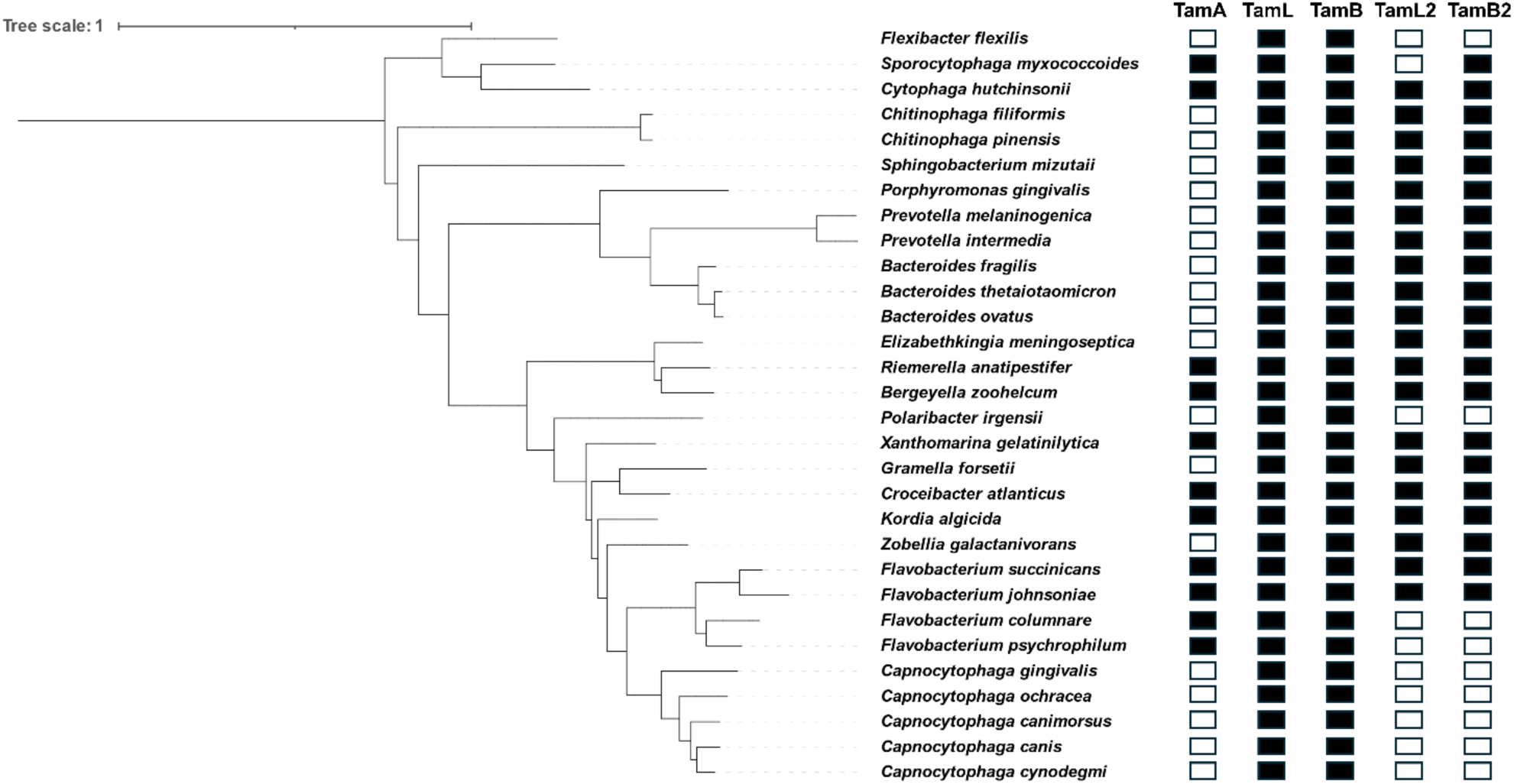
Conservation of TAM proteins in Bacteroidetes. The sequences of TamA, TamL, TamB, TamL2 and TamB2 from *F. johnsoniae* were used to identify their respective homologs in Bacteroidetes by DELTA-BLAST [43]. A phylogenetic tree was then generated via phyloT [99] based on NCBI taxonomy. Black and white squares indicate that the presence or absence of the protein homolog, respectively.

Concerning TamA, our analysis identified putative TamA proteins in eleven species (Figure 8 and Supplementary Table S3). We consider this finding remarkable as it revealed the presence of TamA in other Bacteroidetes species in addition to *F. johnsoniae*, thus confirming that TamA is not only confined to Proteobacteria but is rather more widespread contrary to prior assumptions [28]. In summary, our *in silico* analysis revealed that unlike in Proteobacteria, where the genes coding for the proteins TamA and TamB are found together in an operon [28], the TamL and TamB encoding genes do not consistently co-occur in an operon across all Bacteroidetes. Instead, a distinct characteristic of Bacteroidetes appears to be the presence of monocistronic TamL and TamB encoding genes. Moreover, some Bacteroidetes species seem to possess a putative TamA protein, thus suggesting the possible co-presence of TamA and TamL in the outer membrane of these species, as we observed in *F. johnsoniae*.

### TamL essentiality and its interaction with TamB is not solely restricted to *F. johnsoniae*

As pointed out by our *in silico* analysis, *C. canimorsus*, a human pathogen and a commensal bacterium of cat and dog mouth whose genus is closely related to that of *Flavobaterium* [66], possesses only one putative TamL homologue (Protein accession: WP_042002088.1; Supplementary Table S3) and only one TamB homologue (Protein accession: WP_095900346.1; Supplementary Table S3) encoded by genes *Ccan_17810* and *Ccan_13100*, respectively. Through the inspection of their AlphaFold2-based predicted structures, we could confirm their structural identity as TamL and TamB homologues (Figures 9A and 9B), where cys20 is the predicted site of lipidation (likelihood >0.99) for TamL. Prompted by these observations, we wondered if TamL and TamB were essential proteins also in this species. Similarly to the TamL and TamB homologs in *F. johnsoniae*, the deletion of *Ccan_17810* and *Ccan_13100* was possible only using a wild-type strain (Cc5) previously transformed with a plasmid carrying either *Ccan_17810* or *Ccan_13100* under an IPTG-tunable promoter (Supplementary Table S4). Furthermore, in the case of *Ccan_17810*, this could not be achieved in the presence of a mutation on the predicted site of lipidation (*Ccan_17810 _cys21gly_*). We therefore conclude that also in *C. canimorsus* the TamL and TamB homologs are essential proteins and that the lipidation of TamL is crucial. Notably, TamB was the only AsmA-like protein that we could identify in *C. canimorsus* in contrast to *F. johnsoniae*.

**Figure 9.**
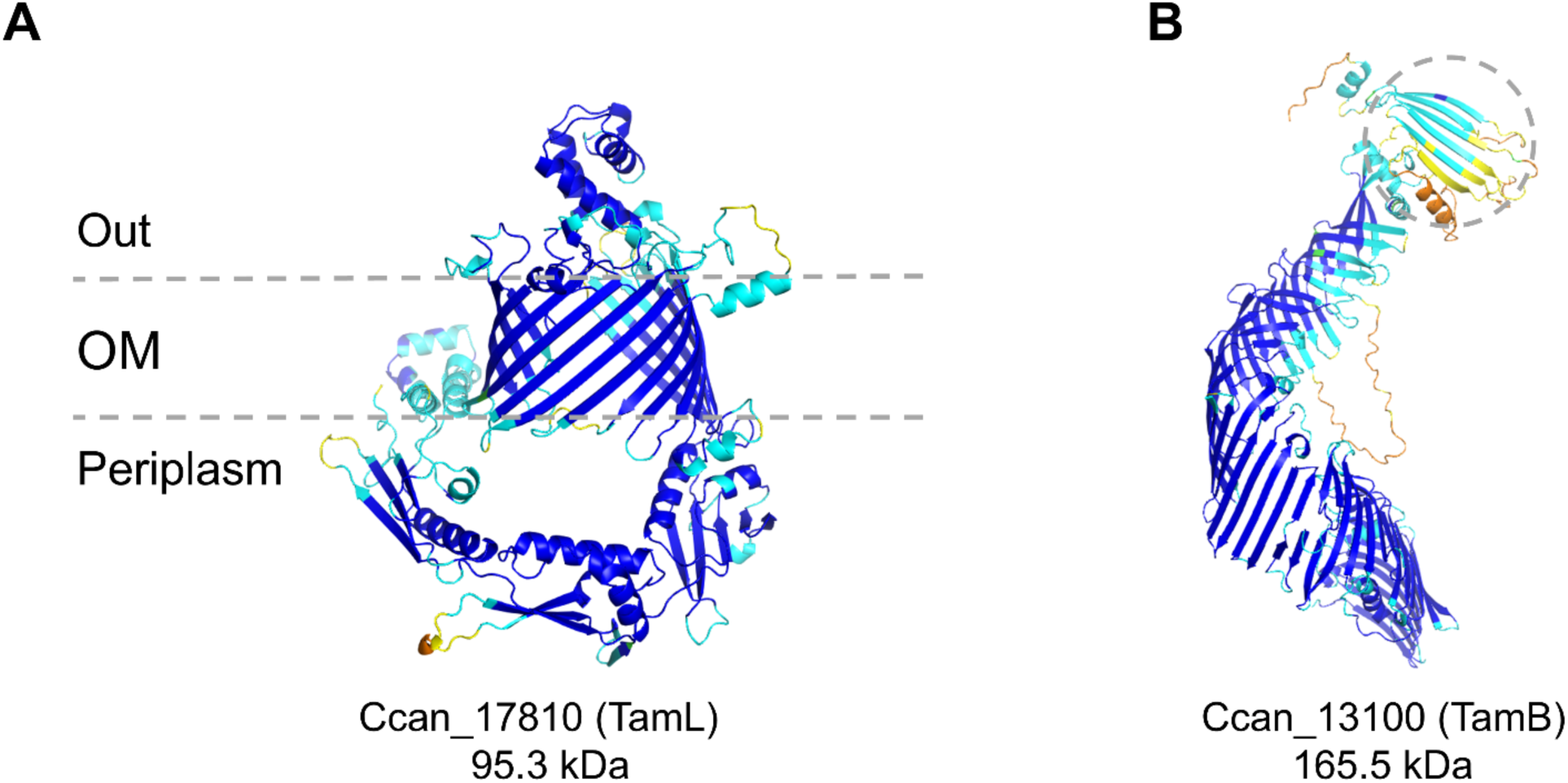
Three-dimensional structures of TamA (A), TamB (B) homologs in *C. canimorsus* as predicted by AlphaFold2 (https://alphafold.ebi.ac.uk) [98]. Protein names and the predicted molecular weights are reported below each structure. Structures are colored based on the per-residue confidence score (pLDDT) between 0 and 100: dark blue (pLDDT>90), cyan (90>pLDDT>70), yellow (70>pLDDT>50), and orange (pLDDT<50). In B, only the last C-terminal 1000 residues are shown. Moreover, the predicted C-terminal pseudosubstrate domain is highlighted via a dashed circle.

Finally, similarly to was observed in *F. johnsoniae*, the pseudosubstrate domain of TamB (residues 1346-1149) was predicted in our AlphaFold multimer model to establish physical interactions with the last strands of TamL β-barrel (Supplementary Figure S6B), thus indicating that such a mechanism of interaction could be typical of the TAM complexes from different bacteria.

## Discussion

In diderm bacteria the transport and assembly of outer membrane proteins (OMPs) are ensured by the BAM and TAM complexes, with BamA and TamA known to be their catalytic cores. Both proteins belong to the Omp85 family and share a similar mechanism for OMP folding and insertion into the OM [8–12]. While BamA is essential in all diderms, TamA is found only in Proteobacteria, where it primarily folds virulence factors during host growth [10,13,67,68]. This is why the TAM complex has always been thought to play a marginal role in OM homeostasis. Recent studies have revealed that TAM may have a more crucial role by promoting phospholipid transport across the two membranes, though this function is currently debated [69].

In Bacteroidetes and Chlorobi, a TamA-like lipoprotein, TamL, replaces TamA. Thanks to the lipid moiety located at the N-terminal end, TamL is likely anchored to the OM [28,29]. The reason TamL replaces TamA is unknown; nor is it clear whether TamL is still able to interact with TamB to form the TAM complex. To our knowledge, no research has been done on TamL so far. To provide new insights into a field of research that has remained largely unexplored, through this work we aimed to investigate the molecular function of TamL in *F. johnsoniae*, an environmental Bacteroidetes living on fresh water and soil, which has been extensively studied for gliding and Type 9 protein Secretion (T9SS) [35,38,40,53].

We first found that *F. johnsoniae* possesses two TamL (Fjoh_1464; Fjoh_1900) and two TamB (Fjoh_4592; Fjoh_1899) homologs, of which only Fjoh_1900 and Fjoh_1899 are probably encoded by genes in an operon given their close genomic proximity. The existence of multiple TamL and TamB proteins in Bacteroidetes was previously reported in *Porphyromonas gingivalis* [70], though the genomic organization of TamL and TamB encoding genes in *F. johnsoniae* is different. To our surprise, we also identified a putative TamA-like protein (Fjoh_0402), which has never been observed before outside Proteobacteria.

Interestingly, we found that TamL and TamB encoding genes, *fjoh_1464* and *fjoh_4592*, are essential, and that the lipidation occurring at the N-terminal end of Fjoh_1464 during its maturation is crucial for cell viability. Whether the lipidation is needed for Fjoh_1464 OM localization or its activity remains unknown. This finding was completely unexpected as, to our knowledge, there are no previous reports on the essentiality of the TAM subunits. The deletion of the other genes (*fjoh_0402*, *fjoh_1899,* and *fjoh_1900)* did not impact cell fitness in any of the tested conditions. Therefore, we concluded that, though *F. johnsoniae* possesses multiple TamL and TamB proteins, only Fjoh_1464 and Fjoh_4592 are critical for cell viability, and we thereby named them TamL and TamB, respectively; whereas TamL2 and TamB2 were assigned to Fjoh_1900 and Fjoh_1899, respectively. We observed that TamL and TamB do not show the same pattern of temporal regulation since TamL is exclusively expressed during the exponential phase of growth, whereas TamB appears constantly present within the cell. A possible explanation is that an active interaction between the two proteins, as evidenced by our pull-down assays (see below), might be specifically required in actively dividing cells. Consequently, the structural complexity of TamB, which spans the periplasm and the peptidoglycan (PG) sacculus, might make it more advantageous to downregulate TamL to deactivate TAM.

We sought to generate TamL and TamB depletion strains, but this turned out to be possible only for TamL. At present, we cannot explain why, though the genomic region where the *tamB* gene is located may play a role. Using the TamL depletion strain, we confirmed its essentiality in both liquid and solid media, which was accompanied by morphological defects, cell aggregations and increased budding of bigger outer membrane vesicles (OMVs). Furthermore, TamL depleted cells exhibit increased sensitivity to several OM perturbing agents, such as Polymyxin B, Vancomycin and SDS. Notably, cell fitness in TamL-depleted cells could be ameliorated by increasing the concentration of Mg^2+^ (30 mM in our assay) in the growth medium. Similar phenotypes were previously observed in *Escherichia coli* and *Brucella suis* [22,23,51,52,68]. It is important to note that these results were limited to TamB mutants, whereas our study is the first to apply them specifically to TamL. We could therefore conclude that TamL depletion causes the disruption of OM integrity, which eventually leads to cell death, thus explaining its essentiality.

We also found that TamL depletion significantly altered the OM proteome determining a sharp increment in OM lipoproteins. We indeed observed a strong enrichment in the LolA (Fjoh_2111, FC: 3.0; Fjoh_1085, FC: 5.7) and LolB (Fjoh_1084, FC: INF) homologs, which are members of the lipoprotein trafficking system, recently investigated also in *F. johnsoniae* [58]. Notably, while the observed upregulation of Fjoh_2111 (LolA1) could account for the increased OM lipoprotein trafficking upon TamL depletion, the sharp upregulation of the LolA and LolB homologs Fjoh_1085 (LolA2) and Fjoh_1084 (LolB2) is intriguing as to date no precise function has been assigned to them, though they are likely implicated in the biosynthesis and/or transport of the OM lipid flexirubin given their genomic proximity with the flexirubin biosynthesis genes [35].

OM lipoproteins are of crucial importance in bacterial survival, especially in maintaining the integrity and functionality of OM under stress conditions. Noticeable examples are the BAM accessory lipoproteins, the Braun lipoprotein (Lpp), and LolB of the Lol system, whose increased abundances were shown to be result of a stress response to OM damage [71–73]. Therefore, it is plausible that TamL depletion disrupts OM integrity, thus triggering a stress response that leads to an increase in OM lipoproteins. In fact, it should be recalled that TamL is a lipoprotein itself and our data clearly demonstrate its involvement in OM stability.

On the other hand, we did not observe an overall downregulation of integral/β-barrel OM proteins, as one might expect from the depletion of a foldase mostly involved in their transport and folding. Prior studies have instead shown that the depletion of BamA and BamD, the two essential subunits of the BAM complex (the main OMP assembly factor in diderm bacteria), were accompanied by a significant reduction in the OM protein content [46,47,74–76]. Therefore, our proteomics data would not support the hypothesis that TamL functions as a general OM foldase in *F. johnsoniae*.

We found that the OMVs released from TamL-depleted cells were twice as big as the control strain, highly enriched in periplasmic proteins, as well as almost completely devoid of OMPs. This finding reinforces our model of TamL as a crucial player in maintaining OM integrity, as its depletion destabilizes the OM and leads to the release of no physiological OMVs lacking their functional protein cargo. Strikingly, we observed that surface-exposed lipoproteins, found enriched in the OM fractions, were also reduced in the OMVs. Since these lipoproteins constitute a significant portion of OMV cargo in Bacteroidetes [77,78], we reasonably assume that TamL may be required to some extent for their incorporation into OMVs.

Except for a slight increment in phosphatidylethanolamine (PE) in the OMVs, we did not observe any significant impact on the main OM lipid content in TamL-depleted cells. Furthermore, in the TamL depletion strain the deletion of the *fjoh_2419* and *fjoh_0833* genes, responsible for the biosynthesis of sulfonolipids (SLs) and ornithine lipids (OLs), respectively [34,60], did not result in the same loss of cell viability observed in the parental strain. Instead, it further impaired cell fitness, suggesting an additive effect on the OM instability. Based on these findings, we concluded that TamL is most likely not involved in OM lipid transport in *F. johnsoniae*.

We then confirmed that TamL and TamB establish physical interaction *in vivo,* and thereby likely form the subunits of the TAM complex in *F. johnsoniae*. Therefore, we find it reasonable to assume that this newly identified TAM complex operates similarly to what is already described in Proteobacteria, with TAM substrate(s) passes through TamB before reaching TamL [13,14,69]. Consequently, the loss of TamL could result in the stalling of TAM substrate(s) on TamB, which could ultimately be detrimental for cell viability by depriving cells of “free” TamB. In this context, the upregulation of TamB we observed upon TamL depletion may represent a cellular attempt to cope with the stalling of substrate(s) on TamB. Interestingly, the sharp upregulation of TamA, observed by our proteomics studies of the OM fractions, is also intriguing. In fact, in *F. johnsoniae,* and by extension in those Bacteroidetes species where *tamA* has been identified by our *in silico* analysis (see below), *tamA* is not part of the a transcriptional unit with a *tamB-*like gene. Therefore, we cannot exclude that in *F. johnsoniae* TamB could also interact with TamA, which might explain why both proteins are sharply upregulated upon TamL depletion. However, likely due to the key structural differences between TamA and TamL which may affect substrate affinities, TamA would not be able to efficiently take over TamL, which may explain why the deletion of *tamA* is not lethal and has no effect on TamL-depleted cells.

Our *in silico* analysis indicates that the presence of monocistronic *tamL* and *tamB* genes is not a genomic trait restricted to *F. johnsoniae* but it is rather a common feature across Bacteroidetes. *C. canimorsus,* a human pathogen primarily recognized as a commensal in the oral cavities of cats and dogs, serves as an example [66]. In this species, we indeed confirmed the essentiality of the TamL and TamB homologs (Ccan_17810 and Ccan_13100, respectively) and our AlphaFold2-based model predicts their interaction. Altogether, these findings open to the possibility that both TAM subunits may be also essential in other Bacteroidetes species. Likewise, the identification of the TamA homologs in other Bacteroidetes clearly suggests that TamA and TamL co-existence in the OM is not restricted to *F. johnsoniae*.

It remains to be understood the molecular contribution of TamL to OM integrity. Several possibilities may be proposed. For example, TamL may be required to fold and insert into the OM one or more proteins that are essential for the stability of the OM. Additionally, as already observed for BamA in *E. coli* [79], TamL may interact with PG, and this interaction may be necessary to give stability to the cell-wall. Still, the POTRA domains of TamA in *P. aeruginosa* have been recently shown to establish interaction with the polar heads of phospholipids, thus modulating membrane properties [16]. We do not exclude that the same interaction might occur between TamL and the OM lipids.

## Materials and Methods

### Bacterial strains and media

*Flavobacterium johnsoniae* UW101 WT (ATCC 17061) and derivative strains were grown in Casitone-Yeast Extract (CYE) medium (10 g/L casitone, 5 g/L yeast extract, 4 mM MgSO_4_ and 10 mM Tris-HCl, pH 7.5) or motility medium (MM) at 30°C [80,81]. *Escherichia coli* strains were grown in Lysogeny Broth (LB) medium at 37°C. All strains used in this study are listed in Supplementary Table S4. When needed, the following antibiotics were used: ampicillin (100 μg/ml); streptomycin (100 µg/ml), gentamicin (20 µg/ml), erythromycin (100 μg/ml); tetracycline (20 μg/ml).

### Strain construction

Plasmids and oligonucleotides used in this study are listed in Supplementary Tables S5 and S6, respectively. Plasmids for mutagenesis of *F. johnsoniae* were generated by either Gibson assembly [82], or by sequential cloning into the non-replicative plasmid pYT354 (Supplementary Table S5). In the former case, plasmid pYT354 and around 2000 base pairs (2 kb) upstream and downstream the region of interest were PCR-amplified by Q5® High-Fidelity DNA Polymerase (NEB) using primers with 20 overlapping base pairs designed with Benchling. Approximately 5 ng of pYT354 and *F. johnsoniae* genomic DNA (gDNA) were used as a template. All the PCR-generated fragments were then assembled by Gibson assembly and cloned in *E. coli* Top10 by electroporation. The Gibson reaction was assembled on ice with 10 µl of the 2x Gibson mix to which the plasmid fragment (80-90 ng) and insert fragments were added in a 1:3:3 or 1:5:5 ratio. Then, the reaction was kept at RT for 30 s and incubated overnight at 50°C. The 2x Gibson mix was made *in-house*. Shortly, for 800 µl final volume, 0.64 µl T5 exonuclease (10 U/µl, NEB), 20 µl Q5® High-Fidelity DNA Polymerase (2 U/µl, NEB), 160 µl *Taq* DNA ligase (40 U/µl, NEB), 320 µl 5x reaction buffer (see below) and 300 µl of dH_2_O. The 5× reaction buffer consisted of 0.5 M Tris-HCl pH 7.5, 50 mM MgCl2, 1 mM of each dNTPs, 50 mM DTT, ¼ W/V PEG 8000, 5 mM NAD and dH_2_O.

For the deletion of *fjoh_1464* (*tamL*), *fjoh_4592* (*tamB*), *fjoh_0402* (*tamA*), *fjoh_1900* (*tamL2*) and *fjoh_0833*, a sequential cloning strategy was performed. Briefly, for the deletion of *fjoh_0402* (*tamA*) and *fjoh_0833* around 2 kb upstream and downstream the region of interest were PCR-amplified using approximately 5 ng of *F. johnsoniae* genomic DNA (gDNA) as a template, then digested for 2-3 hours at 37°C using 1 µl of SphI/XhoI (upstream fragment) and of XhoI/XbaI (downstream fragment). For the deletion of *tamL2*, BamHI/KpnI (upstream fragment) and KpnI/SphI (downstream fragment) were instead used. For the deletion of *tamL* and *tamB*, and for the 21cys-gly substitution of *tamL*, upstream and downstream fragments were first PCR-amplified and then PCR-overlapped prior to digestion with 1 µl of ApaI/SpeI (*tamL* deletion and 21cys-gly substitution) or BamHI/SphI (*tamB* deletion). Digested fragments were then sequentially inserted into pYT354 (50-80 ng), previously digested with the same enzymes and incubated for 1 hour at 37°C with 1 µl of alkaline phosphatase from calf intestine (20.5 U/µl; Roche). Ligation reactions were carried out overnight at 16°C using 1µl of T4 DNA Ligase (20 U/µl; NEB). Plasmids were then cloned in *E. coli* Top10 by transformation.

All the generated suicide plasmids were introduced in the appropriate *F. johnsoniae* background strain by triparental mating, using *E. coli* Top10 and *E. coli* MT607 as donor and helper strains, respectively (Supplementary Table S4). Erythromycin resistance was used to select colonies with chromosomally integrated plasmid. After PCR screening, one single colony was picked and incubated in 5 ml LB overnight at 30°C under vigorous shaking to promote plasmid loss, and then plated onto LB agar plates containing 5% (w/v) *D*-sucrose. Plates were incubated for 2-3 days at 30°C until single colonies were visible. Colonies resistant to *D*-sucrose were selected and PCR-screened for the presence of the desired genomic mutation. When needed, 1 mM isopropyl-beta-D-thiogalactopyranoside (IPTG; ThermoFisher Scientific) was added to the plates.

### Construction of a TamL depletion strain

We generated the P_ompA_::*lacI* construct (Supplementary Table S5) by putting the gene coding for the LacI transcription repressor (*lacI*) under the regulation of the promoter of *fjoh_0697*, encoding the outer membrane protein OmpA. This construct was then cloned in the suicide plasmid pYT354 carrying the homologous regions to *fjoh_0061* and *fjoh_0062*, thus generating the plasmid pYT354-P_ompA_::*lacI* (Supplementary Table S5) which allowed the insertion of this construct in the genomic locus between *fjoh_0061* and *fjoh_0062*. Afterward, we took advantage of a prior study where an IPTG-inducible expression vector for *Bacteroides fragilis* was generated [83]. Shortly, we engineered the constitutive promoter of the cefoxitin resistance gene (P_cfxA_) by adding the binding sequences of the LacI transcription repressor (*lacO_3_* and *lacO_1_*) upstream of the −33 binding box and downstream of the transcriptional initiation site (TIS) of P_cfxA_. Furthermore, upstream of lacO_3_ we added the transcription terminator site of *fjoh_0014* (*ter_fjoh_0014_*), thus generating the *ter_fjoh_0014_*-*lacO_3_*-P_cfxA-lacO1_ construct (P_cfxA-lacO_). Then, we constructed a plasmid (Supplementary Table S5) where either *tamL* or *tamB* were cloned downstream of this construct to generate a strain where their native promoters were replaced with P_cfxA-lacO_.

### Assessment of TamL and TamB profile during cell growth

From a fresh LB plate, the *F. johnsoniae* double-tagged strain was inoculated at OD_600_:0.05 in 10 ml of CYE medium and incubated at 30°C, 160 rpm for 24 hours (pre-culture). Then, the OD_600_ was measured, and accordingly bacterial cells were inoculated in 25 ml CYE medium at OD_600_:0.05, and incubated at 30°C, 160 rpm. At defined time points, samples were collected for immunoblot and bright-field microscopy analyses. Notably, to collect all the indicated time points, a second culture was grown in parallel.

The same protocol was applied to grow the *F. johnsoniae* TamL depletion strain (P_ompa_::*lacI*-P_cfxA-lacO_-*tamL*), though IPTG at the final concentration of 1 mM was added to the pre-culture medium (permissive condition). Afterwards, for the growth in 25 ml CYE medium, the pre-culture was inoculated in two flasks: one with 1 mM IPTG (permissive condition) and one without (non-permissive condition).

### Outer membrane permeability assays

*F. johnsoniae* cultures were grown overnight (20-22 hours) in CYE medium at 30°C at 160 rpm. Only for the TamL depletion strain (P_ompa_::*lacI*-P_cfxA-lacO_-*tamL*), IPTG at the final concentration of 1 mM was added to the medium. Upon measurement of the OD_600_, a culture volume corresponding to OD_600_:1 was harvested at 5,000 x g for 5 min at RT. Bacterial pellet was washed in 1 ml of 1x PBS, and then harvested again as before. Afterwards, cells were diluted to OD_600_: 0.05 in 1 ml of fresh CYE or MM medium. When needed, 1 mM IPTG was added. 200 µl (in triplicate) of cells at OD_600_: 0.05 was deposited onto a 96-well microplate and growth was monitored for 36 hours with continuous shaking at 30°C, in an automated plate reader (Bioscreen C, Lab Systems) measuring the OD_600_ every 10 min. Data are displayed as mean ± standard deviation (SD) of at least three biological replicates. For stress tolerance assays, after washing in 1x PBS, cells were resuspended in fresh CYE medium in the presence of the following compounds: Polymyxin B (40 μg/ml), SDS (0.0025%, w/v), and Vancomycin (62.5 μg/ml). The powders of the three compounds were from Sigma-Aldrich. On the day of the experiment, powders were dissolved in MilliQ water and passed through a 0.2 µm-filter in a sterile environment.

### Separation of the inner membrane and outer membrane

From a previously described protocol [84] we implemented a few substantial modifications to decrease the inner membrane (IM) content in the outer membrane (OM) fractions. Briefly, cells were grown in 10 ml of CYE medium containing 0.25 mM IPTG at 30°C for 12 hours at 160 rpm (pre-culture). Then, the OD_600_ was spectrophotometrically measured, and a culture volume corresponding to OD_600_:2 was harvested at 5,000 x g for 5 min at RT. The bacterial pellet was then resuspended in 1 ml of 1x PBS, and the previous step was repeated again. Afterwards, cells were inoculated at OD_600_:0.05 in two flasks containing 50 ml of CYE medium at 30°C, 160 rpm. IPTG, at the final concentration of 1 mM, was added to just one of the two flasks (permissive condition), while no IPTG was added to the second flask (non-permissive condition). After 12 hours of growth, the OD_600_ of the cultures was measured and the volume equivalent of OD_600_:40 was harvested at 7,000 x g for 10 min at 4°C. In parallel, 1 tablet of protease inhibitors (cOmplete™, Mini Protease Inhibitor Cocktail, EDTA-free) was added to 10 ml of a solution consisting in 10 mM HEPES, pH 7.4, 1 mM EDTA. Bacterial pellet was then resuspended in 4 ml of this solution on ice. Cells were lysed by 2-3 passages through a cell disrupter (Constant Systems) at 35,000 psi. Cell debris and unlysed cells were removed by centrifugation at 2,500 x g for 10 min at 4°C. Before the next step, 200 µl of supernatant was withdrawn and kept at 4°C for downstream analyses. Then, cell extracts (2.8-3.0 ml) were centrifuged at approximately 108,000 x g for 107 min at 4°C using an ultracentrifuge device equipped with a TLA 100.3 fixed-angle rotor (Beckman). The supernatants were retained as soluble fractions and kept at 4°C for downstream analyses, while the pellets, corresponding to the membrane fractions (i.e., inner and outer membranes together), were resuspended in 2 ml of a fresh solution consisting in 10 mM HEPES, pH 7.0. 200 µl of this was withdrawn and kept at 4°C for downstream analyses. The membrane fractions were further fractionated by partial solubilization in 2% (w/v) sodium n-lauroylsarcosinate solution on a rotating wheel for 1 hour at RT, followed by a second round of ultracentrifugation performed as described above. The supernatants, corresponding to solubilized inner membranes, were retained and stored at 4°C, while the pellets, corresponding to the outer membranes, were resuspended in 500 µl of ice-cold 10 mM HEPES, pH 7.4. Unless otherwise specified, all samples were temporarily kept at 4°C to assess sample purity, and then stored at −80°C.

### Assessment of membrane sample purity

The succinate dehydrogenase (SDH) activity was measured as described elsewhere [85] as an indicator for inner membrane contamination in the outer membrane fraction. Notably, due to the presence of the Sarkozyl detergent in the inner membrane fraction, the SDH activity could not be detected in this fraction. Briefly, after membrane extraction, the protein concentration of the different fractions was quantified using the Quick StartTM Bradford Protein Assay (Bio-rad). Accordingly, all the fractions were then diluted to 0.24 mg/ml of total proteins. A 96-well plate was filled in duplicate with either 100 μl of sample, corresponding to 24 µg of total proteins, or dH_2_O. 60 μl of the following freshly made reaction mix was then added to each well: 50 mM Tris-HCl, pH 8.0, 4 mM KCN and 40 mM disodium succinate. After 5 min of incubation at RT, 20 μl of 4 mM DCIP and 20 µl of 2 mM PMS were subsequently added. For the detection of the SDH activity, the absorbance at 600 nm was measured using a SpectraMax ID3 Molecular Devices fluorimeter every 30 seconds for 1 hour at 25°C.

To examine the total protein profile of cell fractions, 2 µg of total proteins were loaded onto 12% SDS-PAGE and then visualized by AgNO_3_ staining as described elsewhere [86].

### Proteomic analyses by mass spectrometry

#### Sample preparation

For the analysis of the outer membrane proteome, 25 µg of total proteins in 5 mM HEPES pH 7.4, from 5 independent biological replicates were given for protein digestion, while 20 µg of total proteins in an *in-house* made 1x Laemmli buffer (47 mM Tris-HCl, pH 6.8, SDS 1% (w/v), 50 mM DTT, glycerol 10% (v/v) and traces of bromophenol blue) from 4 independent biological replicates were used for the OMVs proteome analysis. For the proteomic analysis of the elution fractions from pull-down assays, 25 µg of total proteins in 1xPBS from 2 independent biological replicates was given. Prior to protein digestion, the chemical crosslinker (DTSSP) was cleaved by adding 40 mM DTT and incubating at 37°C for 30 min.

#### Protein digestion

The samples were treated using Filter-Aided Sample Preparation (FASP) using the following protocol. Briefly, to first wash the filter, 100 µl of 1% formic acid were placed in each Millipore Microcon 30 MRCFOR030 Ultracel PL-30 before centrifugation at 14,500 rpm for 15 min. For each sample, proteins were resuspended in 150 µl of urea buffer 8 M (urea 8 M in buffer Tris 0.1 M, pH 8.5), placed individually in a column, and centrifuged at 14,500 rpm for 15 min. The filtrate was discarded, and the columns were washed three times by adding 200 µl of urea buffer followed by a centrifugation at 14,500 rpm for 15 min. For the reduction step, 100 µl of dithiothreitol (DTT) were added and mixed for 1 min at 400 rpm with a thermomixer before an incubation of 15 min at 24°C. Samples were then centrifugated at 14,500 rpm for 15 min, the filtrate was then discarded, and the filter was washed by adding 100 µl of urea buffer before another centrifugation at 14,500 rpm for 15 min. An alkylation step was performed by adding 100 µl of iodoacetamide (IAA, in urea buffer) in the column and mixing at 400 rpm for 1 min in the dark before an incubation of 20 min in the dark and a centrifugation at 14,500 rpm for 10 min. To remove the excess of IAA, 100 µl of urea buffer were added and the samples were centrifugated at 14,500 rpm for 15 min. To quench the rest of IAA, 100 µl of DTT were placed on the column, mixed for 1 min at 400 rpm and incubated for 15 min at 24°C before centrifugation at 14,500 rpm for 10 min. To remove the excess of DTT, 100 µl of urea buffer were placed on the column and centrifuged at 14,500 rpm for 15 min. The filtrate was discarded, and the column was washed three times by adding 100 µl of sodium bicarbonate buffer 50 mM (ABC, in ultrapure water) followed by a centrifugation at 14,500 rpm for 10 min. The last 100 µl were kept at the bottom of the column to avoid any evaporation in the column. The digestion process was performed by adding 80 µl of mass spectrometry grade trypsin (1/50 in ABC buffer) in the column and mixed at 400 rpm for 1 min before an incubation overnight at 24°C in a water saturated environment. The Microcon columns were placed on a LoBind tube of 1.5 ml and centrifuged at 14,500 rpm for 10 min. 40 µl of ABC buffer were placed on the column before centrifugation at 14,500 rpm for 10 min. Trifluoroacetic acid (TFA) 10 % in ultrapure water were added to the contain of the LoBind Tube to obtain 0.2 % TFA. The samples were dried in a SpeedVac up to 20 µl and transferred to an injection vial.

#### Mass Spectrometry (MS)

The digest was analyzed using nano-LC-ESI-MS/MS tims TOF Pro (Bruker, Billerica, MA, USA) coupled with an UHPLC nanoElute2 (Bruker). The different samples were analyzed with a gradient of 60 min. Peptides were separated by nanoUHPLC (nanoElute2, Bruker) on a 75 μm ID, 25 cm C18 column with integrated CaptiveSpray insert (Aurora, ionopticks, Melbourne) at a flow rate of 200 nl/min, at 50°C. LC mobile phases A was water with 0.1% formic acid (v/v) and B ACN with formic acid 0.1% (v/v). Samples were loaded directly on the analytical column at a constant pressure of 800 bar. The digest (1 µl) was injected, and the organic content of the mobile phase was increased linearly from 2% B to 15 % in 22 min, from 15 % B to 35% in 38 min, from 35% B to 85% in 3 min. Data acquisition on the tims TOF Pro was performed using Hystar 6.1 and timsControl 2.0. tims TOF Pro data were acquired using 160 ms TIMS accumulation time, mobility (1/K0) range from 0.75 to 1.42 Vs/cm². Mass-spectrometric analyses were carried out using the parallel accumulation serial fragmentation (PASEF) [87] acquisition method. One MS spectra followed by six PASEF MSMS spectra per total cycle of 1.16 s.

#### Data analysis

Data analysis was performed using Mascot 2.8.1 (Matrix Science). For database searching, tandem mass spectra were extracted, charge state deconvoluted and deisotoped by Data analysis (Bruker) version 5.3. All MS/MS samples were analyzed using Mascot (Matrix Science, London, UK; version 2.8.1). Mascot was set up to search the *Flavobacterium johnsoniae* UW101 proteome from Uniprot (220718-5021 entries) and a contaminants database, assuming the digestion enzyme trypsin with two missed cleavages. Mascot was searched with a fragment ion mass tolerance of 0.050 Da and a parent ion tolerance of 15 ppm. Carbamidomethyl of cysteine was specified in Mascot as fixed modifications. Oxidation of methionine, deamidation of asparagine and glutamine and acetyl of the n-terminus were specified in Mascot as variable modifications.

For protein identification, Scaffold (version Scaffold_5.1.1, Proteome Software Inc., Portland, OR) was used to validate MS/MS based peptide and protein identifications. Peptide identifications were accepted if they could be established at greater than 96.0% probability to achieve a False Discovery Rate (FDR) less than 1.0% by the Scaffold Local FDR algorithm. Protein identifications were accepted if they could be established at greater than 5.0% probability to achieve an FDR less than 1.0% and contained at least 2 identified peptides. Protein probabilities were assigned by the Protein Prophet algorithm (Nesvizhskii, Al et al Anal. Chem. 2003;75(17):4646-58). Proteins that contained similar peptides and could not be differentiated based on MS/MS analysis alone were grouped to satisfy the principles of parsimony. Proteins sharing significant peptide evidence were grouped into clusters.

### *In silico* characterization of the proteins identified by MS

Predicted localization and topology analysis for proteins identified by MS were performed using UniProt database, SignalP 6.0 [44], PSORTb 3.0 [88] and CAZy database [89]. All predicted lipoproteins were manually checked for the presence of the lipoprotein export signal (LES) [32]. When present, they were assigned as surface-exposed lipoproteins; when absent, they were assigned as periplasm-facing lipoproteins.

### Outer membrane vesicles (OMVs) purification and protein quantification

*F. johnsoniae* cultures were grown as described in the membrane fractionation protocol with the sole difference that the volume of the cultures was increased up to 1 L and 200 ml for cultures in permissive (+ IPTG) and non-permissive (- IPTG) conditions, respectively. After 12 hours of growth, bacterial cells were pelleted twice at 5,000 x g for 10 min at 4°C, and then the culture supernatants were passed through a 0.22 μm pore-size filter (Sarstedt) to remove any cell leftover or debris. Afterwards, supernatants were concentrated down to 3 ml using Amicon® Ultra-15 Centrifugal Filter Unit (50 kDa cutoff, Millipore) centrifuging at 3,500 x g for 10 min at 4°C each time. The concentrated supernatants were then centrifuged at approximately 108,000 x g for 3 h at 4°C using the same ultracentrifuge device employed for the membrane fractionation protocol. Supernatants were discarded and the pellets, corresponding to the OMVs, were weighed and stored at −80°C until use.

To quantify the protein content, OMVs were resuspended in an *in-house* made1x Laemmli buffer (47 mM Tris-HCl, pH 6.8, SDS 1% (w/v), 50 mM DTT, glycerol 10% (v/v) and traces of bromophenol blue) and boiled at 99°C for 5 min. Afterwards, the protein concentration was estimated by Pierce BCA protein assay (ThermoFisher Scientific). To get rid of the SDS interference, the ionic detergent compatibility reagent (ThermoFisher Scientific, product reference: 22663) was used in agreement with the manufacturer’s instructions. The OMVs protein profile was examined as described above for membrane fractions.

### Total lipid extraction and analyses from lipid fractions and OMVs

#### Lipid extraction

For total lipid extraction, we adapted a previously described protocol [90]. Shortly, OMVs (4 mg, dry weight) were resuspended in 500 µl of 1x PBS and placed in a glass tube, to which 650 µl of chloroform and 1.3 ml of methanol were added. Samples were vigorously vortexed for 5 min, and then incubated at 30°C, 160 rpm for 3 hours. During the incubation, samples were vigorously vortexed for 1 min every 15 minutes. Afterwards, 650 µl of chloroform and 650 µl of MilliQ water were added, and samples were incubated for 30 min as before. To separate the organic lipid-rich phase from the aqueous phase, samples were centrifuged at 3,220 x g for 5 min at RT. The organic phase was then transferred to a glass vial, which was kept under a laminar flow hood overnight at RT to allow organic solvents to evaporate. The glass vials containing the dry pellets (lipid extracts) were stored at −20°C.

#### Lipid analysis by Thin Layer Chromatography (TLC)

Lipid extracts were first resuspended in 50 µl of chloroform:methanol (1:1) mixture, and then 15 µl (5µl at a time) were spotted via a glass micro-syringe onto a TLC aluminum plate (HPTLC Silica gel 60 F254 10×10 cm, MERCK), 1 cm from the bottom of the plate. For lipid separation, a solvent system of chloroform:methanol:ammonium hydroxide (140:60:10, v/v/v) was used as previously described [34]. After migration, the TLC plate was dried out at RT for 1 hour under a laminar flow hood. Then, to visualize lipids, the TLC plate was first placed in a glass jar saturated with iodine (I_2_) vapor from I_2_ crystals, exposed to for 30 min at RT, and then photographed. Afterwards, the TLC plate was kept at RT for 1 hour to allow I_2_ to decolorize. Then, the same TLC plate was soaked with ninhydrin (0.2% w/v in ethanol) for a few seconds, and then incubated at 150°C for 10 min to develop spots. For 2D-TLC, lipids were separated using the same solvent system described above for the first dimension, whereas a solvent system consisting of chloroform:methanol:glacial acetic acid:acetone:water (130:10:10:20:3, v/v/v/v/v) was used for the second dimension. Lipids were visualized by spraying a solution of primuline (0.05% w/v in a water/acetone mixture (4:1)) [91]. To develop spots, plates were dried out at RT for 15 min, and then exposed to UV-light (364 nm). For the relative quantification of each lipid class ImageJ was used.

#### Lipidomic analysis by LC-MS

Lipid extracts from the leftover volume (35 µl) were dried out under a laminar flow and then analyzed by LC-MS. Normal phase (NP) LC was performed on an Agilent 1200 Quaternary LC system equipped with an Ascentis Silica HPLC column, 5 μm, 25 cm x 2.1 mm (Sigma-Aldrich, St. Louis, MO) as previously described [92]. Mobile phase A was a mixture of chloroform/methanol/aqueous ammonium hydroxide (800:195:5, v/v/v); mobile phase B was a mixture of chloroform/methanol/water/aqueous ammonium hydroxide (600:340:50:5, v/v/v); mobile phase C was a mixture of chloroform/methanol/water/aqueous ammonium hydroxide (450:450:95:5, v/v/v). The elution program was as follows: 100% mobile phase A was held isocratically for 2 min and then linearly increased to 100% mobile phase B over 14 min and held at 100% B for 11 min. The LC gradient was then changed to 100% mobile phase C over 3 min and maintained at 100% C for 3 min, and finally returned to 100% A over 0.5 min and maintained at 100% A for 5 min. The LC eluent (with a total flow rate of 300 μL/min) was introduced into the ESI source of a high-resolution TripleTOF5600 mass spectrometer (Sciex, Framingham, MA). The instrumental settings for negative ion ESI and MS/MS analysis were as follows: IS= −4500 V; CUR= 20 psi; GSI= 20 psi; DP= −55 V; and FP= −150 V. For MS/MS analysis nitrogen was used as the collision gas. Analyst TF1.5 software (Sciex, Framingham, MA) was used for data analysis.

### *In vivo* crosslinking and TamL pull-down assays

The double tagged strain *F. johnsoniae* (3xFLAG)TamL/(2xStrep)TamB was used to pull-down TamL, while the single tagged strain (2xStrep)TamB was used as mock (Supplementary Table S4). From overnight pre-cultures, strains were inoculated at the optical density (OD_600_) of 0.05 in 200 ml of CYE at 30°C, 160 rpm. When OD_600_ reached 0.6-0.8 (usually between 4-5 hours), bacterial cultures were first harvested at 5,000 x g at 4°C for 20 min, washed in 25 ml of 1x PBS, and harvested again at 5,000 x g at 4°C for 10 min. Pellets were resuspended in 10 ml of 1x PBS, and 3,3’- dithiobis(sulfosuccinimidyl propionate) (DTSSP, Sigma-Aldrich) at the final concentration of 1 mM was added for *in vivo* crosslinking. The DTSSP-treated cultures were incubated at 30°C for 30 min under moderate shaking (125 rpm). Then, a solution of Tris-HCl, pH 7.5, at the final concentration of 100 mM was added to quench DTSSP, and cultures were left at 30°C for 30 min at 125 rpm. Cultures were then harvested at 5,000 x g at 4°C for 10 min, washed in 20 ml 1x PBS, and harvested again at 5,000 x g at 4°C for 10 min. Pellets were stored overnight at −80°C. The day after, pellets were resuspended in 3 ml of a solution consisting of 20 mM Tris-HCl, pH 7.5, 150 mM NaCl, 1 mM EDTA, lysozyme (0.4 mg/ml), DNase-I (30 µg/ml), 1% n-Dodecyl-β-D-Maltoside (DDM, ThermoFisher Scientific^TM^), containing half tablet of EDTA-free protease inhibitors (Roche cOmplete). Cells were incubated on ice for 30 min, and then lysed by two passages through a cell disruptor (Constant Systems) at 35,000 psi. To remove unlysed cells and cell debris, the lysates were centrifuged twice at 2,700 x g at 4°C for 10 min. Supernatant volume was measured and then diluted by adding the equivalent of 1.5 times its volume of a solution consisting of 10 mM Tris-HCl, pH 7.5, 150 mM NaCl, 0.5 mM EDTA, 1% DDM. 25 µl of pre-washed FLAG-beads (DYKDDDDK Fab-Trap™ Agarose, ChromoTek) was added to each supernatant and incubated for 2 hours at 4°C on a tube roller under moderate agitation. Afterwards, to remove unbound material, supernatants were centrifuged at 2,500 x g at 4°C for 10 min. Pellets, containing the FLAG-beads, were washed 5 times in 500 µl of 10 mM Tris-HCl, pH 7.5, 150 mM NaCl, 0.5 mM EDTA, 0.005% DDM according to the manufacturer’s instructions. Crosslinked (3xFLAG)TamL was then eluted by incubating the beads with 50 µl of 200 mM glycine, pH 2.5. The elution was repeated two more times. The total protein content of the eluates was determined by Pierce™ BCA Protein Assay (ThermoFisher Scientific^TM^) and the presence of 3xFLAG-tagged TamL in the eluates was assessed by immunoblot by resolving 2 µg of total proteins onto a polyacrylamide gel (8%). In parallel, eluate purity was determined by resolving 2 µg of total proteins onto a polyacrylamide gel (12%) followed by AgNO_3_ staining [86]. The eluates with the highest quantity of (3xFLAG)TamL and protein purity when confronted with their mocks were pooled and analyzed by Mass Spectrometry (MS).

### Immunoblot analysis

Whole-cell lysates were obtained at the specified time points. To normalize samples by cell density, a culture volume corresponding to OD_600_:1 was withdrawn and centrifuged at 5,000 x g for 5 min at RT, and the resulting pellet was resuspended in 100 µl of an *in-house* made 1x Laemmli sample buffer (47 mM Tris-HCl, pH 6.8, SDS 1% (w/v), 50 mM DTT, glycerol 10% (v/v) and traces of bromophenol blue) and stored at −20°C until use. Prior to gel loading, samples were thawed on ice for 10 min, boiled at 99°C for 5 min, and centrifuged at high speed (17,000 x g) for 3 min. 10 µl of sample was loaded onto SDS-polyacrylamide gels (8%) for electrophoresis. Proteins were then transferred onto a nitrocellulose membrane at 25V for 30 min by utilizing a semi-dry transfer cell (Bio-Rad). For (3xFLAG)TamL and GroEL detection, blocking was carried out overnight at 4°C in 5% (w/v) non-fat dry milk in phosphate buffer saline (PBS) with 0.05% (w/v) Tween 20 (PBST), and membranes were probed with anti-FLAG M2 (1:5,000; Sigma-Aldrich, catalogue number: F3165), StrepMAB-Classic (1:2,500; IBA-Lifesciences, catalogue number: 2-1507-001), or anti-GroEL (1:160,000; Sigma-Aldrich, catalogue number: G6532), for 3 and 1 hour, respectively, at RT. For (2xStrep)TamB detection, blocking was performed for 1 hour at RT in 5% (w/v) non-fat dry milk in PBST, and then membranes were probed with StrepMAB-Classic as described above overnight at 4°C. Thereafter, membranes were immunoblotted for 1 hour with secondary antibodies (1:5,000) anti-mouse or anti-rabbit linked to peroxidase (reference number: P0260 and P0217, respectively, Dako Agilent). Signal for (3xFLAG)TamL was generated using Clarity™ Western ECL substrate chemiluminescence reagent (BioRad), while signal for GroEL and (2xStrep)TamB was obtained using KPL LumiGLO Reserve Chemiluminescent Substrate Kit (SeraCare). An Amersham Imager 600 (GE Healthcare) was used for signal detection. Blots in figures are representative of at least three independent experiments.

### Microscopy techniques

#### Bright field microscopy

At the specified time points, cells grown in CYE with 1 mM IPTG (permissive condition) or with no added IPTG (non-permissive condition) were first diluted to OD_600_:1, and then 100 µl was fixed in the presence of 300 µl of 4% (w/v) PFA for 15 min at RT. Afterwards, fixed cells were harvested at 5,000 x g for 5 min at RT, then resuspended in 100 µl of sterile 1x PBS, and stored at 4°C. When needed, 3 µl of fixed bacteria was spotted on 1% agarose PBS pads, and pictures were taken using an Axioscop (Zeiss) microscope with an Orca-Flash 4.0 camera (Hamamatsu) and Zen Softmax Pro software (Zeiss).

#### Transmission electron microscopy (TEM) of F. johnsoniae cells

For *F. johnsoniae* growth, the same protocol as the membrane fractionation experiments was used, with the sole difference that an equivalent of OD_600_:3 was harvested. Then, samples were prepared for TEM analysis following an already described protocol [93]. Briefly, bacterial pellets were resuspended in 400 µl of 2.5% glutaraldehyde and 0.1% cacodylate mixture and incubated for 2 hours and 30 min at 4°C. Then, pellets were spun at 5,000 x g for 3 min, resuspended in 400 µl of 0.2% cacodylate, incubated for 10 min at RT, and then spun again at 5,000 x g for 3 min. This was repeated two more times. Pellets were then resuspended in 400 µl of 2% osmic acid and 0.1% cacodylate mixture and incubated 1 hour at 4°C. Afterwards, pellets were washed three times in 400 µl 0.2% cacodylate as before. Samples were kept at 4°C for a few days/weeks.

Samples were dehydrated by performing two washes in increasing concentrations of ethanol (30%, 50%, 70%, and 85%). For each concentration of ethanol, samples were first resuspended in 400 µl of ethanol, incubated at RT for 5 min, and spun at 5000 x g for 2 min. These steps were repeated a second time, though the incubation at RT was extended to 10 min. After the last centrifugation, sample pellets were resuspended in 400 µl ethanol 99.9%, incubated at RT for 10 min, and spun at 5000 x g for 2 min. This was repeated two more times. Before the last wash, samples were placed into a pyramidal beam capsule. Afterwards, each sample was resuspended in 400 µl propylene oxide (Sigma-Aldrich), incubated at RT for 5 min, and spun at 5000 x g for 2 min. This was repeated 3 more times. Hence, each sample was resuspended in 400µl of a 75%-25% propylene oxide/resin mixture and incubated at RT for 15 min prior to being spun at 5000 x g for 2 min. These steps were repeated with 50%-50% and then 25%-75% propylene oxide/resin mixtures. After the last centrifugation, 100% of pure epoxy resin was gently added to each sample pellet and incubated overnight at 37°C. Resin was dried by incubation at 37°C overnight, then at 45°C for 1 day, and eventually at 60°C for three days. Ultrathin sections were cut with a DiATOME ultra 45° diamond knife, deposited into a copper grid and then viewed on a TECNAi 10 transmission electron microscope (PHILIPS) equipped with a megaview CCD camera of 1024×1024 pixel resolution (Olympus) operating at an acceleration voltage of 80 kV. Images were taken via Soft Imaging System (Olympus) from three independent experiments.

#### Negative staining of OMVs for TEM

A previously described protocol was used [94], though with some modifications. Shortly, carbon-coated copper grids for TEM were negatively charged and hydrophilized using a Q150T S/E/ES device (Quorum Technologies) following the manufacturer’s instructions. The day after, 3 µl of bacterial OMVs were deposited onto a negatively charged and hydrophilized grid, incubated at RT for 3 min and then quickly wash with MilliQ water. The grid was then deposited for 3 min at RT onto 1% aqueous uranyl acetate (Sigma-Aldrich). Excess liquid was gently removed, and grids were allowed to air dry. Samples were views as described above. A total of *n*=102 OMVs from three independent experiments were photographed and their diameter was measured by ImageJ.

### AlphaFold2 structural predictions

When available, the structural models were retrieved from the AlphaFold Protein Structure Database (https://alphafold.ebi.ac.uk/). For TamB and TamB2 from *F. johnsoniae* and TamB from *C. canimorsus*, the structures of the last 1000 C-terminal residues were generated using the AlphaFold2 tool available in the ColabFold platform [95]. The full-length structures of (2xStrep)TamB and (3xFLAG)TamL from *F. johnsoniae*, were instead obtained via the AlphaFold2 tool available in the COSMIC² platform [96]. From the same platform, the AlphaFold Multimer tool was also used to predict TamL-TamB complex from *F. johnsoniae* and *C. canimorsus*.

When present, the residues corresponding to the N-terminal signal peptide (SP) were omitted for the analysis. The best-scored models were used for analysis and displayed in a cartoon representation using PyMOL Molecular Graphics Systems Version 2.5.2 (Schrödinger, LCC).

### Conservation of TamL and TamB proteins in Bacteroidetes

A previously published protocol was used [58]. Briefly, A DELTA-BLAST sequence similarity search was conducted on the 30 Bacteroidetes species using the TamA, - L, -B protein sequences of *F. johnsoniae* ATCC 17061 UW101 as queries: TamA (WP_012022494.1), TamL (WP_012023543.1), TamB (WP_012026557.1), TamL2 (WP_012023976.1) and TamB2 (WP_012023975.1). The genomic neighborhood of all hits with an E value ≤ 0.001 was inspected using MaGe [97]. The structure of the corresponding proteins was predicted using AlphaFold server (with default parameters) [98] and compared with the corresponding structures of the of *F. johnsoniae* initial queries by manual inspection. The phylogenetic tree based on NCBI taxonomy was generated via phyloT [99].

### Statistical analyses

Data are presented as Mean ± standard deviation (SD) from at least three independent experiments. All the statistical analyses were performed using GraphPad Prism 8 software.

For the analysis of the outer membrane and OMVs proteomes, a *t*-test (without correction) was performed to compare the means from samples obtained in permissive (+ IPTG) and non-permissive (- IPTG) conditions of growth. When protein peptides could be detected in only one of the two conditions (either + or - IPTG) but not in the other one, a representative value of |64| was used. The Fold Change (FC) was derived using the permissive condition (+ IPTG) as a reference. Proteins exhibiting a FC ≥ |1.5| and a *p*-value of <0.01 were considered as significant.

## Data availability

The mass spectrometry proteomics data have been deposited to the ProteomeXchange Consortium via the PRIDE [100] partner repository with the dataset identifier PXD056806 and 10.6019/PXD056806.

## Acknowledgements.

We are grateful to Isabelle Hamer (UNamur) for providing training and support for ultracentrifuge. We thank Catherine Demazy (UNamur) for help on samples preparation for proteomics. We are grateful to the Electron microscopy facility of UNamur. This research has been funded by the Incentive Grant for Scientific Research (MIS F.4533.20F) from the Fonds de la Recherche Scientifique-Fonds National de la Recherche Scientifique (FRS-FNRS, http://www.fnrs.be) to F. Renzi. F. Renzi is a research associate of the FRS-FNRS.

## Author Contributions

**Conceptualization:** F.G. and F.R.

**Data Curation**: F.G., M.D., R.D. and F.R.

**Formal Analysis**: F.G., M.D. and F.R.

**Funding acquisition**: F.R.

**Investigation**: F.G, T.D.M., V-G.M.A., F.L., R.D, J.D., Z.G. and L.L.

**Methodology**: F.G., F.L. and F.R.

**Project Administration**: F.G. and F.R.

**Resources**: F.R.

**Supervision**: F.G. and F.R.

**Validation**: F.G. and F.R.

**Visualization**: F.G. and F.R.

**Writing – original draft**: F.G. and F.R.

**Writing – review & editing**: F.G., F.L., C.S. and F.R.

## Notes

### Competing Interest Statement

The authors have declared no competing interest.

## References

1. Silhavy TJ, Kahne D, Walker S. The bacterial cell envelope. Cold Spring Harb Perspect Biol. 2010;2: a000414. doi:10.1101/cshperspect.a000414

2. Konovalova A, Kahne DE, Silhavy TJ. Outer Membrane Biogenesis. Annu Rev Microbiol. 2017;71: 539–556. doi:10.1146/annurev-micro-090816-093754

3. Guest RL, Silhavy TJ. Cracking outer membrane biogenesis. Biochim Biophys Acta - Mol Cell Res. 2023;1870: 119405. doi:10.1016/j.bbamcr.2022.119405

4. Okuda S, Tokuda H. Lipoprotein sorting in bacteria. Annu Rev Microbiol. 2011;65: 239–259. doi:10.1146/annurev-micro-090110-102859

5. Zückert WR. Secretion of Bacterial Lipoproteins: Through the Cytoplasmic Membrane, the Periplasm and Beyond. Biochim Biophys Acta - Mol Cell Res. 2014;1843: 1509–1516. doi:10.1016/j.bbamcr.2014.04.022

6. Calmettes C, Judd A, Moraes TF. Structural aspects of bacterial outer membrane protein assembly. Adv Exp Med Biol. 2015;883: 225–270. doi:10.1007/978-3-319-23603-2_14

7. Gatsos X, Perry AJ, Anwari K, Dolezal P, Wolynec PP, Likić VA, et al. Protein secretion and outer membrane assembly in Alphaproteobacteria. FEMS Microbiol Rev. 2008;32: 995–1009. doi:10.1111/j.1574-6976.2008.00130.x

8. Simmerman RF, Dave AM, Bruce BD. Structure and Function of POTRA Domains of Omp85/TPS Superfamily. International Review of Cell and Molecular Biology. Elsevier Inc.; 2014. doi:10.1016/B978-0-12-800097-7.00001-4

9. Gu Y, Li H, Dong H, Zeng Y, Zhang Z, Paterson NG, et al. Structural basis of outer membrane protein insertion by the BAM complex. Nature. 2016;531: 64–69. doi:10.1038/nature17199

10. Goh KJ, Stubenrauch CJ, Lithgow T. The TAM, a Translocation and Assembly Module for protein assembly and potential conduit for phospholipid transfer. EMBO Rep. 2024;25: 1711–1720. doi:10.1038/s44319-024-00111-y

11. Stubenrauch CJ, Lithgow T. The TAM: A Translocation and Assembly Module of the β-Barrel Assembly Machinery in Bacterial Outer Membranes. EcoSal Plus. 2019;8: 1–7. doi:10.1128/ecosalplus.esp-0036-2018

12. Albenne C, Ieva R. Job contenders: roles of the β-barrel assembly machinery and the translocation and assembly module in autotransporter secretion. Mol Microbiol. 2017;106: 505–517. doi:10.1111/mmi.13832

13. Selkrig J, Mosbahi K, Webb CT, Belousoff MJ, Perry AJ, Wells TJ, et al. Discovery of an archetypal protein transport system in bacterial outer membranes. Nat Struct Mol Biol. 2012;19: 506–510. doi:10.1038/nsmb.2261

14. Selkrig J, Belousoff MJ, Headey SJ, Heinz E, Shiota T, Shen HH, et al. Conserved features in TamA enable interaction with TamB to drive the activity of the translocation and assembly module. Sci Rep. 2015;5: 1–12. doi:10.1038/srep12905

15. Gruss F, Zähringer F, Jakob RP, Burmann BM, Hiller S, Maier T. The structural basis of autotransporter translocation by TamA. Nat Struct Mol Biol. 2013;20: 1318–1320. doi:10.1038/nsmb.2689

16. Mellouk A, Jaouen P, Ruel L-J, Lê M, Martini C, Moraes TF, et al. POTRA domains of the TamA insertase interact with the outer membrane and modulate membrane properties. Proc Natl Acad Sci. 2024;121: e2402543121. doi:10.1073/pnas.2402543121

17. Josts I, Stubenrauch CJ, Vadlamani G, Mosbahi K, Walker D, Lithgow T, et al. The Structure of a Conserved Domain of TamB Reveals a Hydrophobic β Taco Fold. Structure. 2017;25: 1898–1906.e5. doi:10.1016/j.str.2017.10.002

18. Kumar S, Ruiz N. Bacterial AsmA-Like Proteins: Bridging the Gap in Intermembrane Phospholipid Transport. Contact. 2023;6: 1–9. doi:10.1177/25152564231185931

19. Neuman SD, Levine TP, Bashirullah A. A novel superfamily of bridge-like lipid transfer proteins. Trends Cell Biol. 2022;32: 962–974. doi:10.1016/j.tcb.2022.03.011

20. Melia TJ, Reinisch KM. A possible role for VPS13-family proteins in bulk lipid transfer, membrane expansion and organelle biogenesis. J Cell Sci. 2022;135. doi:10.1242/jcs.259357

21. McEwan DG, Ryan KM. ATG2 and VPS13 proteins: molecular highways transporting lipids to drive membrane expansion and organelle communication. FEBS J. 2022;289: 7113–7127. doi:10.1111/febs.16280

22. Ruiz N, Davis RM, Kumar S. YhdP, TamB, and YdbH Are Redundant but Essential for Growth and Lipid Homeostasis of the Gram-Negative Outer Membrane. MBio. 2021;12. doi:10.1128/mBio.02714-21

23. Douglass M V., McLean AB, Trent MS. Absence of YhdP, TamB, and YdbH leads to defects in glycerophospholipid transport and cell morphology in Gram-negative bacteria. PLoS Genet. 2022;18: 1–21. doi:10.1371/journal.pgen.1010096

24. Cooper BF, Clark R, Kudhail A, Bhabha G, Ekiert DC, Khalid S, et al. Phospholipid transport to the bacterial outer membrane through an envelope-spanning bridge. bioRxiv [Preprint]. 2023. doi:10.1101/2023.10.05.561070

25. Rai AK, Sawasato K, Bennett HC, Kozlova A, Sparagna GC, Bogdanov M, et al. Genetic evidence for functional diversification of gram-negative intermembrane phospholipid transporters. PLoS Genet. 2024;20: 1–26. doi:10.1371/journal.pgen.1011335

26. Mcdonnell RT, Patel N, Wehrspan ZJ, Elcock AH. Atomic Models of All Major Trans-Envelope Complexes Involved in Lipid Trafficking in Escherichia Coli Constructed Using a Combination of AlphaFold2, AF2Complex, and Membrane Morphing Simulations Keywords. bioRxiv [Preprint]. 2023. doi:10.1101/2023.04.28.538765

27. Kumar S, Davis RM, Ruiz N. YdbH and YnbE form an intermembrane bridge to maintain lipid homeostasis in the outer membrane of Escherichia coli. Proc Natl Acad Sci. 2024;121. 10.1073/pnas.2321512121

28. Heinz E, Selkrig J, Belousoff MJ, Lithgow T. Evolution of the translocation and assembly module (TAM). Genome Biol Evol. 2015;7: 1628–1643. doi:10.1093/gbe/evv097

29. Heinz E, Lithgow T. A comprehensive analysis of the Omp85/TpsB protein superfamily structural diversity, taxonomic occurrence, and evolution. Front Microbiol. 2014;5: 1–13. doi:10.3389/fmicb.2014.00370

30. Foley MH, Cockburn DW, Koropatkin NM. The Sus operon: a model system for starch uptake by the human gut Bacteroidetes. Cell Mol Life Sci. 2016;73: 2603–2617. doi:10.1007/s00018-016-2242-x

31. Manfredi P, Renzi F, Mally M, Sauteur L, Schmaler M, Moes S, et al. The genome and surface proteome of Capnocytophaga canimorsus reveal a key role of glycan foraging systems in host glycoproteins deglycosylation. Mol Microbiol. 2011;81: 1050–1060. doi:10.1111/j.1365-2958.2011.07750.x

32. Lauber F, Cornelis GR, Renzi F. Identification of a new lipoprotein export signal in gram-negative bacteria. MBio. 2016;7. doi:10.1128/mBio.01232-16

33. Pitta TP, Leadbetter ER, Godchaux W. Increase of ornithine amino lipid content in a sulfonolipid-deficient mutant of Cytophaga johnsonae. J Bacteriol. 1989;171: 952–957. doi:10.1128/jb.171.2.952-957.1989

34. Vences-Guzmán MÁ, Peña-Miller R, Hidalgo-Aguilar NA, Vences-Guzmán ML, Guan Z, Sohlenkamp C. Identification of the Flavobacterium johnsoniae cysteate-fatty acyl transferase required for capnine synthesis and for efficient gliding motility. Environ Microbiol. 2021;23: 2448–2460. doi:10.1111/1462-2920.15445

35. McBride MJ, Xie G, Martens EC, Lapidus A, Henrissat B, Rhodes RG, et al. Novel features of the polysaccharide-digesting gliding bacterium Flavobacterium johnsoniae as revealed by genome sequence analysis. Appl Environ Microbiol. 2009;75: 6864–6875. doi:10.1128/AEM.01495-09

36. Lauber F, Deme JC, Lea SM, Berks BC. Type 9 secretion system structures reveal a new protein transport mechanism. Nature. 2018;564: 77–82. doi:10.1038/s41586-018-0693-y

37. Lauber F, Deme JC, Liu X, Kjær A, Miller HL, Alcock F, et al. Structural insights into the mechanism of protein transport by the Type 9 Secretion System translocon. Nature Microbiology. 2024. doi:10.1038/s41564-024-01644-7

38. McBride MJ, Nakane D. Flavobacterium gliding motility and the type IX secretion system. Curr Opin Microbiol. 2015;28: 72–77. doi:10.1016/j.mib.2015.07.016

39. McBride MJ. Bacteroidetes gliding motility and the type IX secretion system. Protein Secret Bact. 2019;7: 363–374. doi:10.1128/9781683670285.ch29

40. Johnston JJ, Shrivastava A, McBride MJ. Untangling Flavobacterium johnsoniae Gliding Motility and Protein Secretion. J Bacteriol. 2017;20: 2.

41. Mally M, Paroz C, Shin H, Meyer S, Soussoula L V., Schmiediger U, et al. Prevalence of Capnocytophaga canimorsus in dogs and occurrence of potential virulence factors. Microbes Infect. 2009;11: 509–514. doi:10.1016/j.micinf.2009.02.005

42. Renzi F, Dol M, Raymackers A, Manfredi P, Cornelis GR. Only a subset of C. canimorsus strains is dangerous for humans. Emerg Microbes Infect. 2015;4. doi:10.1038/emi.2015.48

43. Boratyn GM, Schäffer AA, Agarwala R, Altschul SF, Lipman DJ, Madden TL. Domain enhanced lookup time accelerated BLAST. Biol Direct. 2012;7: 1–14. doi:10.1186/1745-6150-7-12

44. Teufel F, Almagro Armenteros JJ, Johansen AR, Gíslason MH, Pihl SI, Tsirigos KD, et al. SignalP 6.0 predicts all five types of signal peptides using protein language models. Nat Biotechnol. 2022;40: 1023–1025. doi:10.1038/s41587-021-01156-3

45. Sposato D, Mercolino J, Torrini L, Sperandeo P, Lucidi M, Alegiani R, et al. Redundant essentiality of AsmA-like proteins in *Pseudomonas aeruginosa*. mSphere. 2024;9: e00677–23. doi:10.1128/msphere.00677-23

46. Wu T, Malinverni J, Ruiz N, Kim S, Silhavy TJ, Kahne D. Identification of a multicomponent complex required for outer membrane biogenesis in Escherichia coli. Cell. 2005;121: 235–245. doi:10.1016/j.cell.2005.02.015

47. Werner J, Misra R. YaeT (Omp85) affects the assembly of lipid-dependent and lipid-independent outer membrane proteins of Escherichia coli. Mol Microbiol. 2005;57: 1450–1459. doi:10.1111/j.1365-2958.2005.04775.x

48. Acharya Y, Haldar J. Upgrading the Antibiotic Arsenal Against Gram-Positive Bacteria: Chemical Modifications of Vancomycin. In: Saha T, Deb Adhikari M, Tiwary BK, editors. Alternatives to Antibiotics: Recent Trends and Future Prospects. Singapore: Springer Nature Singapore; 2022. pp. 199–222. doi:10.1007/978-981-19-1854-4_8

49. Mohapatra SS, Dwibedy SK, Padhy I. Polymyxins, the last-resort antibiotics: Mode of action, resistance emergence, and potential solutions. J Biosci. 2021;46. doi:10.1007/s12038-021-00209-8

50. Bhairi SM, Mohan C. A Guide to the properties and uses of Detergents in biology and biochemistry. Calbiochem. 2001; 41. Available: http://wolfson.huji.ac.il/purification/PDF/detergents/CALBIOCHEM-DetergentsIV.pdf

51. Grimm J, Shi H, Wang W, Mitchell AM, Wingreen NS, Huang KC, et al. The inner membrane protein YhdP modulates the rate of anterograde phospholipid flow in Escherichia coli. Proc Natl Acad Sci U S A. 2020;117: 26907–26914. doi:10.1073/pnas.2015556117

52. Sutterlin HA, Shi H, May KL, Miguel A, Khare S, Huang KC, et al. Disruption of lipid homeostasis in the Gram-negative cell envelope activates a novel cell death pathway. Proc Natl Acad Sci U S A. 2016;113: E1565–E1574. doi:10.1073/pnas.1601375113

53. Sato K, Naito M, Yukitake H, Hirakawa H, Shoji M, McBride MJ, et al. A protein secretion system linked to bacteroidete gliding motility and pathogenesis. Proc Natl Acad Sci U S A. 2010;107: 276–281. doi:10.1073/pnas.0912010107

54. Noinaj N, Guillier M, Barnard TJ, Buchanan SK. TonB-dependent transporters: Regulation, structure, and function. Annu Rev Microbiol. 2010;64: 43–60. doi:10.1146/annurev.micro.112408.134247

55. Du D, Wang-Kan X, Neuberger A, van Veen HW, Pos KM, Piddock LJ V, et al. Multidrug efflux pumps: structure, function and regulation. Nat Rev Microbiol. 2018;16: 523–539. doi:10.1038/s41579-018-0048-6

56. Tan Z, Black W, Yoon JM, Shanks J V., Jarboe LR. Improving Escherichia coli membrane integrity and fatty acid production by expression tuning of FadL and OmpF. Microb Cell Fact. 2017;16: 1–15. doi:10.1186/s12934-017-0650-8

57. Grondin JM, Tamura K, Déjean G, Abbott DW, Brumer H. Polysaccharide utilization loci: Fueling microbial communities. J Bacteriol. 2017;199. doi:10.1128/JB.00860-16

58. De Smet T, Baland E, Giovannercole F, Mignon J, Lizen L, Dugauquier R, et al. LolA and LolB are conserved in Bacteroidetes and are crucial for gliding motility and Type IX secretion. bioRxiv [Preprint]. 2024; 1–36. 10.1101/2024.09.13.612696

59. Rhodes RG, Samarasam MN, Van Groll EJ, McBride MJ. Mutations in Flavobacterium johnsoniae sprE result in defects in gliding motility and protein secretion. J Bacteriol. 2011;193: 5322–5327. doi:10.1128/JB.05480-11

60. Vences-Guzmán MÁ, Guan Z, Escobedo-Hinojosa WI, Bermúdez-Barrientos JR, Geiger O, Sohlenkamp C. Discovery of a bifunctional acyltransferase responsible for ornithine lipid synthesis in Serratia proteamaculans. Environ Microbiol. 2015;17: 1487–1496. doi:10.1111/1462-2920.12562

61. Evans R, O’Neill M, Pritzel A, Antropova N, Senior A, Green T, et al. Protein complex prediction with AlphaFold-Multimer. bioRxiv. 2022; 2021.10.04.463034. doi:10.1101/2021.10.04.463034

62. Noinaj N, Gumbart JC, Buchanan SK. The β-barrel assembly machinery in motion. Nat Rev Microbiol. 2017;15: 197–204. doi:10.1038/nrmicro.2016.191

63. Doyle MT, Bernstein HD. Bacterial outer membrane proteins assemble via asymmetric interactions with the BamA β-barrel. Nat Commun. 2019;10: 1–13. doi:10.1038/s41467-019-11230-9

64. Tomasek D, Rawson S, Lee J, Wzorek JS, Harrison SC, Li Z, et al. Structure of a nascent membrane protein as it folds on the BAM complex. Nature. 2020;583: 473–478. doi:10.1038/s41586-020-2370-1

65. Shen C, Chang S, Luo Q, Chan KC, Zhang Z, Luo B, et al. Structural basis of BAM-mediated outer membrane β-barrel protein assembly. Nature. 2023;617: 185–193. doi:10.1038/s41586-023-05988-8

66. Butler T. Capnocytophaga canimorsus: an emerging cause of sepsis, meningitis, and post-splenectomy infection after dog bites. Eur J Clin Microbiol Infect Dis Off Publ Eur Soc Clin Microbiol. 2015;34: 1271–1280. doi:10.1007/s10096-015-2360-7

67. Li MF, Jia BB, Sun YY, Sun L. The Translocation and Assembly Module (TAM) of Edwardsiella tarda Is Essential for Stress Resistance and Host Infection. Front Microbiol. 2020;11: 1–12. doi:10.3389/fmicb.2020.01743

68. Bialer MG, Ruiz-Ranwez V, Sycz G, Estein SM, Russo DM, Altabe S, et al. MapB, the Brucella suis TamB homologue, is involved in cell envelope biogenesis, cell division and virulence. Sci Rep. 2019;9: 1–18. doi:10.1038/s41598-018-37668-3

69. Wang X, Nyenhuis SB, Bernstein HD. The translocation assembly module (TAM) catalyzes the assembly of bacterial outer membrane proteins in vitro. Nat Commun. 2024;15: 1–15. doi:10.1038/s41467-024-51628-8

70. Shoji M, Shibata S, Sueyoshi T, Naito M, Nakayama K. Biogenesis of Type V pili. Microbiol Immunol. 2020;64: 643–656. doi:10.1111/1348-0421.12838

71. Martorana AM, Motta S, Di Silvestre D, Falchi F, Dehò G, Mauri P, et al. Dissecting Escherichia coli outer membrane biogenesis using differential proteomics. PLoS One. 2014;9. doi:10.1371/journal.pone.0100941

72. Asmar AT, Collet JF. Lpp, the Braun lipoprotein, turns 50—major achievements and remaining issues. FEMS Microbiol Lett. 2018;365: 1–8. doi:10.1093/femsle/fny199

73. Liao CT, Li CE, Chang HC, Hsu CH, Chiang YC, Hsiao YM. The lolB gene in Xanthomonas campestris pv. campestris is required for bacterial attachment, stress tolerance, and virulence. BMC Microbiol. 2022;22: 1–13. doi:10.1186/s12866-021-02416-7

74. Rooke JL, Icke C, Wells TJ, Rossiter AE, Browning DF, Morris FC, et al. BamA and BamD Are Essential for the Secretion of Trimeric Autotransporter Adhesins. Front Microbiol. 2021;12: 1–10. doi:10.3389/fmicb.2021.628879

75. Malinverni JC, Werner J, Kim S, Sklar JG, Kahne D, Misra R, et al. YfiO stabilizes the YaeT complex and is essential for outer membrane protein assembly in Escherichia coli. Mol Microbiol. 2006;61: 151–164. doi:10.1111/j.1365-2958.2006.05211.x

76. Mikheyeva I V., Sun J, Huang KC, Silhavy TJ. Mechanism of outer membrane destabilization by global reduction of protein content. Nat Commun. 2023;14: 1–9. doi:10.1038/s41467-023-40396-6

77. Sartorio MG, Pardue EJ, Scott NE, Feldman MF. Human gut bacteria tailor extracellular vesicle cargo for the breakdown of diet- and host-derived glycans. Proc Natl Acad Sci. 2023;120: e2306314120. doi:10.1073/pnas.2306314120

78. Valguarnera E, Scott NE, Azimzadeh P, Feldman MF. Surface Exposure and Packing of Lipoproteins into Outer Membrane Vesicles Are Coupled Processes in Bacteroides. mSphere. 2018;3: 1–14. doi:10.1128/msphere.00559-18

79. Mamou G, Corona F, Cohen-Khait R, Housden NG, Yeung V, Sun D, et al. Peptidoglycan maturation controls outer membrane protein assembly. Nature. 2022;606: 953–959. doi:10.1038/s41586-022-04834-7

80. McBride MJ, Kempf MJ. Development of techniques for the genetic manipulation of the gliding bacterium Cytophaga johnsonae. J Bacteriol. 1996;178: 583–590. doi:10.1128/jb.178.3.583-590.1996

81. Jun L, J. MM, Sriram S. Cell Surface Filaments of the Gliding Bacterium Flavobacterium johnsoniae Revealed by Cryo-Electron Tomography. J Bacteriol. 2007;189: 7503–7506. doi:10.1128/jb.00957-07

82. Gibson DG, Young L, Chuang RY, Venter JC, Hutchison CA, Smith HO. Enzymatic assembly of DNA molecules up to several hundred kilobases. Nat Methods. 2009;6: 343–345. doi:10.1038/nmeth.1318

83. Parker AC, Smith CJ. Development of an IPTG inducible expression vector adapted for Bacteroides fragilis. Plasmid. 2012;68: 86–92. doi:10.1007/978-981-99-9283-6_2167

84. Hunnicutt DW, McBride MJ. Cloning and characterization of the Flavobacterium johnsoniae gliding-motility genes gldB and gldC. J Bacteriol. 2000;182: 911–918. doi:10.1128/JB.182.4.911-918.2000

85. Kasahara M, Anraku Y. Succinate dehydrogenase of Escherichia coli membrane vesicles. Activation and properties of the enzyme. J Biochem. 1974;76: 959–966.

86. Rabilloud T, Carpentier G, Tarroux P. Improvement and simplification of low-background silver staining of proteins by using sodium dithionite. Electrophoresis. 1988;9: 288–291. doi:10.1002/elps.1150090608

87. Meier F, Brunner A-D, Koch S, Koch H, Lubeck M, Krause M, et al. Online Parallel Accumulation-Serial Fragmentation (PASEF) with a Novel Trapped Ion Mobility Mass Spectrometer. Mol Cell Proteomics. 2018;17: 2534–2545. doi:10.1074/mcp.TIR118.000900

88. Yu NY, Wagner JR, Laird MR, Melli G, Rey S, Lo R, et al. PSORTb 3.0: Improved protein subcellular localization prediction with refined localization subcategories and predictive capabilities for all prokaryotes. Bioinformatics. 2010;26: 1608–1615. doi:10.1093/bioinformatics/btq249

89. Drula E, Garron ML, Dogan S, Lombard V, Henrissat B, Terrapon N. The carbohydrate-active enzyme database: Functions and literature. Nucleic Acids Res. 2022;50: D571–D577. doi:10.1093/nar/gkab1045

90. Bligh, E.G. and Dyer WJ. Canadian Journal of Biochemistry and Physiology. Can J Biochem Physiol. 1959;37.

91. Okino N, Ito M. Thin-layer chromatography (TLC) of glycolipids. In: Nishihara S, Angata K, Aoki-Kinoshita KF, Hirabayashi J, editors. Saitama (JP); 2021.

92. Joyce LR, Manzer HS, da C. Mendonça J, Villarreal R, Nagao PE, Doran KS, et al. Identification of a novel cationic glycolipid in Streptococcus agalactiae that contributes to brain entry and meningitis. PLOS Biol. 2022;20: e3001555. Available: 10.1371/journal.pbio.3001555

93. Wenzel M, Dekker MP, Wang B, Burggraaf MJ, Bitter W, van Weering JRT, et al. A flat embedding method for transmission electron microscopy reveals an unknown mechanism of tetracycline. Commun Biol. 2021;4: 306. doi:10.1038/s42003-021-01809-8

94. Pardue EJ, Sartorio MG, Jana B, Scott NE, Beatty WL, Ortiz-Marquez JC, et al. Dual membrane-spanning anti-sigma factors regulate vesiculation in Bacteroides thetaiotaomicron. Proc Natl Acad Sci. 2024;121: e2321910121. doi:10.1073/pnas.2321910121

95. Mirdita M, Schütze K, Moriwaki Y, Heo L, Ovchinnikov S, Steinegger M. ColabFold: making protein folding accessible to all. Nat Methods. 2022;19: 679–682. doi:10.1038/s41592-022-01488-1

96. Cianfrocco MA, Wong-Barnum M, Youn C, Wagner R, Leschziner A. COSMIC2: A Science Gateway for Cryo-Electron Microscopy Structure Determination. Practice and Experience in Advanced Research Computing 2017: Sustainability, Success and Impact. New York, NY, USA: Association for Computing Machinery; 2017. doi:10.1145/3093338.3093390

97. Vallenet D, Labarre L, Rouy Z, Barbe V, Bocs S, Cruveiller S, et al. MaGe: a microbial genome annotation system supported by synteny results. Nucleic Acids Res. 2006;34: 53–65. doi:10.1093/nar/gkj406

98. Jumper J, Evans R, Pritzel A, Green T, Figurnov M, Ronneberger O, et al. Highly accurate protein structure prediction with AlphaFold. Nature. 2021;596: 583–589. doi:10.1038/s41586-021-03819-2

99. Letunic I, Doerks T, Bork P. SMART 7: recent updates to the protein domain annotation resource. Nucleic Acids Res. 2012;40: D302–5. doi:10.1093/nar/gkr931

100. Perez-Riverol Y, Bai J, Bandla C, García-Seisdedos D, Hewapathirana S, Kamatchinathan S, et al. The PRIDE database resources in 2022: a hub for mass spectrometry-based proteomics evidences. Nucleic Acids Res. 2022;50: D543–D552. doi:10.1093/nar/gkab1038

