## Supplementary Figures and Tables for "TamL is a key player of the outer membrane homeostasis in Bacteroidetes"

### Contents

**Supplementary Figure S1:** *tamA, tamB2*, and *tamL2* deletion has no effect on bacterial growth.

**Supplementary Figure S2:** Deletion of *tamA* and *tamL2* in TamL-depleted cells has no effect on cell viability.

**Supplementary Figure S3:** Total protein content (in mg) of whole-cell lysates and membrane fractions of *F. johnsoniae* cells grown in permissive and non-permissive conditions.

**Supplementary Figure S4.** Separation of membrane lipids from *F. johnsoniae cells* grown in permissive (+ IPTG) and non-permissive (- IPTG) conditions by two-dimensional thin-layer chromatography (2D-TLC) and visualized via primuline staining.

**Supplementary Figure S5.** Growth curves of *F. johnsoniae* wild-type (WT), sulfonolipids and ornithine lipids mutants (Δ*fjoh_2419* and Δ*fjoh_0833*, respectively) and of the double mutant (Δ*fjoh_0833*Δ*fjoh_2419*) in CYE (left) and MM (right).

**Supplementary Figure S6.** Structural model of the TamL-TamB interaction in *F. johnsoniae* (A) and *C. canimorsus* (B) predicted via AlphaFold Multimer.

**Supplementary Table S1**. Proteins of outer membrane fractions, isolated from cells grown in permissive (+ IPTG) and non-permissive (- IPTG) conditions and sorted in descending order, identified by label-free mass spectrometry whose spectra count is significantly different (FC ≥ |1.5|) between the two conditions.

**Supplementary Table S2.** Proteins of outer membrane vesicles (OMVs), isolated from cells grown in permissive (+ IPTG) and non-permissive (- IPTG) conditions and sorted in descending order, identified by label-free mass spectrometry whose spectra count is significantly different (FC ≥ |1.5|) between the two conditions.

**Supplementary Table S3**. TAM homologs in several Bacteroidetes species.

**Supplementary Table S4**. List of strains used in this study.

**Supplementary Table S5.** List of plasmids used in this study.

**Supplementary Table S6.** List of oligonucleotides used in this study.

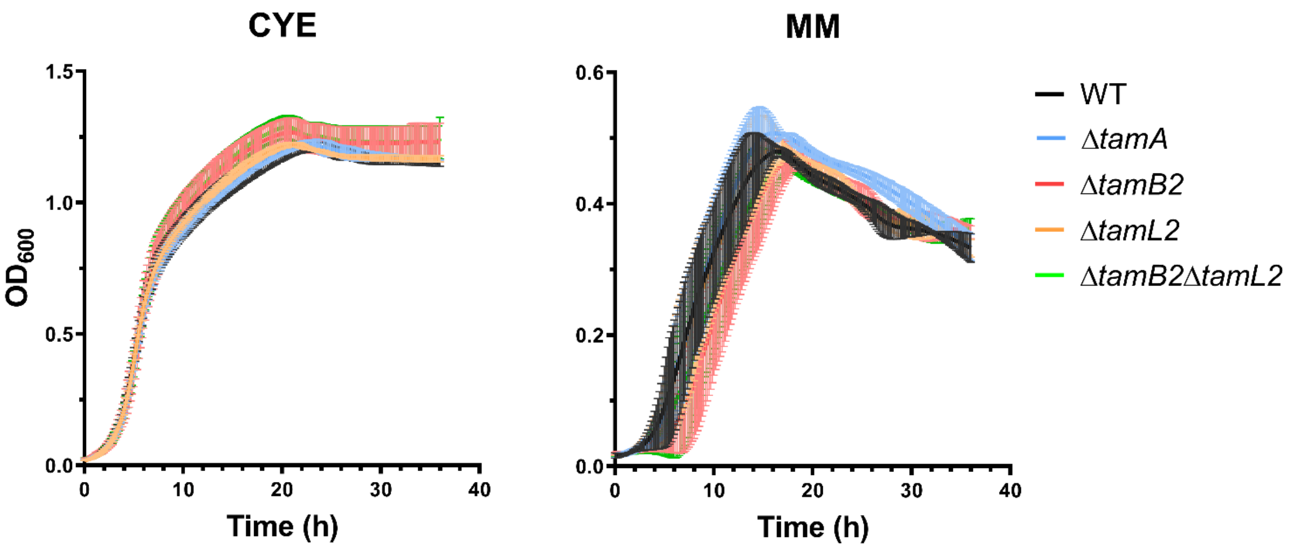

**Supplementary Figure S1: *tamA, tamB2*, and *tamL2* deletion has no effect on bacterial growth.** Cells from overnight cultures (in CYE medium) were freshly inoculated in the same medium or in Motility Medium (MM) in a 96-well plate and incubated at 30°C for 36 hours under constant shaking. Data from three independent experiments are displayed as mean ± standard deviation.

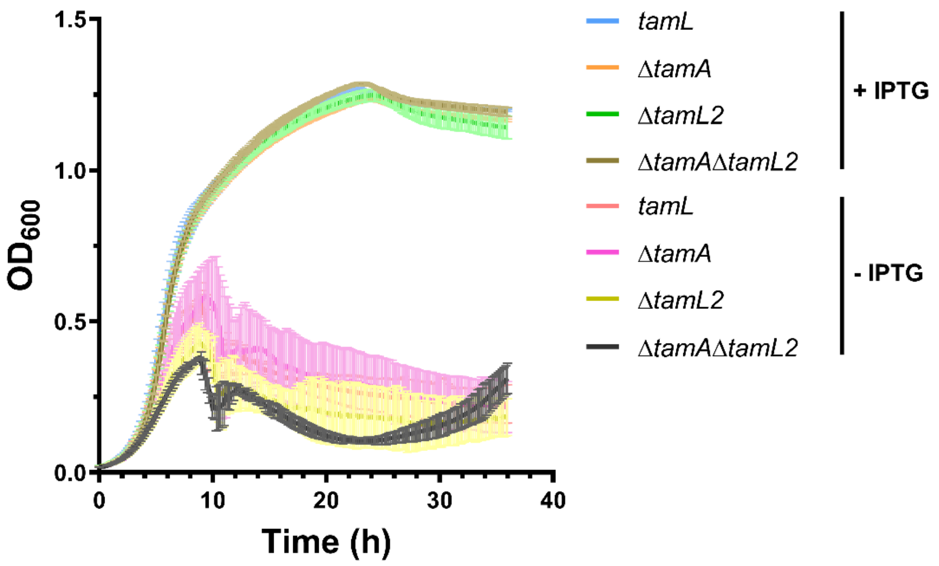

**Supplementary Figure S2: Deletion of *tamA* and *tamL2* in TamL-depleted cells has no effect on cell viability.** Cells from overnight cultures (in CYE medium) grown in permissive conditions (+ IPTG) were $OD_{600}$-normalized, pelleted, washed twice in 1xPBS and freshly inoculated in the same medium with/without IPTG (permissive and non-permissive conditions, respectively) in a 96-well plate and incubated at 30°C for 36 hours under constant shaking. Data from three independent experiments are displayed as mean ± standard deviation.

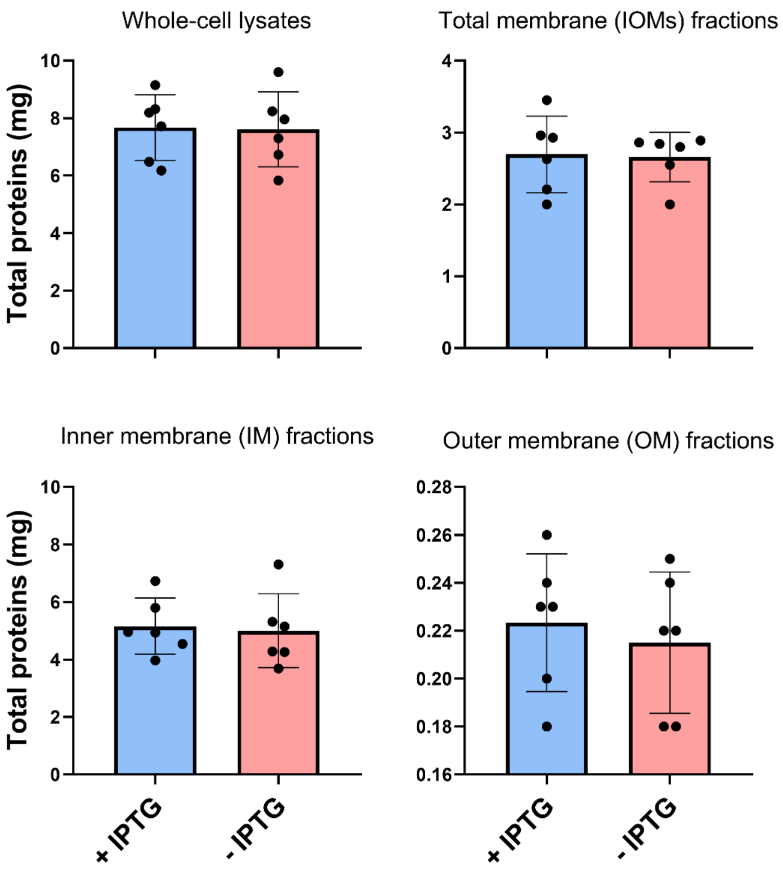

**Supplementary Figure S3: Total protein content (in mg) of whole-cell lysates and membrane fractions of *F. johnsoniae* cells grown in permissive (+ IPTG) and non-permissive (- IPTG) conditions.** The protein concentration of the different fractions was quantified using the Quick StartTM Bradford Protein Assay (Bio-Rad) following the manufacturer`s instructions. Data are displayed as mean ± standard deviation from six independent experiments.

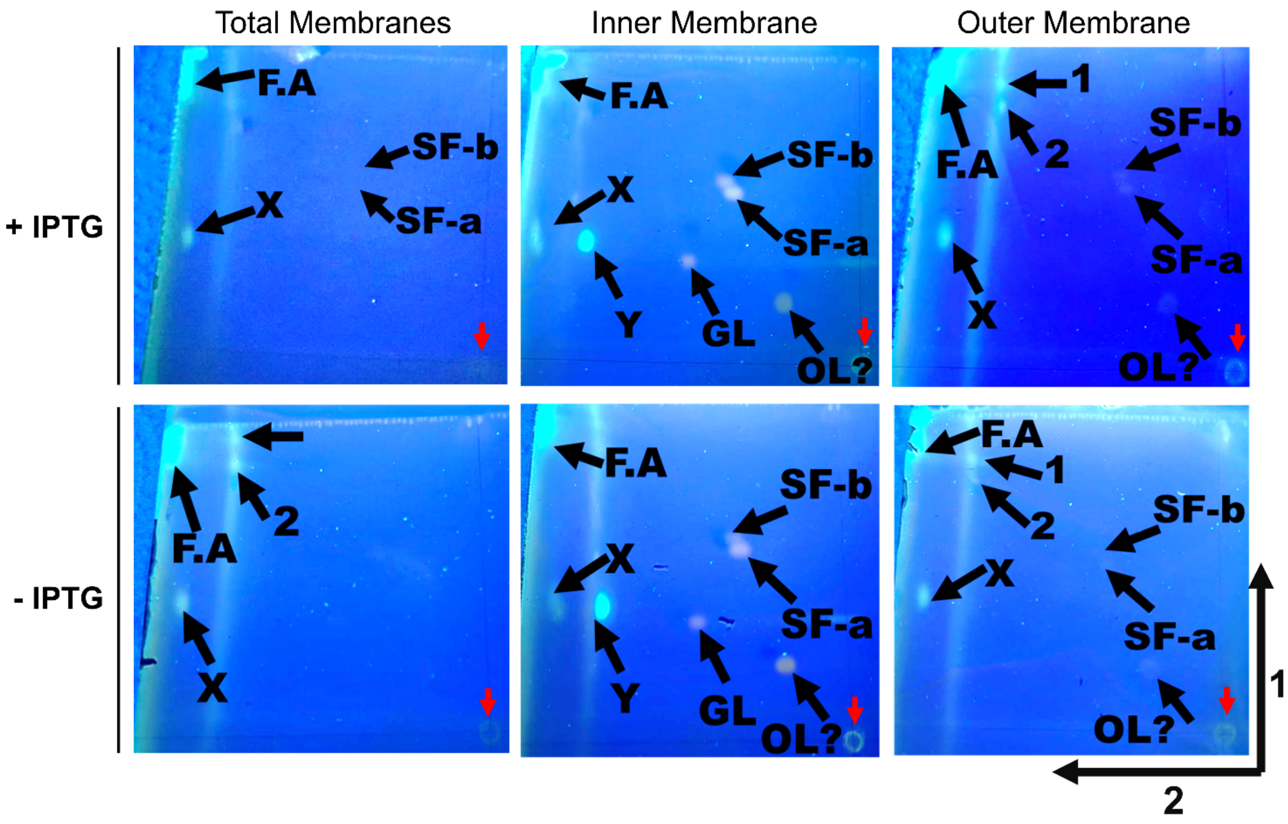

**Supplementary Figure S4. Separation of membrane lipids from *F. johnsoniae cells* grown in permissive (+ IPTG) and non-permissive (- IPTG) conditions by two-dimensional thin-layer chromatography (2D-TLC) and visualized via primuline staining** [1]**.** Lipids are indicated by arrows as: sulfonolipids (SF-a, SF-b), glycine lipids (GL), ornithine lipids (OL), fatty acids (F.A) and unknown lipids (X and Y). Spots were assigned to each lipid based on prior TLC migration performed in the same conditions [2]. Red arrows indicate the migration origins. The black arrows outside the bottom-left panel indicate the two dimensions of migration.

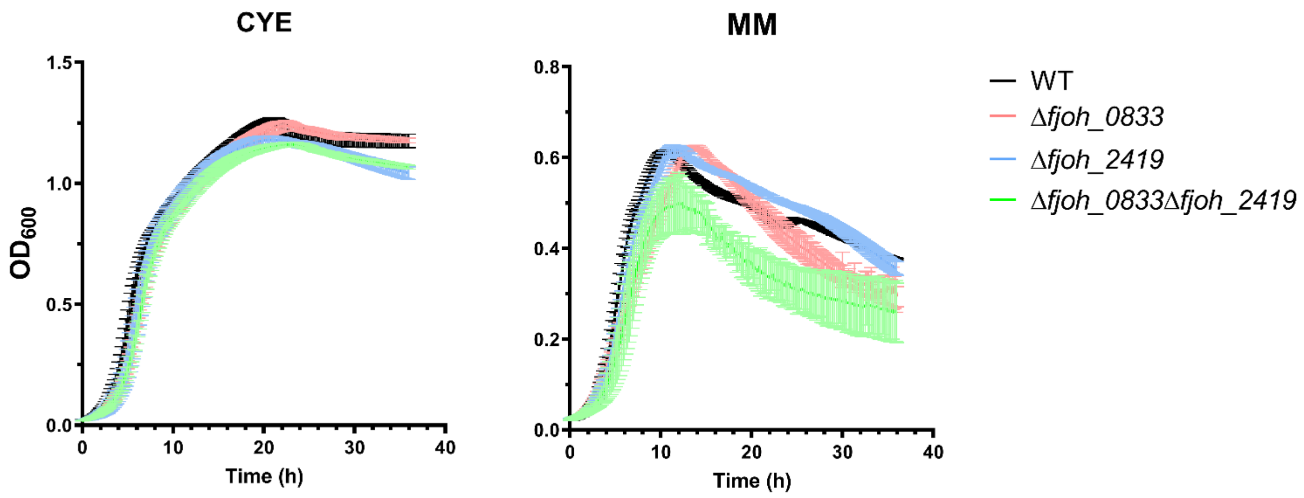

**Supplementary Figure S5. Growth curves of *F. johnsoniae* wild-type (WT), sulfonolipids and ornithine lipids mutants (Δ*fjoh_2419* and Δ*fjoh_0833*, respectively) and of the double mutant (Δ*fjoh_0833*Δ*fjoh_2419*) in CYE (left) and MM (right).** Cells from overnight cultures (in CYE medium) were freshly inoculated in the same medium or in Motility Medium (MM) onto a 96-well plate and incubated at 30°C for 36 hours under constant shaking. Data from three independent experiments are displayed as mean ± standard deviation.

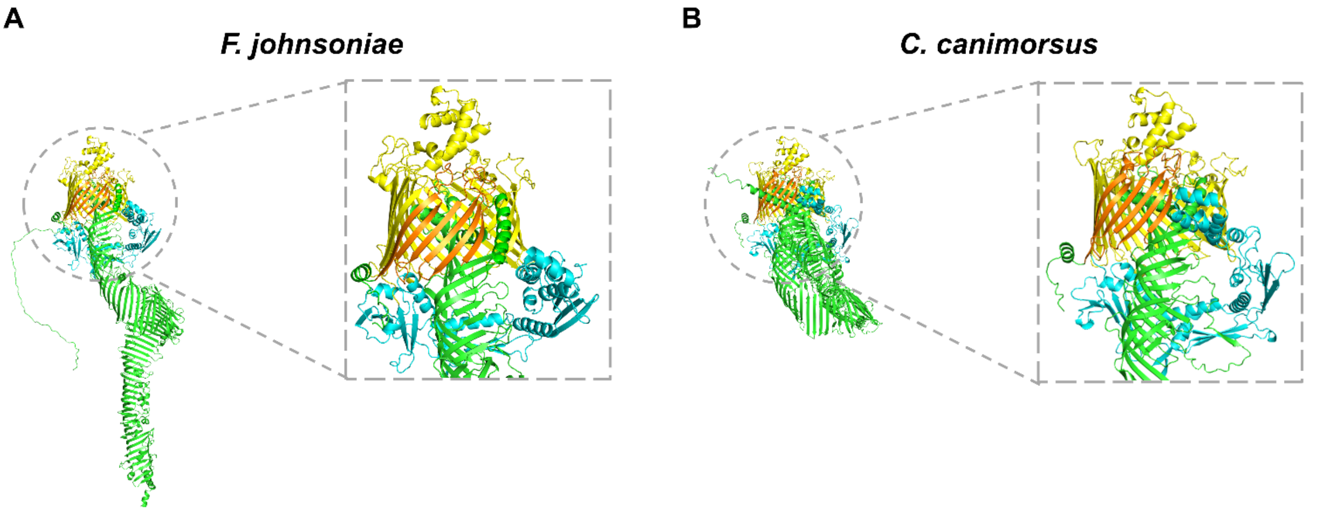

**Supplementary Figure S6. Structural model of the TamL-TamB interaction in *F. johnsoniae* (A) and *C. canimorsus* (B) predicted via AlphaFold Multimer** [3]**.** In (A) and (B), TamB is shown in green, while the three N-terminal POTRA domains and the C-terminal β-barrel domain of TamL are displayed in cyan and yellow, respectively. The pseudosubstrate domain of TamB (residues 1321-1420 in A, residues 1346-1449 in B), which in the model establishes physical interaction with the last β-strand of the β-barrel domain of TamL, is shown in orange. B (inset panel), TamB residues 1-988 are omitted**.**

**Supplementary Table S1. Proteins of outer membrane fractions, isolated from cells grown in permissive (+ IPTG) and non-permissive (- IPTG) conditions and sorted in descending order, identified by label-free mass spectrometry whose spectra count is significantly different (FC ≥ |1.5|) between the two conditions**. Listed are: Protein name; Accession number (as in Uniprot); Signal Peptide (SP) prediction based on SignalP 6.0 [4]; *p*-value (as given by a two-tailed *t*-test); FC: fold change (- IPTG/+ IPTG); Modularity and PULs (as in CAZy database [5]); Protein family (PFAM domain annotation); Gene Ontology (as annotated in MaGe [6]). Empty cells indicate annotation not available. "INF" and "0" were assigned when no peptide was detected in permissive (+ IPTG) or non-permissive (- IPTG) conditions of growth, respectively.

| Protein name | Accession number | Signal peptide | Type | *p*-value | FC | Modularity (CAZy) | PUL (CAZy) | Protein family | Gene Ontology |
|---|---|---|---|---|---|---|---|---|---|
| Fjoh_0402 | A5FMY6 | SPI | Integral/β-barrel protein | 0.0076 | INF | | | | Cell wall/membrane/envelope biogenesis |
| Fjoh_1518 | A5FJR7 | SPI | Integral/β-barrel protein | < 0.00010 | INF | | | Outer membrane protein beta-barrel domain | Cell wall/membrane/envelope biogenesis |
| Fjoh_4284 | A5FBY3 | SPI | Integral/β-barrel protein | 0.00031 | INF | | | Outer membrane efflux protein | Cell wall/membrane/envelope biogenesis; Intracellular trafficking, secretion, and vesicular transport |
| Fjoh_0204 | A5FNH9 | SPI | Integral/β-barrel protein | 0.0065 | INF | | | | Function unknown |
| Fjoh_3920 | A5FCY5 | SPI | Integral/β-barrel protein | < 0.00010 | INF | | | Carboxypeptidase regulatory-like domain; TonB dependent receptor | Inorganic ion transport and metabolism |
| Fjoh_5035 | A5F9T1 | SPI | Integral/β-barrel protein | 0.0014 | INF | SusC | | von Willebrand factor type A domain;<br>TonB-dependent Receptor Plug Domain;<br>Uncharacterized protein YfbK, C-terminal;<br>von Willebrand factor;<br>CarboxypepD_reg-like domain | Inorganic ion transport and metabolism |
| Fjoh_4129 | A5FCD1 | SPI | not assigned | < 0.00010 | INF | | | | Function unknown |
| Fjoh_1847 | A5FIT9 | SPI | not assigned | 0.0081 | INF | | | Lipopolysaccharide-assembly | Function unknown |
| Fjoh_1351 | A5FK82 | SPII | periplasm-facing lipoprotein | 0.0076 | INF | | | GDSL-like Lipase/Acylhydrolase family | Amino acid transport and metabolism |
| Fjoh_1084 | A5FL08 | SPII | periplasm-facing lipoprotein | 0.00053 | INF | | | | Function unknown |
| Fjoh_3197 | A5FF10 | SPII | periplasm-facing lipoprotein | 0.00039 | INF | | | | Transcription |
| Fjoh_3307 | A5FEP9 | SPII | surface-exposed lipoprotein | 0.0081 | INF | | | Glucose / Sorbosone dehydrogenase | Carbohydrate transport and metabolism |
| Fjoh_4979 | A5F9Y7 | SPII | surface-exposed lipoprotein | < 0.00010 | INF | Pept_SE | | Beta-lactamase | Defense mechanisms |
| Fjoh_1783 | A5FJ07 | SPII | surface-exposed lipoprotein | < 0.00010 | INF | | | | Function unknown |
| Fjoh_2432 | A5FH57 | SPII | surface-exposed lipoprotein | 0.0050 | INF | SusD | | SusD family; Starch-binding associating with outer membrane | Function unknown |
| Fjoh_4673 | A5FAT9 | SPII | surface-exposed lipoprotein | 0.0021 | INF | | 31 | | Function unknown |
| Fjoh_3296 | A5FEQ3 | SPI | T9SS C-ter. sorting domain protein | 0.0031 | INF | | | Secretion system C-terminal sorting domain (type C) | Cell motility |
| Fjoh_4097 | A5FCF9 | SPI | periplasmic protein | 0.00051 | INF | GH127 | 22 or 25 | Beta-L-arabinofuranosidase, GH127 catalytic domain; Beta-L- | Function unknown |

| | | | | | | | | | |
|---|---|---|---|---|---|---|---|---|---|
| | | | | | | | | arabinofuranosidase, GH127 middle domain; Glycoside hydrolase family 127 C-terminal domain | |
| Fjoh_4098 | A5FCG0 | SPI | periplasmic protein | < 0.00010 | INF | PL1_2 | 22 or 25 | | Carbohydrate transport and metabolism |
| Fjoh_2471 | A5FH24 | SPI | periplasmic protein | < 0.00010 | INF | Pept_CA | | Transglutaminase-like superfamily | Cell cycle control, cell division, chromosome partitioning |
| Fjoh_0518 | A5FML7 | SPI | periplasmic protein | 0.00045 | INF | | | Protein of unknown function (DUF1501) | Function unknown |
| Fjoh_0954 | A5FLD8 | SPI | periplasmic protein | < 0.00010 | INF | | | | Function unknown |
| Fjoh_1633 | A5FJF0 | SPI | periplasmic protein | < 0.00010 | INF | | | SPOR domain | Function unknown |
| Fjoh_2122 | A5FI17 | SPI | periplasmic protein | < 0.00010 | INF | | | Glycosyl hydrolase-like 10 | Function unknown |
| Fjoh_4602 | A5FB22 | SPI | periplasmic protein | 0.00015 | INF | Pept_SC | | X-Pro dipeptidyl-peptidase (S15 family); X-Pro dipeptidyl-peptidase C-terminal non-catalytic domain | Function unknown |
| Fjoh_4603 | A5FB23 | SPI | periplasmic protein | < 0.00010 | INF | Pept_MH | | Peptidase family M20/M25/M40 | Function unknown |
| Fjoh_0976 | A5FLA5 | SPI | periplasmic protein | 0.00033 | INF | GH23; CBM50 | | LysM domain; Transglycosylase SLT domain | Cell wall/membrane/envelope biogenesis |
| Fjoh_2342 | A5FHF7 | SPII | surface-exposed lipoprotein | < 0.00010 | 67 | | | | Function unknown |
| Fjoh_1915 | A5FIM4 | SPI | periplasmic protein | < 0.00010 | 38 | | | | Cell motility; Inorganic ion transport and metabolism; Signal transduction mechanisms; Intracellular trafficking, secretion, and vesicular transport |
| Fjoh_0019 | A5FP10 | SPI | periplasmic protein | < 0.00010 | 28 | Pept_PA | | PDZ domain; Trypsin-like peptidase domain | Posttranslational modification, protein turnover, chaperones |
| Fjoh_0096 | A5FNT4 | SPI | periplasmic protein | 0.0011 | 27 | | | Domain of unknown function (DUF4252) | Function unknown |
| Fjoh_2959 | A5FFN3 | SPI | periplasmic protein | < 0.00010 | 26 | | | | Function unknown |
| Fjoh_0097 | A5FNT5 | SPII | periplasm-facing lipoprotein | < 0.00010 | 23 | | | Domain of unknown function (DUF4252) | Function unknown |
| Fjoh_0865 | A5FLM0 | SPI | periplasmic protein | < 0.00010 | 21 | | | LysM domain; Transglycosylase SLT domain | Function unknown |
| Fjoh_4779 | A5FAI4 | SPII | periplasm-facing lipoprotein | < 0.00010 | 19 | | | | RNA processing and modification |
| Fjoh_1698 | A5FJ83 | SPII | surface-exposed lipoprotein | 0.00050 | 18 | | | | Function unknown |
| Fjoh_2050 | A5FI90 | SPI | not assigned | 0.0022 | 15 | | | | Intracellular trafficking, secretion, and vesicular transport; Extracellular structures |
| sprE | A1E5T9 | SPII | periplasm-facing lipoprotein | 0.0053 | 14 | | | | Function unknown |
| Fjoh_4843 | A5FAC1 | SPII | surface-exposed lipoprotein | 0.0011 | 14 | | | Lipocalin-like domain | Function unknown |

| | | | | | | | | | |
|---|---|---|---|---|---|---|---|---|---|
| Fjoh_3189 | A5FF16 | SPI | periplasmic protein | 0.0029 | 14 | | | Bacterial virulence protein (VirJ) | Amino acid transport and metabolism; Intracellular trafficking, secretion, and vesicular transport |
| Fjoh_4130 | A5FCD2 | SPII | periplasm-facing lipoprotein | < 0.00010 | 13 | | | OEP family (Outer membrane efflux protein) | Cell wall/membrane/envelope biogenesis; Intracellular trafficking, secretion, and vesicular transport |
| Fjoh_0808 (RemA) | A5FLS4 | SPI | Integral/β-barrel protein | < 0.00010 | 10 | | | Galactose binding lectin domain | Defense mechanisms |
| Fjoh_1022 | A5FL64 | SPI | T9SS C-ter. sorting domain protein | 0.00040 | 9.7 | GH8 | | Secretion system C-terminal sorting domain (type A); Glycosyl hydrolases family 8 | Carbohydrate transport and metabolism |
| Fjoh_0546 | A5FMJ3 | SPII | surface-exposed lipoprotein | < 0.00010 | 9 | | | | Cell motility |
| Fjoh_2151 | A5FHZ5 | SPII | periplasm-facing lipoprotein | < 0.00010 | 7.2 | | | | Function unknown |
| Fjoh_0545 | A5FMJ2 | SPI | Integral/β-barrel protein | 0.00016 | 5.8 | SusC | | TonB dependent receptor-like, beta-barrel; TonB-dependent Receptor Plug Domain; CarboxypepD_reg-like domain | Inorganic ion transport and metabolism |
| Fjoh_1085 | A5FKZ5 | SPI | periplasmic protein | 0.00037 | 5.7 | | | Outer membrane lipoprotein carrier protein LolA | Cell wall/membrane/envelope biogenesis |
| Fjoh_4941 | A5FA29 | SPI | Integral/β-barrel protein | < 0.00010 | 5.6 | | | Outer membrane protein transport protein (OMPP1/FadL/TodX) | Lipid transport and metabolism |
| Fjoh_2150 | A5FHZ4 | SPI | T9SS C-ter. sorting domain protein | < 0.00010 | 5.5 | | | Secretion system C-terminal sorting domain | Energy production and conversion |
| Fjoh_3371 | A5FEI9 | SPII | surface-exposed lipoprotein | 0.0038 | 5.4 | | | Domain of unknown function (DUF4142) | Function unknown |
| Fjoh_0561 | A5FMH4 | SPII | surface-exposed lipoprotein | 0.0047 | 5.3 | | | | Function unknown |
| Fjoh_0415 | A5FMW7 | SPI | periplasmic protein | 0.00016 | 5 | | | Alpha-2-macroglobulin family; MG2 domain; Bacterial Alpha-2-macroglobulin MG10 domain | Function unknown |
| Fjoh_4293 | A5FBX4 | SPI | Integral/β-barrel protein | < 0.00010 | 4.8 | | | Outer membrane efflux protein | Cell wall/membrane/envelope biogenesis; Intracellular trafficking, secretion, and vesicular transport |
| Fjoh_0069 | A5FNV5 | SPII | surface-exposed lipoprotein | < 0.00010 | 4.5 | | | PrcB C-terminal | Function unknown |
| Fjoh_1393 | A5FK44 | SPII | surface-exposed lipoprotein | < 0.00010 | 4.3 | | | | Function unknown |
| Fjoh_1353 | A5FK84 | SPI | Integral/β-barrel protein | 0.0018 | 3.9 | | | Outer membrane efflux protein | Cell wall/membrane/envelope biogenesis; Intracellular trafficking, secretion, and vesicular transport |
| Fjoh_3415 | A5FEE1 | SPI | Integral/β-barrel protein | < 0.00010 | 3.8 | | | OmpA family | Cell wall/membrane/envelope biogenesis |
| Fjoh_0722 | A5FM14 | SPI | Integral/β-barrel protein | < 0.00010 | 3.6 | | | BatD DUF11 like domain | Function unknown |

| | | | | | | | | | |
|---|---|---|---|---|---|---|---|---|---|
| Fjoh_3240 | A5FEW1 | SPII | periplasm-facing lipoprotein | 0.0014 | 3.6 | | | OEP family (Outer membrane efflux protein) | Cell wall/membrane/envelope biogenesis; Intracellular trafficking, secretion, and vesicular transport |
| Fjoh_5000 | A5F9X2 | SPII | periplasm-facing lipoprotein | 0.0040 | 3.6 | | | | Function unknown |
| Fjoh_0270 | A5FNB1 | SPI | periplasmic protein | 0.00058 | 3.4 | | | SelR domain | Posttranslational modification, protein turnover, chaperones |
| Fjoh_0500 | A5FMN5 | SPI | periplasmic protein | < 0.00010 | 3.2 | Pept_SE | | Tetratricopeptide repeat; Beta-lactamase | Defense mechanisms |
| Fjoh_2111 | A5FI22 | SPI | periplasmic protein | < 0.00010 | 3 | | | Outer membrane lipoprotein carrier protein LolA | Cell wall/membrane/envelope biogenesis |
| Fjoh_0275 | A5FNB6 | SPII | periplasm-facing lipoprotein | < 0.00010 | 2.9 | | | META domain | Posttranslational modification, protein turnover, chaperones |
| Fjoh_2313 | A5FHH2 | SPI | periplasmic protein | < 0.00010 | 2.9 | | | YtxH-like protein | Function unknown |
| Fjoh_0831 | A5FLP9 | SPII | periplasm-facing lipoprotein | < 0.00010 | 2.8 | | | OmpA domain | Cell motility |
| Fjoh_4785 | A5FAJ0 | SPI | Integral/β-barrel protein | < 0.00010 | 2.7 | SusC | | TonB dependent receptor-like, beta-barrel;<br>TonB-dependent Receptor Plug Domain;<br>CarboxypepD_reg-like domain | Inorganic ion transport and metabolism |
| Fjoh_4940 | A5FA28 | SPII | periplasm-facing lipoprotein | < 0.00010 | 2.6 | | | | Function unknown |
| Fjoh_1777 | A5FJ14 | SPII | periplasm-facing lipoprotein | 0.00032 | 2.5 | | | | Function unknown |
| Fjoh_2314 | A5FHH3 | SPI | Integral/β-barrel protein | < 0.00010 | 2.4 | | | Outer membrane protein beta-barrel domain | Cell wall/membrane/envelope biogenesis |
| Fjoh_3759 | A5FDE8 | SPI | Integral/β-barrel protein | 0.0079 | 2.4 | | | | Lipid transport and metabolism |
| Fjoh_2057 | A5FI81 | SPII | periplasm-facing lipoprotein | < 0.00010 | 2.3 | | | | Function unknown |
| Fjoh_1067 | A5FL26 | SPI | periplasmic protein | < 0.00010 | 2.2 | Pept_MO | | Peptidase family M23 | Cell cycle control, cell division, chromosome partitioning |
| Fjoh_1907 | A5FIN2 | SPII | periplasm-facing lipoprotein | < 0.00010 | 2 | | | Domain of unknown function (DUF4136) | Function unknown |
| Fjoh_3469 | A5FE81 | SPII | periplasm-facing lipoprotein | 0.00081 | 2 | | | Outer membrane lipoprotein (BamD homologue) | Function unknown |
| Fjoh_0241 | A5FNE1 | SPI | Integral/β-barrel protein | 0.0020 | 1.9 | | | Outer membrane protein beta-barrel family | Inorganic ion transport and metabolism |
| Fjoh_2379 | A5FHB6 | SPI | periplasmic protein | 0.00025 | 1.9 | | | LysM domain | Amino acid transport and metabolism; Cell wall/membrane/envelope biogenesis |
| Fjoh_3108 | A5FF99 | SPI | T9SS C-ter. sorting domain protein | 0.0087 | 1.8 | | | Secretion system C-terminal sorting domain (type A) | Cell motility |
| Fjoh_0718 | A5FM10 | SPI | Integral/β-barrel protein | 0.0060 | 1.7 | | | | Posttranslational modification, protein turnover, chaperones |

| | | | | | | | | | |
|---|---|---|---|---|---|---|---|---|---|
| Fjoh_4343 | A5FBR6 | SPII | periplasm-facing lipoprotein | 0.0014 | 1.7 | | | Domain of unknown function (DUF4369) | Energy production and conversion; Posttranslational modification, protein turnover, chaperones |
| Fjoh_5007 | A5F9V9 | SPI | periplasmic protein | 0.0086 | 1.7 | | | Di-haem cytochrome c peroxidase | Energy production and conversion |
| Fjoh_0417 | A5FMW9 | SPI | Integral/β-barrel protein | 0.0022 | 1.6 | | | Outer membrane protein beta-barrel domain | Cell wall/membrane/envelope biogenesis |
| Fjoh_1522 | A5FJS1 | SPI | Integral/β-barrel protein | 0.0037 | 1.6 | | | Protein of unknown function (DUF3078) | Cell wall/membrane/envelope biogenesis |
| Fjoh_2750 | A5FGA4 | SPI | Integral/β-barrel protein | 0.0059 | 1.6 | | | LamB porin (for maltodextrin uptake) | Function unknown |
| Fjoh_5026 | A5F9U1 | SPI | Integral/β-barrel protein | 0.0057 | 1.6 | | | | Function unknown |
| Fjoh_1789 | A5FIZ8 | SPI | Integral/β-barrel protein | 0.00023 | 1.6 | | | | Transcription |
| Fjoh_1781 | A5FJ05 | SPI | not assigned | 0.0091 | 1.6 | Pept_NA | | YceI-like domain | Function unknown |
| Fjoh_2921 | A5FFS2 | SPII | periplasm-facing lipoprotein | 0.00012 | 1.6 | | | OmpA domain | Cell motility |
| Fjoh_1430 | A5FK03 | SPI | periplasmic protein | 0.0079 | 1.6 | Pept_SK | | C-terminal domain of tail specific protease (DUF3340); PDZ domain; Peptidase family S41; Tail specific protease N-terminal domain | Cell wall/membrane/envelope biogenesis |
| Fjoh_2737 | A5FGB5 | SPI | periplasmic protein | 0.00086 | 1.6 | | | Domain of unknown function (DUF4294) | Function unknown |
| Fjoh_4501 | A5FBC4 | SPI | periplasmic protein | 0.0044 | 1.6 | | | Protein of unknown function (DUF541) | Function unknown |
| Fjoh_0200 | A5FNI7 | SPII | periplasm-facing lipoprotein | 0.00060 | 0 | | | | Function unknown |
| Fjoh_3349 | A5FEL3 | SPI | integral/β-barrel protein | 0.0047 | 0 | | | | Cell wall/membrane/envelope biogenesis |
| Fjoh_1953 | A5FII9 | SPI | integral/β-barrel protein | 0.0050 | 0 | | | Kelch motif; CarboxypepD_reg-like domain | Cell wall/membrane/envelope biogenesis |
| Fjoh_2867 | A5FFZ0 | SPI | periplasmic protein | 0.0033 | 0 | Pept_MA | | Peptidase family M1 domain; Peptidase M1 N-terminal domain | Function unknown |
| Fjoh_3521 | A5FE32 | SPI | periplasmic protein | 0.00027 | 0 | GH3 | 16 or 20 | Glycosyl hydrolase family 3 N terminal domain; Glycosyl hydrolase family 3 C-terminal domain; Fibronectin type III-like domain | Function unknown |
| Fjoh_3861 | A5FD44 | SPI | periplasmic protein | 0.00013 | 0 | GH3 | 19 or 23 | Glycosyl hydrolase family 3 N terminal domain; Glycosyl hydrolase family 3 C-terminal domain; Fibronectin type III-like domain | Function unknown |

| | | | | | | | | | |
|---|---|---|---|---|---|---|---|---|---|
| Fjoh_1118 | A5FKX4 | SPI | periplasmic protein | 0.0020 | 0 | GH51_1 | | Alpha-L-arabinofuranosidase C-terminal domain | Function unknown |
| Fjoh_2042 | A5FI94 | SPI | periplasmic protein | 0.0053 | 0 | GH3 | 7 or 11 | Glycosyl hydrolase family 3 N terminal domain<br>Glycosyl hydrolase family 3 C-terminal domain<br>Fibronectin type III-like domain | Function unknown |
| Fjoh_4700 | A5FAR3 | SPI | periplasmic protein | 0.0034 | 0 | | | Lactonase, 7-bladed beta-propeller | Function unknown |
| Fjoh_4963 | A5FA08 | SPI | periplasmic protein | 0.0014 | 0 | GH3 | | Glycosyl hydrolase family 3 N terminal domain<br>Glycosyl hydrolase family 3 C-terminal domain<br>Fibronectin type III-like domain | Function unknown |
| Fjoh_0998 | A5FL81 | SPI | periplasmic protein | < 0.00010 | 0 | | | | Function unknown |
| Fjoh_0119 | A5FNQ4 | SPI | periplasmic protein | 0.00019 | 0 | Pept_MO | | Peptidase family M23 | Function unknown |
| Fjoh_2831 | A5FG24 | SPI | periplasmic protein | < 0.00010 | 0 | Pept_PB | | Linear amide C-N hydrolases, choloylglycine hydrolase family | Function unknown |
| Fjoh_3383 | A5FEG4 | SPI | periplasmic protein | 0.0010 | 0 | | | AhpC/TSA family | Function unknown |
| Fjoh_0827 | A5FLR2 | SPI | periplasmic protein | < 0.00010 | 0 | Est | | Putative esterase | Function unknown |
| Fjoh_1717 | A5FJ69 | SPI | periplasmic protein | 0.00010 | 0 | | | Serine aminopeptidase, S33 | Function unknown |
| Fjoh_1741 | A5FJ48 | SPI | periplasmic protein | < 0.00010 | 0 | | | | Function unknown |
| Fjoh_2853 | A5FFZ4 | SPI | periplasmic protein | 0.00032 | 0 | | | | Inorganic ion transport and metabolism |
| Fjoh_3206 | A5FEZ9 | SPI | periplasmic protein | 0.0066 | 0 | | | | Inorganic ion transport and metabolism |
| Fjoh_4606 | A5FB12 | SPI | periplasmic protein | 0.0063 | 0 | Pept_SC | | alpha/beta hydrolase fold | Inorganic ion transport and metabolism |
| Fjoh_4758 | A5FAK8 | SPI | periplasmic protein | 0.0077 | 0 | | | | Inorganic ion transport and metabolism |
| Fjoh_4273 | A5FBY8 | SPI | periplasmic protein | 0.00015 | 0 | | | Isochorismatase family | Secondary metabolites biosynthesis, transport and catabolism |
| Fjoh_4659 | A5FAV7 | SPI | periplasmic protein | < 0.00010 | 0 | | | Dienelactone hydrolase family | Secondary metabolites biosynthesis, transport and catabolism |
| Fjoh_2040 | A5FIA5 | SPI | periplasmic protein | 0.0010 | 0.04 | GH29 | 7 or 11 | Alpha-L-fucosidase; Alpha-L-fucosidase C-terminal domain | Function unknown |
| Fjoh_3392 | A5FEF5 | SPI | periplasmic protein | < 0.00010 | 0.06 | GH3 | | Fibronectin type III-like domain; Glycosyl hydrolase family 3 N terminal domain; Glycosyl hydrolase family 3 C-terminal domain | Function unknown |
| Fjoh_4757 | A5FAK7 | SPI | integral/β-barrel protein | < 0.00010 | 0.07 | | | Glycosyl hydrolases family 18 | Carbohydrate transport and metabolism |

| | | | | | | | | | |
|---|---|---|---|---|---|---|---|---|---|
| Fjoh_0276 | A5FN99 | SPI | periplasmic protein | 0.00019 | 0.07 | Pept_MG | | Metallopeptidase family M24; Aminopeptidase P, N-terminal domain | Energy production and conversion; Posttranslational modification, protein turnover, chaperones |
| Fjoh_4806 | A5FAG3 | SPI | periplasmic protein | 0.0016 | 0.08 | GH171 | | Exo-beta-N-acetylmuramidase NamZ, N-terminal | Inorganic ion transport and metabolism |
| Fjoh_3518 | A5FE42 | SPI | T9SS C-ter. sorting domain protein | < 0.00010 | 0.1 | Pept_MH | 16 | Peptidase family M28 | Amino acid transport and metabolism |
| Fjoh_2679 | A5FGG2 | SPII | periplasm-facing lipoprotein | 0.00048 | 0.1 | | | NTF2 fold immunity protein | Function unknown |
| Fjoh_1559 | A5FJN1 | SPI | integral/β-barrel protein | 0.00063 | 0.1 | | | | Energy production and conversion |
| Fjoh_1921 | A5FIL5 | SPII | periplasm-facing lipoprotein | 0.0012 | 0.1 | | | | Function unknown |
| Fjoh_1464 | A5FJW8 | SPII | integral/β-barrel protein | < 0.00010 | 0.1 | | | Omp85 superfamily domain | Carbohydrate transport and metabolism |
| Fjoh_1108 | A5FKY2 | SPI | not assigned | < 0.00010 | 0.1 | | | WG containing repeat | Transcription |
| Fjoh_1313 | A5FKD2 | SPI | periplasmic protein | 0.00055 | 0.1 | Pept_SC | | Prolyl oligopeptidase family; Prolyl oligopeptidase, N-terminal beta-propeller domain | Function unknown |
| Fjoh_4506 | A5FBB1 | SPI | periplasmic protein | < 0.00010 | 0.1 | Pept_PB | | Gamma-glutamyltranspeptidase | Function unknown |
| Fjoh_1556 | A5FJM8 | SPI | T9SS C-ter. sorting domain protein | 0.00068 | 0.2 | Pept_CD | | Peptidase family C25 | Amino acid transport and metabolism |
| Fjoh_4082 | A5FCI3 | SPII | periplasm-facing lipoprotein | 0.0077 | 0.2 | | | Glycosyl hydrolase family 65, C-terminal domain; Mannosylglycerate hydrolase MGH1-like glycoside hydrolase domain | Amino acid transport and metabolism |
| Fjoh_0974 | A5FLB6 | SPII | periplasm-facing lipoprotein | 0.0017 | 0.2 | | | | Function unknown |
| Fjoh_2451 | A5FH49 | SPI | periplasmic protein | < 0.00010 | 0.2 | | | Lactonase, 7-bladed beta-propeller | Function unknown |
| Fjoh_1722 | A5FJ62 | SPI | periplasmic protein | 0.0012 | 0.2 | | | Protein of unknown function (DUF1573) | Function unknown |
| Fjoh_2585 | A5FGQ1 | SPI | periplasmic protein | 0.00012 | 0.2 | | | Tetratricopeptide repeat; | Inorganic ion transport and metabolism |
| Fjoh_3486 | A5FE62 | SPI | periplasmic protein | 0.00055 | 0.2 | | | Uncharacterized protein conserved in bacteria (DUF2147) | Inorganic ion transport and metabolism |
| Fjoh_4816 | A5FAF7 | SPI | periplasmic protein | < 0.00010 | 0.2 | GH171 | 32 or 38 | Exo-beta-N-acetylmuramidase NamZ, N-terminal | Inorganic ion transport and metabolism |
| Fjoh_0959 | A5FLC8 | SPI | periplasmic protein | < 0.00010 | 0.2 | Pept_MA | | Peptidase family M13 | Posttranslational modification, protein turnover, chaperones |
| Fjoh_0983 | A5FL97 | SPI | T9SS C-ter. sorting domain protein | 0.0015 | 0.3 | | | T9SS C-terminal target domain-containing protein | Amino acid transport and metabolism |
| Fjoh_0422 | A5FMV9 | SPII | periplasm-facing lipoprotein | 0.0017 | 0.3 | | | Glucose / Sorbosone dehydrogenase | Amino acid transport and metabolism |
| Fjoh_2358 | A5FHD9 | SPII | periplasm-facing lipoprotein | 0.00041 | 0.3 | | | Glycosyl hydrolase family 92 catalytic domain; Glycosyl hydrolase family 92 N-terminal domain | Amino acid transport and metabolism |

| | | | | | | | | | |
|---|---|---|---|---|---|---|---|---|---|
| Fjoh_1913 | A5FIM2 | SPII | periplasm-facing lipoprotein | < 0.00010 | 0.3 | | | Mannosyl-glycoprotein endo-beta-N-acetylglucosaminidase; LysM domain | Cell wall/membrane/envelope biogenesis; Cell motility; Intracellular trafficking, secretion, and vesicular transport; |
| Fjoh_4478 | A5FBE8 | SPII | surface-exposed lipoprotein | 0.0014 | 0.3 | | | | Function unknown |
| Fjoh_1272 | A5FKG3 | SPI | integral/β-barrel protein | 0.00020 | 0.3 | | | PKD domain | Cell wall/membrane/envelope biogenesis |
| Fjoh_2008 | A5FIC5 | SPI | integral/β-barrel protein | 0.00017 | 0.3 | | | TonB dependent receptor-like, beta-barrel;<br>TonB-dependent Receptor Plug Domain;<br>CarboxypepD_reg-like domain | Cell wall/membrane/envelope biogenesis |
| Fjoh_4808 | A5FAG5 | SPI | periplasmic protein | < 0.00010 | 0.3 | GH20 | | Glycosyl hydrolase family 20, domain 2; Glycosyl hydrolase family 20, catalytic domain; Chitobiase/beta-hexosaminidase C-terminal domain | Function unknown |
| Fjoh_0225 | A5FNF7 | SPI | periplasmic protein | < 0.00010 | 0.3 | Pept_MA | | Peptidase family M48 | Function unknown |
| Fjoh_1318 | A5FKC2 | SPI | periplasmic protein | 0.0064 | 0.3 | | | Alanine racemase, C-terminal domain; Alanine racemase, N-terminal domain; Mur ligase middle domain | Function unknown |
| Fjoh_2626 | A5FGM2 | SPI | periplasmic protein | 0.00036 | 0.3 | Pept_SC | | alpha/beta hydrolase fold | Lipid transport and metabolism |
| Fjoh_3736 | A5FDH2 | SPI | periplasmic protein | < 0.00010 | 0.03 | Pept_SC | | alpha/beta hydrolase fold | Lipid transport and metabolism |
| Fjoh_2367 | A5FHB9 | SPI | periplasmic protein | < 0.00010 | 0.3 | | | Cyclophilin type peptidyl-prolyl cis-trans isomerase/CLD; FKBP-type peptidyl-prolyl cis-trans isomerase | Posttranslational modification, protein turnover, chaperones |
| Fjoh_0820 | A5FLS2 | SPII | periplasm-facing lipoprotein | 0.00024 | 0.4 | | | | Amino acid transport and metabolism |
| Fjoh_0454 | A5FMS1 | SPII | periplasm-facing lipoprotein | < 0.00010 | 0.4 | | | Peptidase family M28 | Function unknown |
| Fjoh_0980 (SprD) | A1E5U4 | SPI | integral/β-barrel protein | 0.00015 | 0.4 | | | Type IX secretion system membrane protein PorP/SprF | Carbohydrate transport and metabolism |
| Fjoh_1476 | A5FJW3 | SPI | integral/β-barrel protein | < 0.00010 | 0.4 | | | CarboxypepD_reg-like domain; Family of unknown function (DUF5686) | Carbohydrate transport and metabolism |
| Fjoh_1517 | A5FJR6 | SPI | integral/β-barrel protein | 0.0076 | 0.4 | | | Outer membrane protein beta-barrel domain | Cell cycle control, cell division, chromosome partitioning; Cell wall/membrane/envelope biogenesis |
| Fjoh_3179 | A5FF33 | SPI | integral/β-barrel protein | 0.0054 | 0.4 | | | TonB dependent receptor-like, beta-barrel;<br>TonB-dependent Receptor Plug Domain;<br>CarboxypepD_reg-like domain | Cell motility |

| | | | | | | | | | |
|---|---|---|---|---|---|---|---|---|---|
| Fjoh_0105 | A5FNS6 | SPI | integral/β-barrel protein | 0.00017 | 0.4 | | | Phosphate-selective porin O and P | Cell wall/membrane/envelope biogenesis |
| Fjoh_0665 | A5FM73 | SPI | integral/β-barrel protein | 0.0061 | 0.4 | | | TonB dependent receptor-like, beta-barrel; TonB-dependent Receptor Plug Domain; CarboxypepD_reg-like domain | Cell wall/membrane/envelope biogenesis |
| Fjoh_4194 | A5FC74 | SPI | integral/β-barrel protein | 0.0068 | 0.4 | | | TonB dependent receptor-like, beta-barrel; TonB-dependent Receptor Plug Domain; CarboxypepD_reg-like domain | Coenzyme transport and metabolism |
| Fjoh_4255 | A5FC08 | SPI | integral/β-barrel protein | 0.0013 | 0.4 | | | TonB dependent receptor-like, beta-barrel; TonB-dependent Receptor Plug Domain; CarboxypepD_reg-like domain | Energy production and conversion |
| Fjoh_1562 | A5FJM0 | SPI | periplasmic protein | 0.00068 | 0.4 | GH51_1 | | Glycosyl hydrolase family 30 TIM-barrel domain | Function unknown |
| Fjoh_1564 | A5FJM2 | SPI | periplasmic protein | 0.0012 | 0.4 | GH30_1 | 4 or 7 | Fibronectin type III-like domain; Glycosyl hydrolase family 3 N terminal domain; Glycosyl hydrolase family 3 C-terminal domain | Function unknown |
| Fjoh_1688 | A5FJA1 | SPI | periplasmic protein | 0.00025 | 0.4 | | | Outer membrane protein (OmpH-like) | Function unknown |
| Fjoh_3906 | A5FD04 | SPI | periplasmic protein | 0.0049 | 0.4 | | | NADH ubiquinone oxidoreductase, 20 Kd subunit; NiFe/NiFeSe hydrogenase small subunit C-terminal | Function unknown |
| Fjoh_1780 | A5FJ04 | SPI | periplasmic protein | < 0.00010 | 0.4 | Pept_na | | YceI-like domain | Function unknown |
| Fjoh_2036 | A5FIA1 | SPII | periplasm-facing lipoprotein | 0.0061 | 0.5 | | | Putative glycosyl hydrolase domain | Function unknown |
| Fjoh_2832 | A5FG07 | SPII | periplasm-facing lipoprotein | 0.0017 | 0.5 | | | | Function unknown |
| Fjoh_0576 | A5FMF7 | SPI | integral/β-barrel protein | < 0.00010 | 0.5 | | | CarboxypepD_reg-like domain; Family of unknown function (DUF5686) | Carbohydrate transport and metabolism |
| Fjoh_1173 | A5FKR4 | SPI | integral/β-barrel protein | 0.0048 | 0.5 | | | Protein of unknown function (DUF3078) | Carbohydrate transport and metabolism |
| Fjoh_4485 | A5FBD6 | SPI | integral/β-barrel protein | < 0.00010 | 0.5 | | | Outer membrane efflux protein | Cell motility |
| Fjoh_1779 | A5FJ03 | SPI | integral/β-barrel protein | 0.0086 | 0.5 | | | | Cell wall/membrane/envelope biogenesis |

| | | | | | | | | | |
|---|---|---|---|---|---|---|---|---|---|
| Fjoh_2466 | A5FH32 | SPI | integral/β-barrel protein | < 0.00010 | 0.5 | | | TonB dependent receptor-like, beta-barrel; TonB-dependent Receptor Plug Domain; CarboxypepD_reg-like domain | Cell wall/membrane/envelope biogenesis |
| Fjoh_3882 | A5FD25 | SPI | integral/β-barrel protein | 0.00034 | 0.5 | | | TonB dependent receptor-like, beta-barrel; TonB-dependent Receptor Plug Domain; CarboxypepD_reg-like domain | Cell wall/membrane/envelope biogenesis; Intracellular trafficking, secretion, and vesicular transport |
| Fjoh_4221 | A5FC34 | SPI | integral/β-barrel protein | 0.00039 | 0.5 | | | TonB dependent receptor-like, beta-barrel; TonB-dependent Receptor Plug Domain; CarboxypepD_reg-like domain | Energy production and conversion |
| Fjoh_0416 | A5FMW8 | SPI | periplasmic protein | 0.0022 | 0.5 | Pept_PA | | Peptidase S46 | Function unknown |
| Fjoh_1419 | A5FK22 | SPI | periplasmic protein | < 0.00010 | 0.5 | Pept_SC | | Dipeptidyl peptidase IV (DPP IV) N-terminal region; Prolyl oligopeptidase family | Function unknown |
| Fjoh_1567 | A5FJM5 | SPI | periplasmic protein | < 0.00010 | 0.5 | GH3 | 4 or 7 | Fibronectin type III-like domain; Glycosyl hydrolase family 3 N terminal domain; Glycosyl hydrolase family 3 C-terminal domain | Function unknown |
| Fjoh_1191 | A5FKP8 | SPI | periplasmic protein | 0.0011 | 0.5 | Pept_MA | | Peptidase family M1 domain | Function unknown |
| Fjoh_4786 (FumC) | A5FAJ1 | SPI | periplasmic protein | 0.0056 | 0.5 | | | Lyase; Fumarase C C-terminus | Function unknown |
| Fjoh_1778 | A5FJ15 | SPI | periplasmic protein | 0.0031 | 0.5 | | | YceI-like domain | Function unknown |
| Fjoh_3122 | A5FF84 | SPI | periplasmic protein | 0.0023 | 0.5 | | 13 | Domain of unknown function (DUF4861) | Inorganic ion transport and metabolism |
| Fjoh_4809 | A5FAG6 | SPI | periplasmic protein | < 0.00010 | 0.5 | | | Glycosyl hydrolase-like 10 | Inorganic ion transport and metabolism |
| Fjoh_3473 | A5FE85 | SPI | periplasmic protein | 0.0016 | 0.5 | | | Calcineurin-like phosphoesterase | Nucleotide transport and metabolism |
| Fjoh_2944 | A5FFP8 | SPI | periplasmic protein | 0.00094 | 0.5 | | | Glutathione peroxidase | Posttranslational modification, protein turnover, chaperones |
| Fjoh_0117 | A5FNS0 | SPII | surface-exposed lipoprotein | 0.0065 | 0.6 | | | | Function unknown |
| Fjoh_3524 | A5FE35 | SPII | surface-exposed lipoprotein | 0.0043 | 0.6 | | | SusD family; Starch-binding associating with outer membrane | Function unknown |
| Fjoh_5008 | A5F9W0 | SPI | integral/β-barrel protein | < 0.00010 | 0.6 | | | | Cell wall/membrane/envelope biogenesis |
| Fjoh_0185 | A5FNK1 | SPI | integral/β-barrel protein | < 0.00010 | 0.6 | | | TonB dependent receptor-like, beta-barrel; TonB-dependent Receptor Plug | Cell wall/membrane/envelope biogenesis |

| | | | | | | | | | |
|---|---|---|---|---|---|---|---|---|---|
| | | | | | | | | Domain;<br>CarboxypepD_reg-like domain | |
| Fjoh_0782 | A5FLW0 | SPI | integral/β-barrel protein | 0.00037 | 0.6 | | | TonB dependent receptor-like, beta-barrel;<br>TonB-dependent Receptor Plug Domain;<br>CarboxypepD_reg-like domain | Cell wall/membrane/envelope biogenesis |
| Fjoh_4039 | A5FCM0 | SPI | integral/β-barrel protein | 0.0060 | 0.6 | | | TonB dependent receptor-like, beta-barrel;<br>TonB-dependent Receptor Plug Domain;<br>CarboxypepD_reg-like domain | Cell wall/membrane/envelope biogenesis; Intracellular trafficking, secretion, and vesicular transport; Cell motility |
| Fjoh_5040 | A5F9T6 | SPI | periplasmic protein | 0.0018 | 0.6 | | | | Inorganic ion transport and metabolism |
| Fjoh_1007 | A5FL74 | SPI | periplasmic protein | 0.0041 | 0.6 | | | SPFH domain / Band 7 family | Posttranslational modification, protein turnover, chaperones |

**Supplementary Table S2. Proteins of outer membrane vesicles (OMVs), isolated from cells grown in permissive (+ IPTG) and non-permissive (- IPTG) conditions and sorted in descending order, identified by label-free mass spectrometry whose spectra count is significantly different (FC ≥ |1.5|) between the two conditions**. Listed are: Protein name; Accession number (as in Uniprot); Signal Peptide (SP) prediction based on SignalP 6.0 [4]; *p*-value (as given by a two-tailed *t*-test); FC: fold change (- IPTG/+ IPTG); Modularity and PULs (as in CAZy database [5]); Protein family (PFAM domain annotation); Gene Ontology (as annotated in MaGe [6]). Empty cells indicate annotation not available. "INF" and "0" were assigned when no peptide was detected in permissive (+ IPTG) or non-permissive (- IPTG) conditions of growth, respectively.

| Protein name | Accession number | Signal peptide | Type | *p*-value | FC | Modularity (CAZy) | PUL (CAZy) | Protein family | Gene Ontology |
|---|---|---|---|---|---|---|---|---|---|
| Fjoh_0519 | A5FML8 | SPI | periplasmic protein | 0.0001 | INF | | | Protein of unknown function (DUF1800) | Function unknown |
| Fjoh_1118 | A5FKX4 | SPI | periplasmic protein | 0.00081 | INF | GH51_1 | | Alpha-L-arabinofuranosidase C-terminal domain | Function unknown |
| Fjoh_1224 | A5FKL1 | SPI | periplasmic protein | 0.0074 | INF | | | cAMP phosphodiesterases class-II | Signal transduction mechanisms |
| Fjoh_1360 | A5FK78 | SPI | periplasmic protein | 0.00024 | INF | Pept_MH | | Peptidase family M20/M25/M40 | Amino acid transport and metabolism |
| Fjoh_1494 | A5FJU6 | SPI | periplasmic protein | 0.0031 | INF | | | Amino acid kinase family; Homoserine dehydrogenase, NAD binding domain; Homoserine dehydrogenase | Amino acid transport and metabolism |
| Fjoh_2023 | A5FIB4 | SPI | periplasmic protein | 0.0022 | INF | GH43_18 | 6 or 10 | | Carbohydrate transport and metabolism |
| Fjoh_2082 | A5FI57 | SPI | periplasmic protein | 0.00044 | INF | GH2 | 8 or 12 | Glycosyl hydrolases family 2, sugar binding domain; Glycosyl hydrolases family 2; Glycosyl hydrolases family 2, TIM barrel domain | Carbohydrate transport and metabolism |
| Fjoh_2338 | A5FHF3 | SPI | periplasmic protein | 0.0071 | INF | | | Secretion system C-terminal sorting domain | Function unknown |
| Fjoh_2400 | A5FH86 | SPI | periplasmic protein | 0.0017 | INF | Pept_MH | | Peptidase family M20/M25/M40 | Amino acid transport and metabolism |
| Fjoh_2751 | A5FGA5 | SPI | periplasmic protein | < 0.00010 | INF | | | | Function unknown |
| Fjoh_2809 | A5FG34 | SPI | periplasmic protein | 0.0012 | INF | | | | |
| Fjoh_4074 | A5FCI9 | SPI | periplasmic protein | < 0.00010 | INF | GH2 | 25 | Glycosyl hydrolases family 2, sugar binding domain; Glycosyl hydrolases family 2; Glycosyl hydrolases family 2, TIM barrel domain; Domain of unknown function (DUF4982); Glycoside hydrolase family 2 C-terminal domain 5 | Carbohydrate transport and metabolism |
| Fjoh_4087 | A5FCH5 | SPI | periplasmic protein | 0.007 | INF | GH43_18 | 22 or 25 | Glycosyl hydrolases family 43 | Carbohydrate transport and metabolism |
| Fjoh_4249 | A5FC13 | SPI | periplasmic protein | 0.00036 | INF | GH43_10/CBM91 | 26 or 29 | Glycosyl hydrolases family 43; Beta xylosidase C-terminal Concanavalin A-like domain | Carbohydrate transport and metabolism |

| | | | | | | | | | |
|---|---|---|---|---|---|---|---|---|---|
| Fjoh_4602 | A5FB22 | SPI | periplasmic protein | 0.00035 | INF | Pept_SC | | X-Pro dipeptidyl-peptidase (S15 family); X-Pro dipeptidyl-peptidase C-terminal non-catalytic domain | Function unknown |
| Fjoh_4724(PurL) | A5FAQ1 | SPI | periplasmic protein | 0.00048 | INF | | | Formylglycinamide ribonucleotide amidotransferase N-terminal; Formylglycinamide ribonucleotide amidotransferase linker domain; AIR synthase related protein, C-terminal domain; AIR synthase related protein, C-terminal domain; CobB/CobQ-like glutamine amidotransferase domain | Nucleotide transport and metabolism |
| Fjoh_4959 | A5FA11 | SPI | periplasmic protein | 0.0001 | INF | GH97 | 33 or 39 | Glycosyl-hydrolase 97 N-terminal; Glycoside hydrolase 97; Glycosyl-hydrolase 97 C-terminal, oligomerisation | Cell wall/membrane/envelope biogenesis |
| Fjoh_1915 | A5FIM4 | SPI | periplasmic protein | < 0,00010 | 38 | | | | Cell motility; Inorganic ion transport and metabolism; Signal transduction mechanisms; Intracellular trafficking, secretion, and vesicular transport |
| Fjoh_3942 | A5FCW1 | SPI | periplasmic protein | 0.0064 | 38 | Pept_SC | 21 or 24 | Prolyl oligopeptidase family | Amino acid transport and metabolism |
| Fjoh_5035 | A5F9T1 | SPI | Integral/β-barrel protein | 0.00026 | 33 | SusC | | von Willebrand factor type A domain; TonB-dependent Receptor Plug Domain; Uncharacterized protein YfbK, C-terminal;von Willebrand factor; CarboxypepD_reg-like domain | Inorganic ion transport and metabolism |
| Fjoh_4083 | A5FCI4 | SPI | periplasmic protein | 0.00015 | 31 | GH95 | 22 or 25 | Glycosyl hydrolase family 65, N-terminal domain | Carbohydrate transport and metabolism |
| Fjoh_4102 | A5FCG4 | SPI | periplasmic protein | 0.00052 | 31 | GH2 | 23 or 25 | Glycosyl hydrolases family 2, sugar binding domain; Glycosyl hydrolases family 2; Glycosyl hydrolases family 2, TIM barrel domain; Beta-galactosidase, domain 4; Beta galactosidase small chain | Carbohydrate transport and metabolism |
| Fjoh_3111 | A5FF86 | SPI | periplasmic protein | < 0.00010 | 29 | GH2 | 13 | Glycosyl hydrolases family 2, sugar binding domain; Glycosyl hydrolases family 2; Glycosyl hydrolases family 2, TIM barrel domain; Domain of unknown function (DUF4982); Glycoside | Carbohydrate transport and metabolism |

| | | | | | | | | | |
|---|---|---|---|---|---|---|---|---|---|
| | | | | | | | | hydrolase family 2 C-terminal domain 5 | |
| Fjoh_4090 | A5FCG5 | SPI | periplasmic protein | < 0.00010 | 22 | | 22 or 25 | Family of unknown function (DUF6298) | Cell wall/membrane/envelope biogenesis |
| Fjoh_1916 | A5FIM5 | SPI | periplasmic protein | < 0.00010 | 19 | | | | Function unknown |
| Fjoh_2042 | A5FI94 | SPI | periplasmic protein | 0.00014 | 19 | GH3 | 7 or 11 | Glycosyl hydrolase family 3 N terminal domain; Glycosyl hydrolase family 3 C-terminal domain; Fibronectin type III-like domain | Function unknown |
| Fjoh_4777 | A5FAJ7 | SPI | periplasmic protein | 0.0085 | 19 | | | | Function unknown |
| Fjoh_1789 | A5FIZ8 | SPI | Integral/β-barrel protein | 0.00036 | 18 | | | | Transcription |
| Fjoh_2959 | A5FFN3 | SPI | periplasmic protein | 0.0032 | 18 | | | | Function unknown |
| Fjoh_0500 | A5FMN5 | SPI | periplasmic protein | 0.00011 | 17 | Pept_SE | | Tetratricopeptide repeat; Beta-lactamase | Defense mechanisms |
| Fjoh_2781 (MenD) | A5FG67 | SPI | periplasmic protein | < 0.00010 | 17 | | | Thiamine pyrophosphate enzyme, N-terminal TPP binding domain; Middle domain of thiamine pyrophosphate; Thiamine pyrophosphate enzyme, C-terminal TPP binding domain | Function unknown |
| Fjoh_1209 | A5FKN5 | SPI | periplasmic protein | 0.00046 | 16 | GH97 | 2 or 5 | Glycosyl-hydrolase 97 C-terminal, oligomerisation; Glycosyl-hydrolase 97 N-terminal; Glycoside hydrolase 97 | Defense mechanisms |
| Fjoh_4501 | A5FBC4 | SPI | periplasmic protein | < 0.00010 | 16 | | | Protein of unknown function (DUF541) | Function unknown |
| Fjoh_0468 | A5FMR6 | SPI | periplasmic protein | 0.0032 | 15 | | | | Function unknown |
| Fjoh_3880 | A5FD23 | SPI | periplasmic protein | < 0.00010 | 15 | GH43_12/CBM91 | 20 or 23 | Glycosyl hydrolases family 43; Beta xylosidase C-terminal Concanavalin A-like domain | Carbohydrate transport and metabolism |
| Fjoh_4603 | A5FB23 | SPI | periplasmic protein | 0.0016 | 15 | Pept_MH | | Peptidase family M20/M25/M40 | Function unknown |
| Fjoh_0271 (MsrA) | A5FNB2 | SPI | periplasmic protein | 0.0014 | 14 | | | Peptide methionine sulfoxide reductase | Posttranslational modification, protein turnover, chaperones |
| Fjoh_3861 | A5FD44 | SPI | periplasmic protein | 0.00075 | 14 | GH3 | 19 or 23 | Glycosyl hydrolase family 3 N terminal domain; Glycosyl hydrolase family 3 C-terminal domain; Fibronectin type III-like domain | Function unknown |
| Fjoh_4086 | A5FCH4 | SPI | periplasmic protein | < 0.00010 | 14 | GH143 | 22 or 25 | | Carbohydrate transport and metabolism |
| Fjoh_4481 | A5FBD2 | SPI | periplasmic protein | 0.00056 | 14 | | | Protein of unknown function (DUF1349) | Function unknown |

| | | | | | | | | | |
|---|---|---|---|---|---|---|---|---|---|
| Fjoh_0096 | A5FNT4 | SPI | periplasmic protein | < 0.00010 | 13 | | | Domain of unknown function (DUF4252) | Function unknown |
| Fjoh_0097 | A5FNT5 | SPII | periplasm-facing lipoprotein | 0.0039 | 13 | | | Domain of unknown function (DUF4252) | Function unknown |
| Fjoh_3877 | A5FD33 | SPI | periplasmic protein | 0.0055 | 13 | GH115 | 20 or 23 | Glycosyl hydrolase family 115; Gylcosyl hydrolase family 115 C-terminal domain | Function unknown |
| Fjoh_2714 | A5FGD3 | SPI | periplasmic protein | 0.0021 | 12 | GH92 | 10 or 14 | Glycosyl hydrolase family 92 catalytic domain; Glycosyl hydrolase family 92 N-terminal domain | Carbohydrate transport and metabolism |
| Fjoh_3473 | A5FE85 | SPI | periplasmic protein | 0.0081 | 12 | | | Calcineurin-like phosphoesterase | Nucleotide transport and metabolism |
| Fjoh_4080 | A5FCI1 | SPI | periplasmic protein | 0.0017 | 12 | GH78 | 22 or 25 | Alpha-L-rhamnosidase N-terminal domain; Bacterial alpha-L-rhamnosidase concanavalin-like domain; Bacterial alpha-L-rhamnosidase 6 hairpin glycosidase domain; Bacterial alpha-L-rhamnosidase C-terminal domain | Carbohydrate transport and metabolism |
| Fjoh_2253 | A5FHP1 | SPI | periplasmic protein | < 0.00010 | 11 | Pept_ME | | Insulinase (Peptidase family M16); Peptidase M16 inactive domain | Function unknown |
| Fjoh_2626 | A5FGM2 | SPI | periplasmic protein | 0.0068 | 11 | Pept_SC | | alpha/beta hydrolase fold | Lipid transport and metabolism |
| Fjoh_2716 | A5FGD5 | SPI | periplasmic protein | 0.0069 | 11 | GH125 | 10 or 14 | Metal-independent alpha-mannosidase (GH125) | Function unknown |
| Fjoh_4915 | A5FA53 | SPI | periplasmic protein | 0.0012 | 11 | | | Uncharacterized protein conserved in bacteria (DUF2141) | Function unknown |
| Fjoh_2038 | A5FIA3 | SPI | periplasmic protein | 0.00049 | 10 | GH2 | 7 or 11 | Beta galactosidase small chain; Glycosyl hydrolases family 2; Glycosyl hydrolases family 2, TIM barrel domain; Glycosyl hydrolases family 2, sugar binding domain; Beta-galactosidase, domain 4 | Carbohydrate transport and metabolism |
| Fjoh_4198 | A5FC62 | SPI | periplasmic protein | 0.0002 | 9.9 | GH2 | 24 or 27 | Glycosyl hydrolases family 2, sugar binding domain; Glycosyl hydrolases family 2; Glycosyl hydrolases family 2, TIM barrel domain | Carbohydrate transport and metabolism |
| Fjoh_3521 | A5FE32 | SPI | periplasmic protein | 0.0013 | 9.8 | GH3 | 16 or 20 | Glycosyl hydrolase family 3 N terminal domain; Glycosyl hydrolase family 3 C-terminal domain; Fibronectin type III-like domain | Function unknown |

| | | | | | | | | | |
|---|---|---|---|---|---|---|---|---|---|
| Fjoh_2750 | A5FGA4 | SPI | Integral/β-barrel protein | 0.0002 | 9.5 | | | LamB porin (for maltodextrin uptake) | Function unknown |
| Fjoh_3113 | A5FF88 | SPI | periplasmic protein | 0.00023 | 9.2 | GH51_2 | 13 | Alpha-L-arabinofuranosidase C-terminal domain | Carbohydrate transport and metabolism |
| Fjoh_4097 | A5FCF9 | SPI | periplasmic protein | 0.00015 | 8.8 | GH127 | 22 or 25 | Beta-L-arabinofuranosidase, GH127 catalytic domain; Beta-L-arabinofuranosidase, GH127 middle domain; Glycoside hydrolase family 127 C-terminal domain | Function unknown |
| Fjoh_0846 | A5FLP6 | SPI | periplasmic protein | 0.0038 | 8.4 | | | Tetratricopeptide repeat | Function unknown |
| Fjoh_2342 | A5FHF7 | SPII | surface-exposed lipoprotein | 0.00011 | 8.4 | | | | Function unknown |
| Fjoh_3873 | A5FD29 | SPI | periplasmic protein | 0.0005 | 8 | GH97 | 20 or 23 | Glycosyl-hydrolase 97 N-terminal; Glycoside hydrolase 97; Glycosyl-hydrolase 97 C-terminal, oligomerisation | Function unknown |
| Fjoh_3817 | A5FD95 | SPI | periplasmic protein | 0.0034 | 7.7 | Pept_SE | | Beta-lactamase | Defense mechanisms |
| Fjoh_4429 | A5FBI0 | SPI | periplasmic protein | 0.00044 | 7.7 | GH97 | 29 or 32 | Glycosyl-hydrolase 97 C-terminal, oligomerisation; Glycosyl-hydrolase 97 N-terminal; Glycoside hydrolase 97 | Defense mechanisms |
| Fjoh_2961 | A5FFN5 | SPI | periplasmic protein | < 0.00010 | 7.6 | Est | | GlcNAc-PI de-N-acetylase | Function unknown |
| Fjoh_1562 | A5FJM0 | SPI | periplasmic protein | 0.0023 | 7.4 | GH51_1 | | Glycosyl hydrolase family 30 TIM-barrel domain | Function unknown |
| Fjoh_3389 | A5FEH0 | SPI | periplasmic protein | < 0.00010 | 7.2 | GH3 | | Glycosyl hydrolase family 3 N terminal domain; Glycosyl hydrolase family 3 C-terminal domain; Fibronectin type III-like domain | Carbohydrate transport and metabolism |
| Fjoh_1541 | A5FJP1 | SPII | periplasm-facing lipoprotein | 0.0098 | 7.1 | | | Heavy-metal-associated domain | Inorganic ion transport and metabolism |
| Fjoh_4995 | A5F9X6 | SPI | periplasmic protein | 0.0064 | 6.7 | | | | Carbohydrate transport and metabolism |
| Fjoh_3194 | A5FF07 | SPI | T9SS C-ter. -sorting domain protein | < 0.00010 | 6.6 | | | Type IX secretion system membrane protein PorP/SprF | Cell motility |
| Fjoh_4808 | A5FAG5 | SPI | periplasmic protein | < 0.00010 | 6 | GH20 | | Glycosyl hydrolase family 20, domain 2; Glycosyl hydrolase family 20, catalytic domain; Chitobiase/beta-hexosaminidase C-terminal domain | Function unknown |
| Fjoh_0778 | A5FLV6 | SPI | periplasmic protein | 0.0072 | 5.8 | GH31_3 | 1 or 4 | Glycosyl hydrolase 31 N-terminal galactose mutarotase-like domain; Glycosyl hydrolases family 31 TIM-barrel domain; Glycosyl | Carbohydrate transport and metabolism |

| | | | | | | | | | |
|---|---|---|---|---|---|---|---|---|---|
| | | | | | | | | hydrolase family 31 C-terminal domain | |
| Fjoh_4088 | A5FCH6 | SPI | periplasmic protein | 0.0015 | 5.7 | GH28 | 22 or 25 | Pectate lyase superfamily protein; Glycosyl hydrolases family 28 | Carbohydrate transport and metabolism |
| Fjoh_1400 | A5FK37 | SPI | periplasmic protein | < 0.00010 | 5.6 | GH97 | 3 | Glycosyl-hydrolase 97 C-terminal, oligomerisation; Glycosyl-hydrolase 97 N-terminal; Glycoside hydrolase 97 | Carbohydrate transport and metabolism |
| Fjoh_0119 | A5FNQ4 | SPI | periplasmic protein | 0.00026 | 5.3 | Pept_MO | | Peptidase family M23 | Cell wall/membrane/envelope biogenesis |
| Fjoh_4779 | A5FAI4 | SPII | periplasm-facing lipoprotein | 0.00054 | 5.3 | | | | RNA processing and modification |
| Fjoh_1419 | A5FK22 | SPI | periplasmic protein | < 0.00010 | 5.2 | Pept_SC | | Dipeptidyl peptidase IV (DPP IV) N-terminal region; Prolyl oligopeptidase family | Function unknown |
| Fjoh_2341 | A5FHF6 | SPI | periplasmic protein | 0.0017 | 5.2 | Est | | Putative esterase | Function unknown |
| Fjoh_2041 | A5FI93 | SPI | periplasmic protein | 0.0055 | 5.1 | GH92 | 7 or 11 | Glycosyl hydrolase family 92 N-terminal domain; Glycosyl hydrolase family 92 catalytic domain | Carbohydrate transport and metabolism |
| Fjoh_3800 | A5FDB0 | SPI | periplasmic protein | 0.0036 | 5.1 | | 18 or 22 | Domain of unknown function (DUF5118); Domain of unknown function (DUF5117); Met-zincin | Posttranslational modification, protein turnover, chaperones |
| Fjoh_1430 | A5FK03 | SPI | periplasmic protein | 0.00028 | 5 | Pept_SK | | C-terminal domain of tail specific protease (DUF3340); PDZ domain; Peptidase family S41; Tail specific protease N-terminal domain | Cell wall/membrane/envelope biogenesis |
| Fjoh_3874 | A5FD30 | SPI | periplasmic protein | 0.0017 | 4.9 | GH3 | 20 or 23 | Glycosyl hydrolase family 3 N terminal domain; Glycosyl hydrolase family 3 C-terminal domain; Fibronectin type III-like domain | Carbohydrate transport and metabolism |
| Fjoh_2358 | A5FHD9 | SPII | periplasm-facing lipoprotein | < 0.00010 | 4.8 | | | Glycosyl hydrolase family 92 catalytic domain; Glycosyl hydrolase family 92 N-terminal domain | Amino acid transport and metabolism |
| Fjoh_1191 | A5FKP8 | SPI | periplasmic protein | < 0.00010 | 4.7 | Pept_MA | | Peptidase family M1 domain | Function unknown |
| Fjoh_1564 | A5FJM2 | SPI | periplasmic protein | 0.0043 | 4.6 | GH30_1 | 4 or 7 | Fibronectin type III-like domain; Glycosyl hydrolase family 3 N terminal domain; Glycosyl hydrolase family 3 C-terminal domain | Function unknown |
| Fjoh_3437 | A5FEB2 | SPI | periplasmic protein | < 0.00010 | 4.4 | Pept_MA | | Peptidase family M1 domain | Amino acid transport and metabolism |
| Fjoh_0200 | A5FNI7 | SPII | periplasm-facing lipoprotein | 0.00035 | 4.2 | | | | Function unknown |

| | | | | | | | | | |
|---|---|---|---|---|---|---|---|---|---|
| Fjoh_0415 | A5FMW7 | SPI | periplasmic protein | 0.0002 | 4.2 | | | Alpha-2-macroglobulin family; MG2 domain; Bacterial Alpha-2-macroglobulin MG10 domain | Function unknown |
| Fjoh_3392 | A5FEF5 | SPI | periplasmic protein | 0.0069 | 4.1 | GH3 | | Fibronectin type III-like domain; Glycosyl hydrolase family 3 N terminal domain; Glycosyl hydrolase family 3 C-terminal domain | Carbohydrate transport and metabolism |
| Fjoh_0023 | A5FP14 | SPI | periplasmic protein | < 0.00010 | 4 | | | Type I phosphodiesterase / nucleotide pyrophosphatase | Function unknown |
| Fjoh_0246 | A5FND2 | SPI | periplasmic protein | 0.0026 | 4 | CBM50/Est | | GDSL-like Lipase/Acylhydrolase family; LysM domain | Amino acid transport and metabolism; Cell wall/membrane/envelope biogenesis |
| Fjoh_2181 | A5FHV4 | SPI | periplasmic protein | 0.0003 | 4 | | | Monomeric isocitrate dehydrogenase | Energy production and conversion |
| Fjoh_2715 | A5FGD4 | SPI | periplasmic protein | 0.00087 | 4 | GH92 | 10 or 14 | Glycosyl hydrolase family 92 N-terminal domain; Glycosyl hydrolase family 92 catalytic domain | Carbohydrate transport and metabolism |
| Fjoh_3114 | A5FF89 | SPI | periplasmic protein | 0.0079 | 4 | GH28 | 13 | Glycosyl hydrolases family 28; Pectate lyase superfamily protein | Carbohydrate transport and metabolism |
| Fjoh_2111 | A5FI22 | SPI | periplasmic protein | 0.00012 | 3.9 | | | Outer membrane lipoprotein carrier protein LolA | Cell wall/membrane/envelope biogenesis |
| Fjoh_1399 | A5FK36 | SPI | periplasmic protein | 0.0026 | 3.8 | GH13_46 | 3 | Alpha amylase, catalytic domain; Cyclo-malto-dextrinase C-terminal domain; Cyclomaltodextrinase, N-terminal | Carbohydrate transport and metabolism |
| Fjoh_1773 | A5FJ10 | SPI | periplasmic protein | < 0.00010 | 3.8 | Pept_MA | | Peptidase family M1 domain | Amino acid transport and metabolism |
| Fjoh_2780 | A5FG66 | SPI | periplasmic protein | 0.004 | 3.8 | | | Thioredoxin-like | Posttranslational modification, protein turnover, chaperones |
| Fjoh_0636 | A5FM97 | SPI | periplasmic protein | 0.00033 | 3.7 | Pept_SC | | Prolyl oligopeptidase family; WD40-like Beta Propeller Repeat | Amino acid transport and metabolism |
| Fjoh_2040 | A5FIA5 | SPI | periplasmic protein | 0.0016 | 3.7 | GH29 | 7 or 11 | Alpha-L-fucosidase; Alpha-L-fucosidase C-terminal domain | Function unknown |
| Fjoh_4250 | A5FC14 | SPII | periplasm-facing lipoprotein | 0.0097 | 3.7 | GH105 | 26 or 29 | Glycosyl Hydrolase Family 88 | Function unknown |
| Fjoh_1152 | A5FKT5 | SPI | periplasmic protein | 0.0045 | 3.6 | Pept_CA | | Peptidase C1-like family | Amino acid transport and metabolism |
| Fjoh_3112 | A5FF87 | SPI | periplasmic protein | 0.0022 | 3.6 | GH95 | 13 | Glyco_hyd_65N_2; Glyco_hydro_95_C | Carbohydrate transport and metabolism |
| Fjoh_2122 | A5FI17 | SPI | periplasmic protein | 0.0028 | 3.5 | | | Glycosyl hydrolase-like 10 | Function unknown |

| | | | | | | | | | |
|---|---|---|---|---|---|---|---|---|---|
| Fjoh_4900 | A5FA77 | SPI | periplasmic protein | 0.008 | 3.5 | Pept_MA | | Peptidase family M1 domain; Peptidase M1 N-terminal domain | Cell wall/membrane/envelope biogenesis |
| Fjoh_0189 | A5FNI9 | SPI | periplasmic protein | 0.002 | 3.2 | | | Protein of unknown function (DUF3347) | Cell wall/membrane/envelope biogenesis |
| Fjoh_0660 | A5FM68 | SPII | surface-exposed lipoprotein | 0.0041 | 3.2 | | | Tetratricopeptide repeat | Function unknown |
| Fjoh_1993 | A5FIE2 | SPI | periplasmic protein | 0.0034 | 3.2 | GH67 | | Lycosyl hydrolase family 67 C-terminus; Glycosyl hydrolase family 67 middle domain; Glycosyl hydrolase family 67 N-terminus | |
| Fjoh_2928 | A5FFR3 | SPI | periplasmic protein | 0.003 | 3 | Pept_MA | | Peptidase family M1 domain; Peptidase M1 N-terminal domain | Amino acid transport and metabolism |
| Fjoh_1543 | A5FJP3 | SPI | periplasmic protein | 0.00098 | 2.9 | Pept_ME | | Peptidase M16 inactive domain | Function unknown |
| Fjoh_2730 | A5FGC1 | SPI | periplasmic protein | 0.0018 | 2.8 | | | Ankyrin repeats (3 copies) | Function unknown |
| Fjoh_4868 | A5FA98 | SPI | periplasmic protein | 0.0048 | 2.7 | Pept_MA | | Peptidase family M1 domain | Amino acid transport and metabolism |
| Fjoh_0488 | A5FMN9 | SPI | periplasmic protein | 0.00098 | 2.4 | | | Tetratricopeptide repeat | Function unknown |
| Fjoh_1529 | A5FJR3 | SPII | periplasm-facing lipoprotein | 0.0098 | 2.4 | | | Bacterial alpha-2-macroglobulin MG3 domain; MG2 domain; acterial Alpha-2-macroglobulin MG5 domain; Bacterial macroglobulin domain 6; Alpha-2-macroglobulin bait region domain; Alpha-2-macroglobulin family; A-macroglobulin TED domain; Bacterial Alpha-2-macroglobulin MG10 domain | Function unknown |
| Fjoh_4556 | A5FB64 | SPI | periplasmic protein | 0.0014 | 2.4 | GH20 | 30 or 34 or 35 | Glycosyl hydrolase family 20, domain 2; Glycosyl hydrolase family 20, catalytic domain | Carbohydrate transport and metabolism |
| Fjoh_4809 | A5FAG6 | SPI | periplasmic protein | 0.0022 | 2.4 | | | Glycosyl hydrolase-like 10 | Inorganic ion transport and metabolism |
| Fjoh_2749 | A5FGA3 | SPI | Integral/β-barrel protein | 0.00073 | 2.2 | | | Putative auto-transporter adhesin, head GIN domain | Function unknown |
| Fjoh_1190 | A5FKP7 | SPII | periplasm-facing lipoprotein | 0.0064 | 1.5 | Pept_SB | | Subtilase family | Posttranslational modification, protein turnover, chaperones |
| Fjoh_2042 | A5FI94 | SPI | periplasmic protein | 0.00014 | 19 | GH3 | 7 or 11 | Glycosyl hydrolase family 3 N terminal domain; Glycosyl hydrolase family 3 C-terminal domain; Fibronectin type III-like domain | Function unknown |

| | | | | | | | | | |
|---|---|---|---|---|---|---|---|---|---|
| Fjoh_4777 | A5FAJ7 | SPI | periplasmic protein | 0.0085 | 19 | | | | Function unknown |
| Fjoh_1789 | A5FIZ8 | SPI | Integral/β-barrel protein | 0.00036 | 18 | | | | Transcription |
| Fjoh_2959 | A5FFN3 | SPI | periplasmic protein | 0.0032 | 18 | | | | Function unknown |
| Fjoh_0500 | A5FMN5 | SPI | periplasmic protein | 0.00011 | 17 | Pept_SE | | Tetratricopeptide repeat; Beta-lactamase | Defense mechanisms |
| Fjoh_2781 (MenD) | A5FG67 | SPI | periplasmic protein | < 0.00010 | 17 | | | Thiamine pyrophosphate enzyme, N-terminal TPP binding domain; Middle domain of thiamine pyrophosphate; Thiamine pyrophosphate enzyme, C-terminal TPP binding domain | Function unknown |
| Fjoh_1209 | A5FKN5 | SPI | periplasmic protein | 0.00046 | 16 | GH97 | 2 or 5 | Glycosyl-hydrolase 97 C-terminal, oligomerisation; Glycosyl-hydrolase 97 N-terminal; Glycoside hydrolase 97 | Defense mechanisms |
| Fjoh_4501 | A5FBC4 | SPI | periplasmic protein | < 0.00010 | 16 | | | Protein of unknown function (DUF541) | Function unknown |
| Fjoh_0468 | A5FMR6 | SPI | periplasmic protein | 0.0032 | 15 | | | | Function unknown |
| Fjoh_3880 | A5FD23 | SPI | periplasmic protein | < 0.00010 | 15 | GH43_12/CBM91 | 20 or 23 | Glycosyl hydrolases family 43; Beta xylosidase C-terminal Concanavalin A-like domain | Carbohydrate transport and metabolism |
| Fjoh_4603 | A5FB23 | SPI | periplasmic protein | 0.0016 | 15 | Pept_MH | | Peptidase family M20/M25/M40 | Function unknown |
| Fjoh_0271 (MsrA) | A5FNB2 | SPI | periplasmic protein | 0.0014 | 14 | | | Peptide methionine sulfoxide reductase | Posttranslational modification, protein turnover, chaperones |
| Fjoh_3861 | A5FD44 | SPI | periplasmic protein | 0.00075 | 14 | GH3 | 19 or 23 | Glycosyl hydrolase family 3 N terminal domain; Glycosyl hydrolase family 3 C-terminal domain; Fibronectin type III-like domain | Function unknown |
| Fjoh_4086 | A5FCH4 | SPI | periplasmic protein | < 0.00010 | 14 | GH143 | 22 or 25 | | Carbohydrate transport and metabolism |
| Fjoh_4481 | A5FBD2 | SPI | periplasmic protein | 0.00056 | 14 | | | Protein of unknown function (DUF1349) | Function unknown |
| Fjoh_0096 | A5FNT4 | SPI | periplasmic protein | < 0.00010 | 13 | | | Domain of unknown function (DUF4252) | Function unknown |
| Fjoh_0097 | A5FNT5 | SPII | periplasm-facing lipoprotein | 0.0039 | 13 | | | Domain of unknown function (DUF4252) | Function unknown |
| Fjoh_3877 | A5FD33 | SPI | periplasmic protein | 0.0055 | 13 | GH115 | 20 or 23 | Glycosyl hydrolase family 115; Gylcosyl hydrolase family 115 C-terminal domain | Function unknown |
| Fjoh_2714 | A5FGD3 | SPI | periplasmic protein | 0.0021 | 12 | GH92 | 10 or 14 | Glycosyl hydrolase family 92 catalytic domain; Glycosyl | Carbohydrate transport and metabolism |

| | | | | | | | | | |
|---|---|---|---|---|---|---|---|---|---|
| | | | | | | | | hydrolase family 92 N-terminal domain | |
| Fjoh_3473 | A5FE85 | SPI | periplasmic protein | 0.0081 | 12 | | | Calcineurin-like phosphoesterase | Nucleotide transport and metabolism |
| Fjoh_4080 | A5FCI1 | SPI | periplasmic protein | 0.0017 | 12 | GH78 | 22 or 25 | Alpha-L-rhamnosidase N-terminal domain; Bacterial alpha-L-rhamnosidase concanavalin-like domain; Bacterial alpha-L-rhamnosidase 6 hairpin glycosidase domain; Bacterial alpha-L-rhamnosidase C-terminal domain | Carbohydrate transport and metabolism |
| Fjoh_2253 | A5FHP1 | SPI | periplasmic protein | < 0.00010 | 11 | Pept_ME | | Insulinase (Peptidase family M16); Peptidase M16 inactive domain | Function unknown |
| Fjoh_2626 | A5FGM2 | SPI | periplasmic protein | 0.0068 | 11 | Pept_SC | | alpha/beta hydrolase fold | Lipid transport and metabolism |
| Fjoh_2716 | A5FGD5 | SPI | periplasmic protein | 0.0069 | 11 | GH125 | 10 or 14 | Metal-independent alpha-mannosidase (GH125) | Function unknown |
| Fjoh_4915 | A5FA53 | SPI | periplasmic protein | 0.0012 | 11 | | | Uncharacterized protein conserved in bacteria (DUF2141) | Function unknown |
| Fjoh_2038 | A5FIA3 | SPI | periplasmic protein | 0.00049 | 10 | GH2 | 7 or 11 | Beta galactosidase small chain; Glycosyl hydrolases family 2; Glycosyl hydrolases family 2, TIM barrel domain; Glycosyl hydrolases family 2, sugar binding domain; Beta-galactosidase, domain 4 | Carbohydrate transport and metabolism |
| Fjoh_4198 | A5FC62 | SPI | periplasmic protein | 0.0002 | 9.9 | GH2 | 24 or 27 | Glycosyl hydrolases family 2, sugar binding domain; Glycosyl hydrolases family 2; Glycosyl hydrolases family 2, TIM barrel domain | Carbohydrate transport and metabolism |
| Fjoh_3521 | A5FE32 | SPI | periplasmic protein | 0.0013 | 9.8 | GH3 | 16 or 20 | Glycosyl hydrolase family 3 N terminal domain; Glycosyl hydrolase family 3 C-terminal domain; Fibronectin type III-like domain | Function unknown |
| Fjoh_2750 | A5FGA4 | SPI | Integral/β-barrel protein | 0.0002 | 9.5 | | | LamB porin (for maltodextrin uptake) | Function unknown |
| Fjoh_3113 | A5FF88 | SPI | periplasmic protein | 0.00023 | 9.2 | GH51_2 | 13 | Alpha-L-arabinofuranosidase C-terminal domain | Carbohydrate transport and metabolism |
| Fjoh_4097 | A5FCF9 | SPI | periplasmic protein | 0.00015 | 8.8 | GH127 | 22 or 25 | Beta-L-arabinofuranosidase, GH127 catalytic domain; Beta-L-arabinofuranosidase, GH127 middle domain; Glycoside | Function unknown |

| | | | | | | | | | |
|---|---|---|---|---|---|---|---|---|---|
| | | | | | | | | hydrolase family 127 C-terminal domain | |
| Fjoh_0846 | A5FLP6 | SPI | periplasmic protein | 0.0038 | 8.4 | | | Tetratricopeptide repeat | Function unknown |
| Fjoh_2342 | A5FHF7 | SPII | surface-exposed lipoprotein | 0.00011 | 8.4 | | | | Function unknown |
| Fjoh_3873 | A5FD29 | SPI | periplasmic protein | 0.0005 | 8 | GH97 | 20 or 23 | Glycosyl-hydrolase 97 N-terminal; Glycoside hydrolase 97; Glycosyl-hydrolase 97 C-terminal, oligomerisation | Function unknown |
| Fjoh_3817 | A5FD95 | SPI | periplasmic protein | 0.0034 | 7.7 | Pept_SE | | Beta-lactamase | Defense mechanisms |
| Fjoh_4429 | A5FBI0 | SPI | periplasmic protein | 0.00044 | 7.7 | GH97 | 29 or 32 | Glycosyl-hydrolase 97 C-terminal, oligomerisation; Glycosyl-hydrolase 97 N-terminal; Glycoside hydrolase 97 | Defense mechanisms |
| Fjoh_2961 | A5FFN5 | SPI | periplasmic protein | < 0.00010 | 7.6 | Est | | GlcNAc-PI de-N-acetylase | Function unknown |
| Fjoh_1562 | A5FJM0 | SPI | periplasmic protein | 0.0023 | 7.4 | GH51_1 | | Glycosyl hydrolase family 30 TIM-barrel domain | Function unknown |
| Fjoh_3389 | A5FEH0 | SPI | periplasmic protein | < 0.00010 | 7.2 | GH3 | | Glycosyl hydrolase family 3 N terminal domain; Glycosyl hydrolase family 3 C-terminal domain; Fibronectin type III-like domain | Carbohydrate transport and metabolism |
| Fjoh_1541 | A5FJP1 | SPII | periplasm-facing lipoprotein | 0.0098 | 7.1 | | | Heavy-metal-associated domain | Inorganic ion transport and metabolism |
| Fjoh_4995 | A5F9X6 | SPI | periplasmic protein | 0.0064 | 6.7 | | | | Carbohydrate transport and metabolism |
| Fjoh_3194 | A5FF07 | SPI | T9SS C-ter. -sorting domain protein | < 0.00010 | 6.6 | | | Type IX secretion system membrane protein PorP/SprF | Cell motility |
| Fjoh_4808 | A5FAG5 | SPI | periplasmic protein | < 0.00010 | 6 | GH20 | | Glycosyl hydrolase family 20, domain 2; Glycosyl hydrolase family 20, catalytic domain; Chitobiase/beta-hexosaminidase C-terminal domain | Function unknown |
| Fjoh_0778 | A5FLV6 | SPI | periplasmic protein | 0.0072 | 5.8 | GH31_3 | 1 or 4 | Glycosyl hydrolase 31 N-terminal galactose mutarotase-like domain; Glycosyl hydrolases family 31 TIM-barrel domain; Glycosyl hydrolase family 31 C-terminal domain | Carbohydrate transport and metabolism |
| Fjoh_4088 | A5FCH6 | SPI | periplasmic protein | 0.0015 | 5.7 | GH28 | 22 or 25 | Pectate lyase superfamily protein; Glycosyl hydrolases family 28 | Carbohydrate transport and metabolism |

| | | | | | | | | | |
|---|---|---|---|---|---|---|---|---|---|
| Fjoh_1400 | A5FK37 | SPI | periplasmic protein | < 0.00010 | 5.6 | GH97 | 3 | Glycosyl-hydrolase 97 C-terminal, oligomerisation; Glycosyl-hydrolase 97 N-terminal; Glycoside hydrolase 97 | Carbohydrate transport and metabolism |
| Fjoh_0119 | A5FNQ4 | SPI | periplasmic protein | 0.00026 | 5.3 | Pept_MO | | Peptidase family M23 | Cell wall/membrane/envelope biogenesis |
| Fjoh_4779 | A5FAI4 | SPII | periplasm-facing lipoprotein | 0.00054 | 5.3 | | | | RNA processing and modification |
| Fjoh_1419 | A5FK22 | SPI | periplasmic protein | < 0.00010 | 5.2 | Pept_SC | | Dipeptidyl peptidase IV (DPP IV) N-terminal region; Prolyl oligopeptidase family | Function unknown |
| Fjoh_2341 | A5FHF6 | SPI | periplasmic protein | 0.0017 | 5.2 | Est | | Putative esterase | Function unknown |
| Fjoh_2041 | A5FI93 | SPI | periplasmic protein | 0.0055 | 5.1 | GH92 | 7 or 11 | Glycosyl hydrolase family 92 N-terminal domain; Glycosyl hydrolase family 92 catalytic domain | Carbohydrate transport and metabolism |
| Fjoh_3800 | A5FDB0 | SPI | periplasmic protein | 0.0036 | 5.1 | | 18 or 22 | Domain of unknown function (DUF5118); Domain of unknown function (DUF5117); Met-zincin | Posttranslational modification, protein turnover, chaperones |
| Fjoh_1430 | A5FK03 | SPI | periplasmic protein | 0.00028 | 5 | Pept_SK | | C-terminal domain of tail specific protease (DUF3340); PDZ domain; Peptidase family S41; Tail specific protease N-terminal domain | Cell wall/membrane/envelope biogenesis |
| Fjoh_3874 | A5FD30 | SPI | periplasmic protein | 0.0017 | 4.9 | GH3 | 20 or 23 | Glycosyl hydrolase family 3 N terminal domain; Glycosyl hydrolase family 3 C-terminal domain; Fibronectin type III-like domain | Carbohydrate transport and metabolism |
| Fjoh_2358 | A5FHD9 | SPII | periplasm-facing lipoprotein | < 0.00010 | 4.8 | | | Glycosyl hydrolase family 92 catalytic domain; Glycosyl hydrolase family 92 N-terminal domain | Amino acid transport and metabolism |
| Fjoh_1191 | A5FKP8 | SPI | periplasmic protein | < 0.00010 | 4.7 | Pept_MA | | Peptidase family M1 domain | Function unknown |
| Fjoh_1564 | A5FJM2 | SPI | periplasmic protein | 0.0043 | 4.6 | GH30_1 | 4 or 7 | Fibronectin type III-like domain; Glycosyl hydrolase family 3 N terminal domain; Glycosyl hydrolase family 3 C-terminal domain | Function unknown |
| Fjoh_3437 | A5FEB2 | SPI | periplasmic protein | < 0.00010 | 4.4 | Pept_MA | | Peptidase family M1 domain | Amino acid transport and metabolism |

| | | | | | | | | | |
|---|---|---|---|---|---|---|---|---|---|
| Fjoh_0200 | A5FNI7 | SPII | periplasm-facing lipoprotein | 0.00035 | 4.2 | | | | Function unknown |
| Fjoh_0415 | A5FMW7 | SPI | periplasmic protein | 0.0002 | 4.2 | | | Alpha-2-macroglobulin family; MG2 domain; Bacterial Alpha-2-macroglobulin MG10 domain | Function unknown |
| Fjoh_3392 | A5FEF5 | SPI | periplasmic protein | 0.0069 | 4.1 | GH3 | | Fibronectin type III-like domain; Glycosyl hydrolase family 3 N terminal domain; Glycosyl hydrolase family 3 C-terminal domain | Carbohydrate transport and metabolism |
| Fjoh_0023 | A5FP14 | SPI | periplasmic protein | < 0.00010 | 4 | | | Type I phosphodiesterase / nucleotide pyrophosphatase | Function unknown |
| Fjoh_0246 | A5FND2 | SPI | periplasmic protein | 0.0026 | 4 | CBM50/Est | | GDSL-like Lipase/Acylhydrolase family; LysM domain | Amino acid transport and metabolism; Cell wall/membrane/envelope biogenesis |
| Fjoh_2181 | A5FHV4 | SPI | periplasmic protein | 0.0003 | 4 | | | Monomeric isocitrate dehydrogenase | Energy production and conversion |
| Fjoh_2715 | A5FGD4 | SPI | periplasmic protein | 0.00087 | 4 | GH92 | 10 or 14 | Glycosyl hydrolase family 92 N-terminal domain; Glycosyl hydrolase family 92 catalytic domain | Carbohydrate transport and metabolism |
| Fjoh_3114 | A5FF89 | SPI | periplasmic protein | 0.0079 | 4 | GH28 | 13 | Glycosyl hydrolases family 28; Pectate lyase superfamily protein | Carbohydrate transport and metabolism |
| Fjoh_2111 | A5FI22 | SPI | periplasmic protein | 0.00012 | 3.9 | | | Outer membrane lipoprotein carrier protein LolA | Cell wall/membrane/envelope biogenesis |
| Fjoh_1399 | A5FK36 | SPI | periplasmic protein | 0.0026 | 3.8 | GH13_46 | 3 | Alpha amylase, catalytic domain; Cyclo-malto-dextrinase C-terminal domain; Cyclomaltodextrinase, N-terminal | Carbohydrate transport and metabolism |
| Fjoh_1773 | A5FJ10 | SPI | periplasmic protein | < 0.00010 | 3.8 | Pept_MA | | Peptidase family M1 domain | Amino acid transport and metabolism |
| Fjoh_2780 | A5FG66 | SPI | periplasmic protein | 0.004 | 3.8 | | | Thioredoxin-like | Posttranslational modification, protein turnover, chaperones |
| Fjoh_0636 | A5FM97 | SPI | periplasmic protein | 0.00033 | 3.7 | Pept_SC | | Prolyl oligopeptidase family; WD40-like Beta Propeller Repeat | Amino acid transport and metabolism |
| Fjoh_2040 | A5FIA5 | SPI | periplasmic protein | 0.0016 | 3.7 | GH29 | 7 or 11 | Alpha-L-fucosidase; Alpha-L-fucosidase C-terminal domain | Function unknown |
| Fjoh_4250 | A5FC14 | SPII | periplasm-facing lipoprotein | 0.0097 | 3.7 | GH105 | 26 or 29 | Glycosyl Hydrolase Family 88 | Function unknown |

| | | | | | | | | | |
|---|---|---|---|---|---|---|---|---|---|
| Fjoh_1152 | A5FKT5 | SPI | periplasmic protein | 0.0045 | 3.6 | Pept_CA | | Peptidase C1-like family | Amino acid transport and metabolism |
| Fjoh_3112 | A5FF87 | SPI | periplasmic protein | 0.0022 | 3.6 | GH95 | 13 | Glyco_hyd_65N_2; Glyco_hydro_95_C | Carbohydrate transport and metabolism |
| Fjoh_2122 | A5FI17 | SPI | periplasmic protein | 0.0028 | 3.5 | | | Glycosyl hydrolase-like 10 | Function unknown |
| Fjoh_4900 | A5FA77 | SPI | periplasmic protein | 0.008 | 3.5 | Pept_MA | | Peptidase family M1 domain; Peptidase M1 N-terminal domain | Cell wall/membrane/envelope biogenesis |
| Fjoh_0189 | A5FNI9 | SPI | periplasmic protein | 0.002 | 3.2 | | | Protein of unknown function (DUF3347) | Cell wall/membrane/envelope biogenesis |
| Fjoh_0660 | A5FM68 | SPII | surface-exposed lipoprotein | 0.0041 | 3.2 | | | Tetratricopeptide repeat | Function unknown |
| Fjoh_1993 | A5FIE2 | SPI | periplasmic protein | 0.0034 | 3.2 | GH67 | | Lycosyl hydrolase family 67 C-terminus; Glycosyl hydrolase family 67 middle domain; Glycosyl hydrolase family 67 N-terminus | |
| Fjoh_2928 | A5FFR3 | SPI | periplasmic protein | 0.003 | 3 | Pept_MA | | Peptidase family M1 domain; Peptidase M1 N-terminal domain | Amino acid transport and metabolism |
| Fjoh_1543 | A5FJP3 | SPI | periplasmic protein | 0.00098 | 2.9 | Pept_ME | | Peptidase M16 inactive domain | Function unknown |
| Fjoh_2730 | A5FGC1 | SPI | periplasmic protein | 0.0018 | 2.8 | | | Ankyrin repeats (3 copies) | Function unknown |
| Fjoh_4868 | A5FA98 | SPI | periplasmic protein | 0.0048 | 2.7 | Pept_MA | | Peptidase family M1 domain | Amino acid transport and metabolism |
| Fjoh_0488 | A5FMN9 | SPI | periplasmic protein | 0.00098 | 2.4 | | | Tetratricopeptide repeat | Function unknown |
| Fjoh_1529 | A5FJR3 | SPII | periplasm-facing lipoprotein | 0.0098 | 2.4 | | | Bacterial alpha-2-macroglobulin MG3 domain; MG2 domain; acterial Alpha-2-macroglobulin MG5 domain; Bacterial macroglobulin domain 6; Alpha-2-macroglobulin bait region domain; Alpha-2-macroglobulin family; A-macroglobulin TED domain; Bacterial Alpha-2-macroglobulin MG10 domain | Function unknown |
| Fjoh_4556 | A5FB64 | SPI | periplasmic protein | 0.0014 | 2.4 | GH20 | 30 or 34 or 35 | Glycosyl hydrolase family 20, domain 2; Glycosyl hydrolase family 20, catalytic domain | Carbohydrate transport and metabolism |
| Fjoh_4809 | A5FAG6 | SPI | periplasmic protein | 0.0022 | 2.4 | | | Glycosyl hydrolase-like 10 | Inorganic ion transport and metabolism |
| Fjoh_2749 | A5FGA3 | SPI | Integral/β-barrel protein | 0.00073 | 2.2 | | | Putative auto-transporter adhesin, head GIN domain | Function unknown |

| | | | | | | | | | |
|---|---|---|---|---|---|---|---|---|---|
| Fjoh_1190 | A5FKP7 | SPII | periplasm-facing lipoprotein | 0.0064 | 1.5 | Pept_SB | | Subtilase family | Posttranslational modification, protein turnover, chaperones |
| Fjoh_3203 | A5FEZ6 | SPI | T9SS C-ter. -sorting domain protein | 0.00016 | 0 | GH87 | | Bacterial Ig-like domain (group 2); Secretion system C-terminal sorting domain (type A) | Cell motility |
| Fjoh_4327 | A5FBT1 | SPII | surface-exposed lipoprotein | 0.0052 | 0 | GH99 | 27 or 30 | Glycosyl hydrolase family 99 | Function unknown |
| Fjoh_1231 | A5FKK5 | SPI | T9SS C-ter. -sorting domain protein | 0.0016 | 0.02 | PL1/CBM77 | | Carbohydrate binding module 77; Pectate lyase; Secretion system C-terminal sorting domain | Carbohydrate transport and metabolism |
| Fjoh_1606 | A5FJI1 | SPII | surface-exposed lipoprotein | 0.0051 | 0.02 | | | Lipocalin-like domain | Function unknown |
| Fjoh_4175 | A5FC89 | SPI | T9SS C-ter. -sorting domain protein | 0.0073 | 0.02 | GH18/CBM 6 | | Carbohydrate binding module (family 6); Glycosyl hydrolases family 18; Secretion system C-terminal sorting domain (type A) | Carbohydrate transport and metabolism |
| Fjoh_1408 | A5FK34 | SPI | T9SS C-ter. -sorting domain protein | 0.00014 | 0.05 | CBM98/CB M48/GH13_ 47 | 3 or 6 | Alpha amylase, catalytic domain; Secretion system C-terminal sorting domain | Carbohydrate transport and metabolism |
| Fjoh_1771 | A5FJ21 | SPI | periplasmic protein | 0.0055 | 0.05 | | | | Function unknown |
| Fjoh_0185 | A5FNK1 | SPI | integral/β-barrel protein | 0.0057 | 0.06 | | | TonB dependent receptor-like, beta-barrel; TonB-dependent Receptor Plug Domain; CarboxypepD_reg-like domain | Cell wall/membrane/envelope biogenesis |
| Fjoh_1122 | A5FKW4 | SPII | surface-exposed lipoprotein | 0.0044 | 0.06 | EPI | | Aldose 1-epimerase | Carbohydrate transport and metabolism |
| Fjoh_3296 | A5FEQ3 | SPI | T9SS C-ter. -sorting domain protein | < 0.00010 | 0.06 | | | Secretion system C-terminal sorting domain (type C) | Cell motility |
| Fjoh_1211 | A5FKM2 | SPII | surface-exposed lipoprotein | 0.0059 | 0.08 | | 2 or 5 | SusE outer membrane protein | Function unknown |
| Fjoh_4177 | A5FC91 | SPI | T9SS C-ter. -sorting domain protein | < 0.00010 | 0.08 | GH16_3/CB M102/CBM 32 | | F5/8 type C domain; Glycosyl hydrolases family 16; PKD-like domain; Secretion system C-terminal sorting domain (type A) | Carbohydrate transport and metabolism |
| Fjoh_0774 | A5FLW7 | SPII | surface-exposed lipoprotein | 0.00068 | 0.09 | GH5_4 | 1 or 4 | BACON domain; Cellulase (glycosyl hydrolase family 5) | Carbohydrate transport and metabolism |
| Fjoh_1269 | A5FKG0 | SPI | T9SS C-ter. -sorting domain protein | 0.0058 | 0.09 | | | SprB repeat; Secretion system C-terminal sorting domain | Posttranslational modification, protein turnover, chaperones |
| Fjoh_0248 | A5FND4 | SPII | surface-exposed lipoprotein | 0.0096 | 0.1 | | | HmuY protein | Function unknown |
| Fjoh_0561 | A5FMH4 | SPII | surface-exposed lipoprotein | 0.0047 | 0.1 | | | | Function unknown |
| Fjoh_0618 | A5FMB4 | SPI | integral/β-barrel protein | 0.003 | 0.1 | | | Bacterial type II and III secretion system protein | Intracellular trafficking, secretion, and vesicular transport |
| Fjoh_0764 | A5FLX3 | SPII | surface-exposed lipoprotein | 0.0013 | 0.1 | | | Protein of unknown function (DUF4876) | Function unknown |

| | | | | | | | | | |
|---|---|---|---|---|---|---|---|---|---|
| Fjoh_1208 | A5FKN4 | SPI | T9SS C-ter. -sorting domain protein | 0.00024 | 0.1 | GH13/CBM 26 | 2 or 5 | Alpha amylase, catalytic domain; Secretion system C-terminal sorting domain (type A); Bacterial Ig-like domain (group 2); Starch-binding module 26 | Cell motility |
| Fjoh_1905 | A5FIN0 | SPI | T9SS C-ter. -sorting domain protein | 0.00064 | 0.1 | GH30_3/CB M13 | | Glycosyl hydrolase family 30 TIM-barrel domain; Glycosyl hydrolase family 30 beta sandwich domain; Ricin-type beta-trefoil lectin domain-like; Secretion system C-terminal sorting domain | Carbohydrate transport and metabolism |
| Fjoh_2057 | A5FI81 | SPII | periplasm-facing lipoprotein | 0.0087 | 0.1 | | | | Function unknown |
| Fjoh_2498 | A5FGZ4 | SPI | integral/β-barrel protein | 0.00074 | 0.1 | | | Domain of unknown function (DUF6048) | Function unknown |
| Fjoh_2566 | A5FGT3 | SPII | surface-exposed lipoprotein | 0.00018 | 0.1 | PL1 | | Pectate lyase | Carbohydrate transport and metabolism |
| Fjoh_3108 | A5FF99 | SPI | T9SS C-ter. -sorting domain protein | 0.00032 | 0.1 | | | Secretion system C-terminal sorting domain (type A) | Cell motility |
| Fjoh_3180 | A5FF18 | SPII | surface-exposed lipoprotein | 0.0025 | 0.1 | | | | Function unknown |
| Fjoh_3246 | A5FEV3 | SPI | T9SS C-ter. -sorting domain protein | 0.0001 | 0.1 | | | CARDB; Secretion system C-terminal sorting domain (type A) | Function unknown |
| Fjoh_3290 | A5FER3 | SPII | surface-exposed lipoprotein | 0.0036 | 0.1 | | | | Function unknown |
| Fjoh_3418 | A5FEE4 | SPII | surface-exposed lipoprotein | 0.0032 | 0.1 | | | | Function unknown |
| Fjoh_3818 | A5FD96 | SPI | T9SS C-ter. -sorting domain protein | 0.0024 | 0.1 | | | Secretion system C-terminal sorting domain (type C) | Cell motility |
| Fjoh_4095 | A5FCH0 | SPI | integral/β-barrel protein | 0.0026 | 0.1 | SusC | 22 or 25 | CarboxypepD_reg-like domain; TonB-dependent receptor, plug domain | Inorganic ion transport and metabolism |
| Fjoh_4174 | A5FC88 | SPI | T9SS C-ter. -sorting domain protein | 0.0031 | 0.1 | CBM13/CB M6 | | Carbohydrate binding module (family 6); Ricin-type beta-trefoil lectin domain-like; Secretion system C-terminal sorting domain | Carbohydrate transport and metabolism |
| Fjoh_4176 | A5FC90 | SPI | T9SS C-ter. -sorting domain protein | 0.0014 | 0.1 | GH64/CBM 13/CBM6 | | Carbohydrate binding module (family 6); Ricin-type beta-trefoil lectin domain-like; Secretion system C-terminal sorting domain; Beta-1,3-glucanase | Carbohydrate transport and metabolism |
| Fjoh_4432 | A5FBI3 | SPII | surface-exposed lipoprotein | 0.0021 | 0.1 | | 29 or 32 | SusE outer membrane protein; Outer membrane protein SusF_SusE | Function unknown |

| | | | | | | | | | |
|---|---|---|---|---|---|---|---|---|---|
| Fjoh_4434 | A5FBI5 | SPI | integral/β-barrel protein | 0.0028 | 0.1 | SusC | 29 or 32 | CarboxypepD_reg-like domain; TonB-dependent receptor, plug domain | Inorganic ion transport and metabolism |
| Fjoh_4720 | A5FAP7 | SPII | surface-exposed lipoprotein | 0.00075 | 0.1 | | | | Function unknown |
| Fjoh_4951 | A5FA22 | SPI | integral/β-barrel protein | 0.0027 | 0.1 | SusC | 33 or 39 | CarboxypepD_reg-like domain; TonB dependent receptor; TonB-dependent receptor, plug domain | Coenzyme transport and metabolism |
| Fjoh_0093 | A5FNT1 | SPII | surface-exposed lipoprotein | 0.00014 | 0.2 | Pept_SK | | Peptidase S41 N-terminal domain; Peptidase family S41 | Cell wall/membrane/envelope biogenesis |
| Fjoh_0183 | A5FNJ9 | SPII | surface-exposed lipoprotein | 0.0045 | 0.2 | | 1 | Domain of unknown function (DUF4957); Domain of unknown function (DUF5123); Fibronectin type III domain | Function unknown |
| Fjoh_0454 | A5FMS1 | SPII | periplasm-facing lipoprotein | 0.0035 | 0.2 | | | Peptidase family M28 | Function unknown |
| Fjoh_0621 | A5FMB7 | SPII | surface-exposed lipoprotein | 0.00018 | 0.2 | | | | Function unknown |
| Fjoh_0798 | A5FLU4 | SPI | T9SS C-ter. -sorting domain protein | 0.0016 | 0.2 | | | Proprotein convertase P-domain; Metallo-peptidase family M12B Reprolysin-like; Secretion system C-terminal sorting domain (type A) | Posttranslational modification, protein turnover, chaperones |
| Fjoh_0806 | A5FLT6 | SPI | periplasmic protein | 0.00011 | 0.2 | | | PA14 domain | Cell motility |
| Fjoh_1189 | A5FKP6 | SPI | T9SS C-ter. -sorting domain protein | 0.00024 | 0.2 | | | Concanavalin A-like lectin/glucanases superfamily; Regulator of chromosome condensation (RCC1) repeat; SprB repeat; Secretion system C-terminal sorting domain (type A); Ig-like domain CHU_C associated | Cytoskeleton |
| Fjoh_1393 | A5FK44 | SPII | surface-exposed lipoprotein | 0.0018 | 0.2 | | | | Function unknown |
| Fjoh_1561 | A5FJL9 | SPII | surface-exposed lipoprotein | 0.009 | 0.2 | SusD | 4 or 7 | SusD family; Starch-binding associating with outer membrane | Carbohydrate transport and metabolism |
| Fjoh_1698 | A5FJ83 | SPII | surface-exposed lipoprotein | < 0.00010 | 0.2 | | | | Function unknown |
| Fjoh_1874 | A5FIR7 | SPII | surface-exposed lipoprotein | 0.00063 | 0.2 | | | Domain of unknown function (DUF5074) | Function unknown |
| Fjoh_2150 | A5FHZ4 | SPI | T9SS C-ter. -sorting domain protein | 0.0043 | 0.2 | | | Secretion system C-terminal sorting domain | Energy production and conversion |
| Fjoh_2376 | A5FHB3 | SPII | surface-exposed lipoprotein | 0.0075 | 0.2 | | | | Function unknown |
| Fjoh_2434 | A5FH59 | SPII | surface-exposed lipoprotein | < 0.00010 | 0.2 | GH16_3/CBM103 | 9 or 13 | SusD family | Carbohydrate transport and metabolism |
| Fjoh_2466 | A5FH32 | SPI | integral/β-barrel protein | < 0.00010 | 0.2 | | | TonB dependent receptor-like, beta-barrel; TonB-dependent Receptor | Cell wall/membrane/envelope biogenesis |

| | | | | | | | | | |
|---|---|---|---|---|---|---|---|---|---|
| | | | | | | | | Plug Domain; CarboxypepD_reg-like domain | |
| Fjoh_2510 | A5FGY9 | SPII | surface-exposed lipoprotein | 0.0017 | 0.2 | | | | Function unknown |
| Fjoh_3247 | A5FEV4 | SPI | T9SS C-ter. -sorting domain protein | 0.00042 | 0.2 | | | Dockerin type I domain; Leucine Rich Repeat; Secretion system C-terminal sorting domain | Function unknown |
| Fjoh_3478 | A5FE73 | SPI | T9SS C-ter. -sorting domain protein | 0.0023 | 0.2 | | | CHU_C Type IX secretion signal domain (type A); Domain of unknown function DUF11; Ig-like domain CHU_C associated | Cell wall/membrane/envelope biogenesis |
| Fjoh_3525 | A5FE36 | SPI | integral/β-barrel protein | < 0.00010 | 0.2 | SusC | 16 or 20 | CarboxypepD_reg-like domain; TonB dependent receptor; TonB-dependent receptor, plug domain | Inorganic ion transport and metabolism |
| Fjoh_3881 | A5FD24 | SPII | surface-exposed lipoprotein | 0.0045 | 0.2 | SusD | | Starch-binding associating with outer membrane; SusD family | Amino acid transport and metabolism |
| Fjoh_4221 | A5FC34 | SPI | integral/β-barrel protein | < 0.00010 | 0.2 | | | CarboxypepD_reg-like domain; TonB dependent receptor; TonB-dependent receptor, plug domain | Energy production and conversion |
| Fjoh_4555 (ChiA) | A5FB63 | SPI | T9SS C-ter. -sorting domain protein | 0.00037 | 0.2 | GH18 | 30 or 34 or 35 | Bacterial Ig domain; Carboxypeptidase regulatory-like domain; Glycosyl hydrolases family 18; Secretion system C-terminal sorting domain (type C) | Carbohydrate transport and metabolism |
| Fjoh_4902 | A5FA64 | SPI | integral/β-barrel protein | 0.0012 | 0.2 | | | Outer membrane efflux protein | Cell wall/membrane/envelope biogenesis |
| Fjoh_5007 | A5F9V9 | SPI | periplasmic protein | 0.0035 | 0.2 | | | Di-haem cytochrome c peroxidase | Energy production and conversion |
| Fjoh_0151 | A5FNN3 | SPII | surface-exposed lipoprotein | 0.009 | 0.3 | | | Domain of unknown function (DUF4249) | Function unknown |
| Fjoh_0736 | A5FLZ8 | SPI | integral/β-barrel protein | 0.0014 | 0.3 | | | CarboxypepD_reg-like domain; TonB dependent receptor; TonB-dependent receptor, plug domain | Inorganic ion transport and metabolism |
| Fjoh_0808 (RemA) | A5FLS4 | SPI | Integral/β-barrel protein | 0.0045 | 0.3 | | | Galactose binding lectin domain | Defense mechanisms |
| Fjoh_0821 | A5FLQ6 | SPI | integral/β-barrel protein | 0.0024 | 0.3 | SusC | | CarboxypepD_reg-like domain; TonB dependent receptor; TonB-dependent receptor, plug domain | Inorganic ion transport and metabolism |
| Fjoh_0823 | A5FLQ8 | SPII | surface-exposed lipoprotein | 0.00035 | 0.3 | | | Domain of unknown function (DUF4270) | Function unknown |
| Fjoh_0886 | A5FLJ5 | SPI | T9SS C-ter. -sorting domain protein | 0.0028 | 0.3 | Pept_MA | | Fibronectin type III domain; Fungalysin/Thermolysin Propeptide Motif; GEVED domain; Thermolysin metallopeptidase, catalytic domain; Thermolysin metallopeptidase, alpha-helical | Amino acid transport and metabolism |

| | | | | | | | | | |
|---|---|---|---|---|---|---|---|---|---|
| | | | | | | | | domain; Secretion system C-terminal sorting domain | |
| Fjoh_0928 | A5FLE9 | SPI | integral/β-barrel protein | 0.00087 | 0.3 | SusC | | CarboxypepD_reg-like domain; TonB dependent receptor; TonB-dependent receptor, plug domain | Inorganic ion transport and metabolism |
| Fjoh_1022 | A5FL64 | SPI | T9SS C-ter. -sorting domain protein | 0.0013 | 0.3 | GH8 | | Secretion system C-terminal sorting domain (type A); Glycosyl hydrolases family 8 | Carbohydrate transport and metabolism |
| Fjoh_1067 | A5FL26 | SPI | periplasmic protein | 0.00053 | 0.3 | Pept_MO | | Peptidase family M23 | Cell cycle control, cell division, chromosome partitioning |
| Fjoh_1188 | A5FKP5 | SPI | T9SS C-ter. -sorting domain protein | 0.003 | 0.3 | | | Secretion system C-terminal sorting domain (type A) | Cell cycle control, cell division, chromosome partitioning |
| Fjoh_1311 | A5FKD0 | SPI | integral/β-barrel protein | 0.00012 | 0.3 | | | Outer membrane protein beta-barrel domain | Cell wall/membrane/envelope biogenesis |
| Fjoh_1405 | A5FK31 | SPI | integral/β-barrel protein | 0.0016 | 0.3 | SusC | 3 or 6 | CarboxypepD_reg-like domain; TonB dependent receptor; TonB-dependent receptor, plug domain | Inorganic ion transport and metabolism |
| Fjoh_1410 | A5FK25 | SPII | surface-exposed lipoprotein | 0.0061 | 0.3 | | | PKD-like domain; Domain of unknown function (DUF5074) | Signal transduction mechanisms |
| Fjoh_2321 | A5FHI0 | SPI | integral/β-barrel protein | 0.00014 | 0.3 | | | Family of unknown function (DUF5723); OmpA family; Thrombospondin type 3 repeat | Cell wall/membrane/envelope biogenesis |
| Fjoh_2433 | A5FH58 | SPII | surface-exposed lipoprotein | 0.0075 | 0.3 | CBM102 | 9 or 13 | | Carbohydrate transport and metabolism |
| Fjoh_2499 | A5FGZ0 | SPII | surface-exposed lipoprotein | 0.0076 | 0.3 | | | Domain of unknown function (DUF4249) | Function unknown |
| Fjoh_2614 | A5FGM9 | SPI | periplasmic protein | 0.0054 | 0.3 | | | | Function unknown |
| Fjoh_2960 | A5FFN4 | SPII | surface-exposed lipoprotein | 0.0029 | 0.3 | | | LVIVD repeat | Function unknown |
| Fjoh_3307 | A5FEP9 | SPII | surface-exposed lipoprotein | 0.0037 | 0.3 | | | Glucose / Sorbosone dehydrogenase | Carbohydrate transport and metabolism |
| Fjoh_3324 | A5FEN6 | SPI | T9SS C-ter. -sorting domain protein | 0.00054 | 0.3 | CBM6 | | Glucose / Sorbosone dehydrogenase; Carbohydrate binding module (family 6); PKD domain; Secretion system C-terminal sorting domain | Carbohydrate transport and metabolism |
| Fjoh_3882 | A5FD25 | SPI | integral/β-barrel protein | 0.00043 | 0.3 | | | CarboxypepD_reg-like domain; TonB dependent receptor; TonB-dependent receptor, plug domain | Cell wall/membrane/envelope biogenesis; Intracellular trafficking, secretion, and vesicular transport |
| Fjoh_4558 | A5FB66 | SPII | surface-exposed lipoprotein | 0.00077 | 0.3 | SusD | 30 or 34 or 35 | SusD and RagB outer membrane lipoprotein | Function unknown |
| Fjoh_4600 | A5FB20 | SPI | periplasmic protein | 0.0074 | 0.3 | | | | Function unknown |

| | | | | | | | | | |
|---|---|---|---|---|---|---|---|---|---|
| Fjoh_4814 | A5FAF5 | SPI | integral/β-barrel protein | 0.0024 | 0.3 | SusC | 32 or 38 | CarboxypepD_reg-like domain; TonB dependent receptor; TonB-dependent receptor, plug domain | Inorganic ion transport and metabolism |
| Fjoh_4815 | A5FAF6 | SPII | surface-exposed lipoprotein | 0.0091 | 0.3 | SusD | 32 or 38 | SusD family; Starch-binding associating with outer membrane | Function unknown |
| Fjoh_4819 | A5FAE4 | SPII | periplasm-facing lipoprotein | 0.0076 | 0.3 | Pept_SE/GH 3 | 32 or 38 | Beta-lactamase; Glycosyl hydrolase family 3 C-terminal domain; Glycosyl hydrolase family 3 N terminal domain | Carbohydrate transport and metabolism |
| Fjoh_5008 | A5F9W0 | SPI | integral/β-barrel protein | 0.0012 | 0.3 | | | | Cell wall/membrane/envelope biogenesis |
| Fjoh_0074 | A5FNW0 | SPI | T9SS C-ter. -sorting domain protein | 0.00078 | 0.4 | | | Secretion system C-terminal sorting domain (type A) | Function unknown |
| Fjoh_0411 | A5FMX9 | SPI | periplasmic protein | 0.00093 | 0.4 | | | | Function unknown |
| Fjoh_1407 | A5FK33 | SPII | surface-exposed lipoprotein | 0.0011 | 0.4 | | 3 or 6 | SusE outer membrane protein; Outer membrane protein SusF_SusE | Function unknown |
| Fjoh_1778 | A5FJ15 | SPI | periplasmic protein | 0.0066 | 0.4 | | | YceI-like domain | Function unknown |
| Fjoh_1873 | A5FIR6 | SPII | surface-exposed lipoprotein | 0.0027 | 0.4 | | | PKD-like domain; Domain of unknown function | Function unknown |
| Fjoh_3417 | A5FEE3 | SPII | surface-exposed lipoprotein | 0.00025 | 0.4 | | | | Function unknown |
| Fjoh_3440 | A5FEB5 | SPI | integral/β-barrel protein | 0.0086 | 0.4 | | | Protein of unknown function (DUF3575) | Function unknown |
| Fjoh_4761 | A5FAL1 | SPII | periplasm-facing lipoprotein | 0.00087 | 0.4 | | | Copper/zinc superoxide dismutase (SODC) | Inorganic ion transport and metabolism |
| Fjoh_4941 | A5FA29 | SPI | Integral/β-barrel protein | 0.00064 | 0.4 | | | Outer membrane protein transport protein (OMPP1/FadL/TodX) | Lipid transport and metabolism |
| Fjoh_0237 | A5FNF3 | SPI | Not assigned | 0.0045 | 0.5 | | | PQQ-like domain | Function unknown |
| Fjoh_0353 | A5FN31 | SPI | Not assigned | 0.0063 | 0.5 | | | Polysaccharide biosynthesis/export protein; SLBB domain | Cell wall/membrane/envelope biogenesis |
| Fjoh_0403 | A5FMY7 | SPI | Integral/β-barrel protein | 0.0046 | 0.5 | SusC | 2 | CarboxypepD_reg-like domain; TonB dependent receptor; TonB-dependent receptor, plug domain | Inorganic ion transport and metabolism |
| Fjoh_0978 | A5FLA7 | SPI | Integral/β-barrel protein | 0.00025 | 0.5 | | | Type IX secretion system membrane protein PorP/SprF | Cell wall/membrane/envelope biogenesis |
| Fjoh_1557 (GldJ) | A5FJM9 | SPII | periplasm-facing lipoprotein | 0.00028 | 0.5 | | | Sulfatase-modifying factor enzyme 1 | Function unknown |
| Fjoh_2621 | A5FGL7 | SPII | periplasm-facing lipoprotein | 0.0047 | 0.5 | | | 3-keto-disaccharide hydrolase | Function unknown |

**Supplementary Table S3**. **TAM homologs in several Bacteroidetes species.** Homologs identified by DELTA-BLAST [7] search using TamA (WP_012022494.1), TamL (WP_012023543.1), TamL2 (WP_012023976.1), TamB (WP_012026557.1) and TamB2 (WP_012023975.1) sequences of *F. johnsoniae* as queries (E value ≤ 0.001).

| Query | Species | RefSeq assembly | Protein accession | % ID | E value |
|---|---|---|---|---|---|
| TamL | *Bacteroides fragilis* | GCF_000025985.1 | WP_005787690.1 | 27.6 | 4.71E-77 |
| TamB | *Bacteroides fragilis* | GCF_000025985.1 | WP_010993036.1 | 23.5 | 1.74E-90 |
| TamL2 | *Bacteroides fragilis* | GCF_000025985.1 | WP_005799192.1 | 30.4 | 2.94E-104 |
| TamB2 | *Bacteroides fragilis* | GCF_000025985.1 | WP_010993654.1 | 23.4 | 3.98E-81 |
| TamL | *Bacteroides ovatus* | GCF_001314995.1 | WP_004295889.1 | 38 | 1.54E-39 |
| TamB | *Bacteroides ovatus* | GCF_001314995.1 | WP_004295986.1 | 22.6 | 2.82E-79 |
| TamL2 | *Bacteroides ovatus* | GCF_001314995.1 | WP_004299936.1 | 30.7 | 3.78E-107 |
| TamB2 | *Bacteroides ovatus* | GCF_001314995.1 | WP_004299935.1 | 24.8 | 4.06E-38 |
| TamL | *Bacteroides thetaioataomicron* | GCF_014131755.1 | WP_011107463.1 | 27.8 | 2.78E-74 |
| TamB | *Bacteroides thetaioataomicron* | GCF_014131755.1 | WP_011107519.1 | 23.9 | 4.1E-83 |
| TamL2 | *Bacteroides thetaioataomicron* | GCF_014131755.1 | WP_008766928.1 | 29.7 | 2.86E-99 |
| TamB2 | *Bacteroides thetaioataomicron* | GCF_014131755.1 | WP_162303127.1 | 23.5 | 1.52E-38 |
| TamA | *Bergeyella zoohelcum* | GCF_000301075.1 | WP_245946064.1 | 27.6 | 8.13E-41 |
| TamL | *Bergeyella zoohelcum* | GCF_000301075.1 | WP_245946060.1 | 29.6 | 8.71E-100 |
| TamB | *Bergeyella zoohelcum* | GCF_000301075.1 | WP_002663326.1 | 23.6 | 9.04E-101 |
| TamL2 | *Bergeyella zoohelcum* | GCF_000301075.1 | WP_002661987.1 | 50.9 | <1.49E-177 |
| TamB2 | *Bergeyella zoohelcum* | GCF_000301075.1 | WP_002661989.1 | 40.7 | <1.49E-177 |
| TamL | *Capnocytophaga canimorsus* | GCF_000220625.1 | WP_042002088.1 | 43.8 | <1.49E-177 |
| TamB | *Capnocytophaga canimorsus* | GCF_000220625.1 | WP_095900346.1 | 37.8 | <1.49E-177 |
| TamL | *Capnocytophaga canis* | GCF_000827555.1 | WP_042345225.1 | 43.1 | <1.49E-177 |
| TamB | *Capnocytophaga canis* | GCF_000827555.1 | WP_042343900.1 | 35.6 | <1.49E-177 |
| TamL | *Capnocytophaga cynodegmi* | GCF_000379185.1 | WP_026193979.1 | 44.1 | <1.49E-177 |
| TamB | *Capnocytophaga cynodegmi* | GCF_000379185.1 | WP_026193778.1 | 38.8 | <1.49E-177 |
| TamL | *Capnocytophaga gingivalis* | GCF_000174755.1 | WP_002670585.1 | 39.3 | <1.49E-177 |
| TamB | *Capnocytophaga gingivalis* | GCF_000174755.1 | WP_002669221.1 | 37.2 | <1.49E-177 |
| TamL | *Capnocytophaga ochracea* | GCF_000023285.1 | WP_015781761.1 | 41.2 | <1.49E-177 |
| TamB | *Capnocytophaga ochracea* | GCF_000023285.1 | WP_015782581.1 | 37.8 | <1.49E-177 |
| TamL | *Chitinophaga filiformis* | GCF_900102545.1 | WP_089835211.1 | 24.9 | 3.12E-60 |
| TamB | *Chitinophaga filiformis* | GCF_900102545.1 | WP_089833917.1 | 22.7 | 3.73E-85 |
| TamL2 | *Chitinophaga filiformis* | GCF_900102545.1 | WP_176842178.1 | 41.7 | <1.49E-177 |
| TamB2 | *Chitinophaga filiformis* | GCF_900102545.1 | WP_143011371.1 | 31.7 | <1.49E-177 |
| TamL | *Chitinophaga pinensis* | GCF_000024005.1 | WP_012789346.1 | 24.9 | 2.8E-63 |
| TamB | *Chitinophaga pinensis* | GCF_000024005.1 | WP_012793517.1 | 29.6 | 7.33E-05 |
| TamL2 | *Chitinophaga pinensis* | GCF_000024005.1 | WP_012788409.1 | 41.7 | <1.49E-177 |
| TamB2 | *Chitinophaga pinensis* | GCF_000024005.1 | WP_012788408.1 | 31.4 | <1.49E-177 |
| TamL | *Gramella forsetii* | GCF_000060345.1 | WP_011708308.1 | 47.9 | <1.49E-177 |
| TamB | *Gramella forsetii* | GCF_000060345.1 | WP_229664785.1 | 40.9 | <1.49E-177 |
| TamL2 | *Gramella forsetii* | GCF_000060345.1 | WP_011709449.1 | 34.9 | 2.14E-149 |
| TamB2 | *Gramella forsetii* | GCF_000060345.1 | WP_011709448.1 | 27.7 | <1.49E-177 |
| TamA | *Croceibacter atlanticus* | GCF_000196315.1 | WP_148232783.1 | 35.3 | 5.58E-90 |
| TamL | *Croceibacter atlanticus* | GCF_000196315.1 | WP_013188192.1 | 49.7 | <1.49E-177 |
| TamB | *Croceibacter atlanticus* | GCF_000196315.1 | WP_238524714.1 | 42.6 | <1.49E-177 |
| TamL2 | *Croceibacter atlanticus* | GCF_000196315.1 | WP_013186526.1 | 35.6 | 1.9E-150 |
| TamB2 | *Croceibacter atlanticus* | GCF_000196315.1 | WP_013186525.1 | 27.9 | <1.49E-177 |
| TamA | *Cytophaga hutchinsonii* | GCF_000014145.1 | WP_011583574.1 | 23.9 | 1.83E-24 |
| TamL | *Cytophaga hutchinsonii* | GCF_000014145.1 | WP_011585300.1 | 24.9 | 2.3E-47 |
| TamB | *Cytophaga hutchinsonii* | GCF_000014145.1 | WP_011585340.1 | 23.8 | 7.74E-84 |
| TamL2 | *Cytophaga hutchinsonii* | GCF_000014145.1 | WP_011584611.1 | 37.8 | 1.49E-177 |
| TamB2 | *Cytophaga hutchinsonii* | GCF_000014145.1 | WP_041932225.1 | 29.1 | <1.49E-177 |
| TamA | *Flavobacterium columnare* | GCF_007990835.1 | WP_014166290.1 | 42.1 | 1.13E-126 |
| TamL | *Flavobacterium columnare* | GCF_007990835.1 | WP_014164941.1 | 48 | <1.49E-177 |
| TamB | *Flavobacterium columnare* | GCF_007990835.1 | WP_097609278.1 | 47.6 | <1.49E-177 |
| TamA | *Flavobacterium johnsoniae* | GCF_034479105.1 | WP_012022494.1 | 100 | <1.49E-177 |
| TamL | *Flavobacterium johnsoniae* | GCF_034479105.1 | WP_012023543.1 | 100 | <1.49E-177 |
| TamB | *Flavobacterium johnsoniae* | GCF_034479105.1 | WP_012026557.1 | 100 | <1.49E-177 |
| TamL2 | *Flavobacterium johnsoniae* | GCF_034479105.1 | WP_012023976.1 | 100 | <1.49E-177 |
| TamB2 | *Flavobacterium johnsoniae* | GCF_034479105.1 | WP_012023975.1 | 100 | <1.49E-177 |
| TamL | *Elizabethkingia meningoseptica* | GCF_000367325.1 | WP_016199247.1 | 30.3 | 3.69E-94 |

| | | | | | |
|---|---|---|---|---|---|
| TamB | *Elizabethkingia meningoseptica* | GCF_000367325.1 | WP_016200382.1 | 23.1 | 2.19E-86 |
| TamL2 | *Elizabethkingia meningoseptica* | GCF_000367325.1 | WP_016199631.1 | 52.3 | <1.49E-177 |
| TamB2 | *Elizabethkingia meningoseptica* | GCF_000367325.1 | WP_019050827.1 | 44.3 | <1.49E-177 |
| TamA | *Flavobacterium psychrophilum* | GCF_900101925.1 | WP_034099287.1 | 48.1 | 1.01E-168 |
| TamL | *Flavobacterium psychrophilum* | GCF_900101925.1 | WP_034099138.1 | 54 | <1.49E-177 |
| TamB | *Flavobacterium psychrophilum* | GCF_900101925.1 | WP_052079116.1 | 51.1 | <1.49E-177 |
| TamA | *Flavobacterium succinicans* | GCF_000611675.1 | WP_024981504.1 | 46.4 | 4.07E-162 |
| TamL | *Flavobacterium succinicans* | GCF_000611675.1 | WP_024980311.1 | 68.4 | <1.49E-177 |
| TamB | *Flavobacterium succinicans* | GCF_000611675.1 | WP_035717715.1 | 59.2 | <1.49E-177 |
| TamL2 | *Flavobacterium succinicans* | GCF_000611675.1 | WP_035717176.1 | 54.9 | <1.49E-177 |
| TamB2 | *Flavobacterium succinicans* | GCF_000611675.1 | WP_024979783.1 | 43.2 | <1.49E-177 |
| TamL | *Flexibacter flexilis* | GCF_900112255.1 | WP_091512127.1 | 24.1 | 5.57E-49 |
| TamB | *Flexibacter flexilis* | GCF_900112255.1 | WP_091508846.1 | 23.4 | 8.85E-77 |
| TamA | *Kordia algicida* | GCF_000154725.1 | WP_007094993.1 | 37.5 | 5.61E-99 |
| TamL | *Kordia algicida* | GCF_000154725.1 | WP_007096862.1 | 49.2 | <1.49E-177 |
| TamB | *Kordia algicida* | GCF_000154725.1 | WP_007093137.1 | 41.5 | <1.49E-177 |
| TamL2 | *Kordia algicida* | GCF_000154725.1 | WP_007094918.1 | 36.4 | 6.66E-150 |
| TamB2 | *Kordia algicida* | GCF_000154725.1 | WP_007094919.1 | 26.4 | <1.49E-177 |
| TamL | *Polaribacter irgensii* | GCF_000153225.1 | WP_004569772.1 | 37.4 | 4.98E-167 |
| TamB | *Polaribacter irgensii* | GCF_000153225.1 | WP_018945076.1 | 34.6 | <1.49E-177 |
| TamL | *Porphyromonas gingivalis* | GCF_000010505.1 | WP_012457925.1 | 26.5 | 4.56E-69 |
| TamB | *Porphyromonas gingivalis* | GCF_000010505.1 | WP_230847034.1 | 20.9 | 7.31E-63 |
| TamL2 | *Porphyromonas gingivalis* | GCF_000010505.1 | WP_012457279.1 | 29.7 | 2.57E-92 |
| TamB2 | *Porphyromonas gingivalis* | GCF_000010505.1 | WP_043876309.1 | 21.4 | 7.39E-56 |
| TamL | *Prevotella intermedia* | GCF_000439065.1 | WP_028905715.1 | 25.3 | 1.92E-60 |
| TamB | *Prevotella intermedia* | GCF_000439065.1 | WP_028905192.1 | 21.9 | 4.49E-49 |
| TamL2 | *Prevotella intermedia* | GCF_000439065.1 | WP_028905308.1 | 28.6 | 2.81E-90 |
| TamB2 | *Prevotella intermedia* | GCF_000439065.1 | WP_028905307.1 | 23.7 | 6.14E-39 |
| TamL | *Prevotella melaninogenica* | GCF_000144405.1 | WP_013265768.1 | 25.1 | 2.57E-57 |
| TamB | *Prevotella melaninogenica* | GCF_000144405.1 | WP_013265564.1 | 22.1 | 4.31E-53 |
| TamL2 | *Prevotella melaninogenica* | GCF_000144405.1 | WP_088582096.1 | 27.6 | 1.59E-77 |
| TamB2 | *Prevotella melaninogenica* | GCF_000144405.1 | WP_044046063.1 | 21.4 | 4.11E-34 |
| TamA | *Riemerella anatipestifer* | GCF_000183155.1 | WP_004916258.1 | 27.2 | 3.39E-39 |
| TamL | *Riemerella anatipestifer* | GCF_000183155.1 | WP_013446822.1 | 29.8 | 3.98E-101 |
| TamB | *Riemerella anatipestifer* | GCF_000183155.1 | WP_004918022.1 | 23.5 | 8.76E-90 |
| TamL2 | *Riemerella anatipestifer* | GCF_000183155.1 | WP_004917060.1 | 46.7 | <1.49E-177 |
| TamB2 | *Riemerella anatipestifer* | GCF_000183155.1 | WP_004917063.1 | 39.7 | <1.49E-177 |
| TamL | *Sphingobacterium mizutaii* | GCF_007990895.1 | WP_236736519.1 | 33.1 | 5.59E-35 |
| TamB | *Sphingobacterium mizutaii* | GCF_007990895.1 | WP_236736499.1 | 23.3 | 3.14E-84 |
| TamL2 | *Sphingobacterium mizutaii* | GCF_007990895.1 | WP_236736426.1 | 43.7 | <1.49E-177 |
| TamB2 | *Sphingobacterium mizutaii* | GCF_007990895.1 | WP_236736427.1 | 37.2 | <1.49E-177 |
| TamA | *Sporocytophaga myxococcoides* | GCF_000426725.1 | WP_156027096.1 | 22.9 | 2.6E-22 |
| TamL | *Sporocytophaga myxococcoides* | GCF_000426725.1 | WP_051313097.1 | 27.1 | 1.46E-62 |
| TamB | *Sporocytophaga myxococcoides* | GCF_000426725.1 | WP_028979539.1 | 27.4 | 4.5E-54 |
| TamB2 | *Sporocytophaga myxococcoides* | GCF_000426725.1 | WP_028981824.1 | 22.4 | 1.24E-75 |
| TamA | *Xanthomarina gelatinilytica* | GCF_000348685.1 | WP_238307659.1 | 34.3 | 5.4E-80 |
| TamL | *Xanthomarina gelatinilytica* | GCF_000348685.1 | WP_007648347.1 | 49 | <1.49E-177 |
| TamB | *Xanthomarina gelatinilytica* | GCF_000348685.1 | WP_238307618.1 | 42.9 | <1.49E-177 |
| TamL2 | *Xanthomarina gelatinilytica* | GCF_000348685.1 | WP_007647185.1 | 36.4 | 2.54E-145 |
| TamB2 | *Xanthomarina gelatinilytica* | GCF_000348685.1 | WP_238307542.1 | 25.9 | 1.72E-172 |
| TamL | *Zobellia galactanivorans* | GCF_000973105.1 | WP_013992141.1 | 45.7 | <1.49E-177 |
| TamB | *Zobellia galactanivorans* | GCF_000973105.1 | WP_013995062.1 | 39.8 | <1.49E-177 |
| TamL2 | *Zobellia galactanivorans* | GCF_000973105.1 | WP_013992464.1 | 37.9 | 7.35E-158 |
| TamB2 | *Zobellia galactanivorans* | GCF_000973105.1 | WP_046287358.1 | 28.1 | <1.49E-177 |

**Supplementary Table S4. List of strains used in this study.**

| Name | Genotype | Reference |
|---|---|---|
| ***Escherichia coli* strains** | | |
| Top10 | F- *mcrA Δ(mrr-hsdRMS-mcrBC) φ80lacZΔM15 ΔlacX74 recA1 araD139 Δ(araleu)7697 galU galK rpsL endA1 nupG; Smr* | Invitrogen |
| DH10B | F- *mcrA* Δ(*mrr-hsdRMS-mcrBC*) φ80*lacZ*ΔM15 Δ*lacX74 recA1 endA1 araD139* Δ (*ara-leu*)7697 *galU galK* λ– *rpsL*($Str^R$) *nupG* | Thermo Fisher Scientific |
| MT607 | *pro-82 thi-I hsdR17 (r-m+) supE44 recA56* | [8] |
| ***Flavobacterium johnsoniae* strains** | | |
| Wild type (WT) | *Flavoacterium johnsoniae* UW101 | [9] |
| Δ*tamA* | Deletion of *fjoh_0402* | This study |
| Δ*tamL2* | Deletion of *fjoh_1900* | This study |
| Δ*tamB2* | Deletion of *fjoh_1899* | This study |
| Δ*tamB2*Δ*tamL2* | Deletion of *fjoh*_1899 and of *fjoh_1900* | This study |
| (3xFLAG)*tamL* | 3xFLAG-tag flanked upstream and downstream by 4x Glycine (4Gly) residues added after residues $^{574}$TNQV$^{577}$ and followed by the repetition of the same residues, thus originating $^{574}$TNQV$^{577}$-4Gly-3xFLAG-tag-4Gly-$^{608}$TNQV$^{611}$ | This study |
| (2xStrep)*tamB* | Twin-Strep-tag added at the C-terminus of *tamB* | This study |
| (3xFLAG)*tamL*/(2xStrep)*tamB* | Combination of (3xFLAG)*tamL* and of (2xStrep)*tamB* | This study |
| $P_{ompA}$::*lacI* | $P_{ompA}$::*lacI* construct added within the intergenic region between *fjoh_0061* and *fjoh_0062* | This study |
| $P_{cfxA\text{-}lacO}$::*tamL* | $P_{cfxA\text{-}lacO}$ construct added downstream of the putative native promoter of *tamL* | This study |
| $P_{ompA}$::*lacI*-$P_{cfxA\text{-}lacO}$::*tamL* | TamL depletion strain in the (3xFLAG)*tamL*/(2xStrep)*tamB* background | This study |
| Δ*tamA*-$P_{ompA}$::*lacI*-$P_{cfxA\text{-}lacO}$::*tamL* | Deletion of *fjoh_0402* in $P_{ompA}$::*lacI*-$P_{cfxA\text{-}lacO}$::*tamL* background | This study |
| Δ*tamL2*-$P_{ompA}$::*lacI*-$P_{cfxA\text{-}lacO}$::*tamL* | Deletion of *fjoh_1900* in $P_{ompA}$::*lacI*-$P_{cfxA\text{-}lacO}$::*tamL* background | This study |
| Δ*tamA*Δ*tamL2*-$P_{ompA}$::*lacI*-$P_{cfxA\text{-}lacO}$::*tamL* | Deletion of *fjoh_0402* and *fjoh_1900* in $P_{ompA}$::*lacI*-$P_{cfxA\text{-}lacO}$::*tamL* background | This study |
| Δ*fjoh_2419* | Deletion of *fjoh_2419* | This study |
| Δ*fjoh_0833* | Deletion of *fjoh_0833* | This study |
| Δ*fjoh_0833*Δ*fjoh_2419* | Deletion of *fjoh_2419* and of *fjoh_0833* | This study |
| Δ*fjoh_2419*-$P_{ompA}$::*lacI*-$P_{cfxA\text{-}lacO}$::*tamL* | Deletion of *fjoh_2419* in $P_{ompA}$::*lacI*-$P_{cfxA\text{-}lacO}$::*tamL* background | This study |
| Δ*fjoh_0833*-$P_{ompA}$::*lacI*-$P_{cfxA\text{-}lacO}$::*tamL* | Deletion of *fjoh_0833* in $P_{ompA}$::*lacI*-$P_{cfxA\text{-}lacO}$::*tamL* background | This study |

| | | |
|---|---|---|
| Δ*fjoh_0833*Δ*fjoh_2419*-$P_{ompA}$::*lacI*-$P_{cfxA\text{-}lacO}$::*tamL* | Deletion of *fjoh_2419* and of *fjoh_0833* in $P_{ompA}$::*lacI*-$P_{cfxA\text{-}lacO}$::*tamL* background | This study |

**Supplementary Table S5. List of plasmids used in this study.**

| Plasmid name | Description | Reference |
|---|---|---|
| **Vectors** | | |
| pYT354 | *sacB*-containing suicide vector; Apr (Emr) | [10] |
| pBAD33 | pACYC *ori*; CmR. Low copy *E. coli* expression plasmid with arabinose inducible promoter | [11] |
| pCP23 | ColE1 *ori*; (pCP1 ori); Apr(Tcr); *E. coli-F. johnsoniae* shuttle plasmid | [12] |
| pMM47.A | ColE1 *ori*; (pCC7 *ori*); Apr; (Cfxr). *E. coli-C. canimorsus* expression shuttle plasmid with *ermF* promoter | [13] |
| **Suicide plasmids** | | |
| pYT354-Δ*tamL* | Deletion of Δ*fjoh_1464*. Upstream and downstream regions (2 kb) of *fjoh_1464* were PCR-amplified with oligonucleotides 8526/ 8439 and 8440/8527, respectively, from gDNA, PCR-overlapped and then cloned into pYT354 using ApaI/SpeI restriction sites. | This study |
| pYT354-Δ*tamB* | Deletion of Δ*fjoh_4592*. Upstream and downstream regions (2 kb) of *fjoh_4592* were PCR-amplified with oligonucleotides 8356/8522 and 8378/8359, respectively, from gDNA, PCR-overlapped and then cloned into pYT354 using BamHI/SpHI restriction sites. | This study |
| pYT354-Δ*fjoh_0402* | Deletion of *fjoh_0402*. Upstream and downstream regions (2 kb) of *fjoh_0402* were PCR-amplified with oligonucleotides FG179_KOUp_Fw/FG185_KOUp_Rv and FG181_KODown_Fw/FG187_KODown_Rv, respectively, from gDNA and cloned sequentially into pYT354 using SphI/XhoI and XhoI/XbaI restriction sites. | This study |
| pYT354-Δ*fjoh_1900* | Deletion of *fjoh_1900*. Upstream and downstream regions of *fjoh_1900* (2 kb) were PCR-amplified with oligonucleotides FR634 UP_Fw BamHI/FG1 UP_Rev KpnI and FG2 Dwn_Fw KpnI/FR637 Dwn_Rev SphI, respectively, from gDNA and cloned sequentially into pYT354 using BamHI/KpnI and KpnI/SpHI restriction sites. | This study |
| pYT354-Δ*fjoh_1899* | Deletion of *fjoh_1899*. Upstream and downstream regions of *fjoh_1899* (2 kb) were amplified with oligonucleotides FG17_KOFj_1899Fw/FG18_KOFj_1899Rv and FG19_KOFj_1899Fw/FG20_KOFj_1899Rv, respectively, from gDNA; pYT354 plasmid was amplified with oligonucleotides FG21_KOFj_1899Fw/FG22_KOFj_1899Rv from pYT354 plasmid. All fragments were overlapped and cloned in pYT354 by Gibson assembly [14]. | This study |

| | | |
|---|---|---|
| pYT354-Δ*fjoh_2419* | Deletion of *fjoh_2419*. Upstream and downstream regions (2 kb) of *fjoh_2419* were amplified with oligonucleotides 8815/8816 and 8817/8818, respectively, from gDNA; pYT354 plasmid was amplified with oligonucleotides 8819/8820 from pYT354 plasmid. All fragments were overlapped and cloned in pYT354 by Gibson assembly [14]. | |
| pYT354-Δ*fjoh_0833* | Deletion of *fjoh_0833*. Upstream and downstream regions (2 kb) of *fjoh_0833* were PCR-amplified with oligonucleotides FG204_1stfr_SphI/FG205_1stfr_XhoI and FG206_2ndfr_XhoI/FG207_2ndfr_XbaI, respectively, from gDNA and cloned sequentially into pYT354 using SphI/XhoI and XhoI/XbaI restriction sites. | This study |
| pYT354-$tamL_{21cys\text{-}gly}$ | $^{21}$Cys of TamL replaced with Gly. Upstream and downstream regions (2 kb) of $fjoh\_1464_{21Cys}$ were PCR-amplified with oligonucleotides 8534/8555 and 8556/8535, respectively, from gDNA and cloned into pYT354 using ApaI/SpeI restriction sites. | This study |
| pYT354-*fjoh*_1464-$^{574}$TNQV$^{577}$-4xGly-3xFLAG-tag-4xGly-$^{608}$TNQV$^{611}$ | Insertion of a 3xFLAG-tag sequence, flanked upstream and downstream by 4 Glycine residues (4xGly), inserted after residues $^{574}$TNQV$^{577}$ of TamL and followed by the repetition of the same residues. Upstream and downstream regions of *tamL*-$^{574}$TNQV$^{577}$ (2 kb) were PCR-amplified with oligonucleotides 3loop_1fragm_Fw/FG57_FjhTamL1_Rv and FG70_FjhTamL1_Fw/3loop_3fragm_Rev, respectively, from gDNA. In parallel, the 3xFLAG-tag sequence flanked upstream and downstream by 4xGly residues was PCR-amplified using oligonucleotides FG51_4xG3xFlagFw/FG52_4xG3xFlagRv from plasmid pNPTS138 lptD::lptD-3Flag [15]. pYT354 plasmid was amplified with oligonucleotides 3loop_4fragm_Fw/3loop_4fragm_Rev from pYT354 plasmid. All the fragments were overlapped and cloned into pYT354 by Gibson assembly [14]. | This study |
| pYT354-*fjoh_4592*-2xStrep | Twin-Strep-tag sequence fused in frame with the C-terminus of TamB. Upstream and downstream regions (2 kb) of the C-terminus of TamB were PCR-amplified with oligonucleotides FG61_FjTamB1_Fw/FG62_FjTamB1_Rv and FG63_FjTamB1_Fw/FG64_FjTamB1_Rv, respectively, from gDNA. In parallel, the 2xStrep-tag sequence was PCR-amplified using oligonucleotides LL03-2Strep-fw and FG69_2xStrep_Rv from pYT313-SprE-TAG (plasmid received from Ben Berg`s lab). pYT354 plasmid was amplified with | This study |

| | | |
|---|---|---|
| | oligonucleotides FG65_FjTamB1_Fw/FG66_FjTamB1_Rv from pYT354 plasmid. All the fragments were overlapped and cloned into pYT354 by Gibson assembly [14]. | |
| pYT354-*fjoh_0061-fjoh_0062* | The 2 kb-genomic regions of *fjoh_0061* and *fjoh_0062* were PCR-amplified with oligonucleotides 8580/8581 and 8582/8583, respectively, from gDNA, and cloned sequentially into pYT354 using ApaI/SalI and XhoI/BamHI and restriction sites. | This study |
| pYT354-$P_{ompA}$::*lacI* | Insertion of the $P_{ompA}$::*lacI* construct within the intergenic region between *fjoh_0061* and *fjoh_0062*. The $P_{ompA}$::*lacI* construct was PCR-amplified using oligonucleotides 8882/8883 from pCP23-$P_{ompA}$::*lacI,* and cloned into pYT354-*fjoh_0061-fjoh_0062* using XhoI restriction site. | This study |
| pYT354-$P_{cfxA-lacO}$::*tamL* | Insertion of the $P_{cfxA-lacO}$::*tamL* construct upstream of *tamL*. Upstream and downstream regions (2 kb) of *tamL* start codon were PCR-amplified with oligonucleotides FG154_1stfrag_Fw/FG155_1stfrag_Rv and FG157_3rdfrag_Fw/FG158_3rdfrag_Rv, respectively, from gDNA. In parallel, the $P_{cfxA-lacO}$ construct was PCR-amplified using oligonucleotides FG156_2ndfrag_Fw/FR806-PcfxlacORv from pFL32. pYT354 plasmid was amplified with oligonucleotides FG159_4thfrag_Fw/FG160_4thfrag_Rv from pYT354 plasmid. All the fragments were cloned into pYT354 by Gibson assembly [14]. | This study |
| pYT354-$P_{cfxA-lacO}$::*tamB* | Insertion of the $P_{cfxA-lacO}$::*tamB* construct upstream of *tamB*. Upstream and downstream regions (2 kb) of *tamB* start codon were PCR-amplified with oligonucleotides FG152_1stfrag_Fw/FG146_1stfrag_Rv and FG148_3rdfrag_Fw/FG149_3rdfrag_Rv, respectively, from gDNA. In parallel, the $P_{cfxA-lacO}$ construct was PCR-amplified using oligonucleotides FR789_PcfxH2fw/FG147_2ndfrag_Rv from pFL32. pYT354 plasmid was amplified with oligonucleotides FG150_4thfrag_Fw/FG153_4thfrag_Rv from pYT354 plasmid. All the fragments were cloned into pYT354 by Gibson assembly [14]. | This study |
| pYT354-$P_{ara}$::*tamB* | Insertion of the $P_{ara}$::*tamB* construct upstream of *tamB*. Upstream and downstream regions of *tamB* start codon (2 kb) were PCR-amplified with oligonucleotides FG152_1stfrag_Fw/FG173frg1_ParaRv and FG178frg3_ParaFw/FG170_5thfrag_Rv, respectively, from pYT354-$P_{cfxA-lacO}$::*tamB.* In parallel, the $P_{ara}$ construct was PCR-amplified using oligonucleotides | This study |

| | | |
|---|---|---|
| | FG177frg2_ParaFw/FG175frg2_ParaRv from gDNA. pYT354 plasmid was amplified with oligonucleotides FG171_4thfrag_Fw/FG153_4thfrag_Rv from pYT354 plasmid. All the fragments were cloned into pYT354 by Gibson assembly [14]. | |
| **Expression plasmids** | | |
| pCP23-$P_{ompA}$::*lacI* | The $P_{ompA}$ sequence was PCR-amplified with oligonucleotides 7146/7147 from gDNA. The *lacI* sequence was PCR-amplified with oligonucleotides 7148/7149 from *E. coli* MG1655 gDNA. PCR fragments were overlapped and then cloned into pCP23 using HindIII/SphI restriction sites. | This study |
| pFL32 | *lacO3-PcfxA-lacO1* amplified from pFD1146 with primers 7144/7145 and cloned into pMM47.A using SalI/NcoI restriction sites. | [13] |

**Supplementary Table S6. List of oligonucleotides used in this study.** Nucleotides corresponding to restriction sites are shown as underlined in lower case and preceded by the enhancer cut sequence in italics. Nucleotides corresponding to overlapping regions for Gibson assembly [14] are in lower case, while those annealing with the DNA/plasmid template are in upper case.

| **Name** | **Sequence 5'-3'** |
|---|---|
| FG179_KOUp_Fw | *acat*gcatgcATCATCAGGATGAATACCTTCAATACCTTC |
| FG185_KOUp_Rv | *ccg*ctcgagGAATAAAGTTTTTGTTTGGTTATATGTGATGC |
| FG181_KODown_Fw | *ccg*ctcgagTTCAAATACCATAAAACACAAACACC |
| FG187_KODown_Rv | *cgc*ggatccTTGTAAGTAAAGTTTGCGTAACCTATG |
| FR634 UP_Fw BamHI | *gtc*ggatccTTTAATTAACGGAAATGCTCTGGTC |
| FG1 UP_Rev KpnI | ggggtaccTTATGTTTTTTGTTCATTTCC |
| FG2 Dwn_Fw KpnI | ggggtaccGTTTTAAGTTTTTCGCCACG |
| FR637 Dwn_Rev SphI | *gcc*gcatgcTAAAGAAAATCCCTTATGATATTTC |
| FG17_KOFj_1899Fw | ccgctctagaactagtggatTGTAATTCAGAGAAGCATTC |
| FG18_KOFj_1899Rv | catttccttcaatttgatttAGGTCAATAAAATACAGGAT |
| FG19_KOFj_1899Fw | atcctgtattttattgacctAAATCAAATTGAAGGAAATG |
| FG20_KOFj_1899Rv | tgatatcgaattcctgcagcACAATACTAAACAATTTTGC |
| FG21_KOFj_1899Fw | gcaaaattgtttagtattgtGCTGCAGGAATTCGATATCA |
| FG22_KOFj_1899Rv | gaatgcttctctgaattacaATCCACTAGTTCTAGAGCGG |
| 8815 | gttatgcagcggaaaaattcgggAAAATCGCTTTTCCAATACGACAG |
| 8816 | gagaatttatttttaaatcaAAATATAGTGTAGTAATTTTTTAACTTAATCAGCGAGC |
| 8817 | aaaattactacactatatttTGATTTAAAAATAAATTCTCACATGGGAATTATG |
| 8818 | atgaccatgattacgccaagcttTATGACGGATGGTATAAAGAGAAAG |
| 8819 | ttataccatccgtcataAAGCTTGGCGTAATCATGGTCATAG |
| 8820 | attggaaaagcgattttCCCGAATTTTTCCGCTGCATAACC |
| FG204_1stfr_SphI | *acat*gcatgcAGGCTCAAAAGTTAAAACCTTTGAG |
| FG205_1stfr_XhoI | *ccg*ctcgagTGCCTTTGCAACTTCTTTCGC |
| FG206_2ndfr_XhoI | *ccg*ctcgagTACATTAGAATCGCAGATATTCCTGAAAG |
| FG207_2ndfr_XbaI | *gc*tctagaATGGAAACTACAAAACCAGAATCATGG |
| 8534 | *at*gggcccGAAGTGGGAAAAAAGAAAGTTGAG |
| 8555 | ACAGCATTccGGCtccAATAAGTATTGCTATTAGAATAAATGCTGT |
| 8556 | TACTTATTggaGCCggaAATGCTGTAAAAAGAGTTCCTGA |
| 8535 | *tgcact*actagtTTTCCACATCTTTATATTGTGGATTATC |
| 8526 | *at*gggcccAGGAAAAAAAACTCTCTAATGTAATTGATCTG |
| 8439 | taggttgGTGTTTTTTAATATTTAATTCAAAAGTACATTATTTTATGGT |
| 8440 | gaattaaatattaaaaaacacCAACCTAAAAACTAAAAACAACTATAAAAAAAATTAC |
| 8527 | *tgcact*actagtCAATTCTTTTCTAGGCGTGATAAAGTC |
| 8356 | *gtc*ggatccATTTTTTTCATAGAATTGTAATTTG |
| 8522 | acaaattatAAGTAAAATTAAGCCAATCAG |

| | |
|---|---|
| 8378 | gattggcttaattttacttATAATTTGTTAAAAAGAGTATAAAAAAACCATTC |
| 8359 | *gcc*gcatgcTGATTGCAGCTGCACGTC |
| 3loop_1fragm_Fw | ccgctctagaactagtggatCTAAAGCCAGAATTAAACCAG |
| FG57_FjhTamL1_Rv | tataatcaccgtcatggtctttgtagtcgccgccgccgccTACTTGATTCGTGAAATTTGTAGTTC |
| FG70_FjhTamL1_Fw | catcgattacaaggatgacgatgacaagggcggcggcggcACGAATCAAGTACTAACCAATCAAACAGCTTTAACAC |
| 3loop_3fragm_Rev | tgatatcgaattcctgcagcGGAACGTCCATCTTATCTTC |
| FG51_4xG3xFlagFw | GGCGGCGGCGGCGACTACAAAGACCATGACGGTG |
| FG52_4xG3xFlagRv | GCCGCCGCCGCCCTTGTCATCGTCATCCTTGTAATC |
| 3loop_4fragm_Fw | gaagataagatggacgttccGCTGCAGGAATTCGATATCA |
| 3loop_4fragm_Rev | ctggtttaattctggctttagATCCACTAGTTCTAGAGCGG |
| FG61_FjTamB1_Fw | ccgctctagaactagtggatCAAAACAGCCGTCCTTTGTCTG |
| FG62_FjTamB1_Rv | ctccggaacctccaccttttcgaactgcgggtggctccaAAAATCATTGTCAGGAATTAAGCCTTC |
| FG63_FjTamB1_Fw | ggcgtggtcacatccacaatttgagaagtagATAATTTGTTAAAAAGAGTATAAAAAAACCATTCG |
| FG64_FjTamB1_Rv | tgatatcgaattcctgcagcGATAACTTTTCCAACCGCTTTAGCAG |
| LL03-2Strep-fw | TGGAGCCACCCGCAGTTCGAAAAAG |
| FG69_2xStrep_Rv | CTTCTCAAATTGTGGATGTGACCAC |
| FG65_FjTamB1_Fw | ctgctaaagcggttggaaaagttatcGCTGCAGGAATTCGATATCAAGC |
| FG66_FjTamB1_Rv | cagacaaaggacggctgttttgATCCACTAGTTCTAGAGCGGC |
| 8580 | *gt*gggcccAGACCATTACGCTTGATAACATGA |
| 8581 | ctcgagTCAGAgtcgacAAGATGGCAAAAATGAGTATAAATCTG |
| 8582 | GCCATCTTgtcgacTCTGActcgagGGGGAACGAAGCAGCCAC |
| 8583 | *caga*ggatccGTAAAGGTACTTAAACAATGTTTACCTTTATGGC |
| 8882 | *ccg*ctcgagTCACTGCCCGCTTTCCAGTCGG |
| 8883 | tcgagTTTTTTTTAACATTTGATTTTGTATTTA |
| FG154_1stfrag_Fw | ccgctctagaactagtggatCCGGGGACTTAGCTATTTTAATGG |
| FG155_1stfrag_Rv | atttcgggatttttttgcaGTGTTTTTTAATATTTAATTCAAAAGTACATTATTTTATGG |
| FG157_3rdfrag_Fw | gttatgtacctttgtcggcaattgtgagcggataacaattTTGAAAAATAATTCCACAAAAAATAACAGC |
| FG158_3rdfrag_Rv | tgatatcgaattcctgcagcCAAAAATAAGAATTAATTAGCATTAGAACGG |
| FG156_2ndfrag_Fw | aattaaatattaaaaaacacTGCAAAAAAAATCCCGAAATAAATTCGG |
| FR806-PcfxlacORv | AATTGTTATCCGCTCACAATTGCCG |
| FG159_4thfrag_Fw | ctaattaattcttatttttgGCTGCAGGAATTCGATATCAAGC |
| FG160_4thfrag_Rv | taaaatagctaagtccccggATCCACTAGTTCTAGAGCGGC |
| FG152_1stfrag_Fw | ccgctctagaactagtggatATTTTTTTCATAGAATTGTAATTTGTCTACTCGTTTTG |
| FG146_1stfrag_Rv | ctgcccaaacaaaaaaatcccgaatttatttcgggatttttttgcaAAGTAAAATTAAGCCAATCAGGGTTCTG |
| FG148_3rdfrag_Fw | attgtgagcggataacaattTTGCTGATACTTGCTATCACTCTGTC |

| | |
|---|---|
| FG149_3rdfrag_Rv | tgatatcgaattcctgcagcCAGGTCTGCATCAATTTTACCGC |
| FR789_PcfxH2fw | aaaatcccgaaataaattcgggatttttttgtttgGGCAGTGAGCGCAACGCAATTTTAC |
| FG147_2ndfrag_Rv | gtgatagcaagtatcagcaaAATTGTTATCCGCTCACAATTGCCG |
| FG150_4thfrag_Fw | gcggtaaaattgatgcagacctgGCTGCAGGAATTCGATATCAAGC |
| FG153_4thfrag_Rv | tacaattctatgaaaaaaatATCCACTAGTTCTAGAGCGGC |
| FG173frg1_ParaRv | aggaggtaaattgatctctacaaacaaaaaaaTCCCGAATTTATTTCGGG |
| FG178frg3_ParaFw | ttttataagtatatttgtcaccattaaaaataaatttcgATTGCTGATACTTGCTATCACTCTG |
| FG170_5thfrag_Rv | tgatatcgaattcctgcagcGAAGTGTTATAAAAATCTCACTTAGAG |
| FG177frg2_ParaFw | tgcaaaaaaaaatcccgaaataaattcgggatttttttgtttgTAGAGATCAATTTACCTCCTTTTTCAG |
| FG175frg2_ParaRv | gtgatagcaagtatcagcaaTCGAAATTTATTTTTAATGGTGACAAATATAC |
| FG171_4thfrag_Fw | taagtgagatttttataacacttCGCTGCAGGAATTCGATATCAAGC |
| 7146 | ggaagcTTTTTTTTTTAACATTTGATTTTG |
| 7147 | gtataacgttACTGGTTTCATACTTAATTTTTTAATTA |
| 7148 | gtaattaaaaaaattaagtATGAAACCAGTAACGTTATACG |
| 7149 | gggcatgcTCACTGCCCGCTTTCCAGTCGG |
| 7144 | gggtcgacGGCAGTGAGCGCAACGC |
| 7145 | ggccatggAATTGTTATCCGCTCACAATTGC |

#### References

1. Okino N, Ito M. Thin-layer chromatography (TLC) of glycolipids. In: Nishihara S, Angata K, Aoki-Kinoshita KF, Hirabayashi J, editors. Saitama (JP); 2021.

2. Vences-Guzmán MÁ, Peña-Miller R, Hidalgo-Aguilar NA, Vences-Guzmán ML, Guan Z, Sohlenkamp C. Identification of the Flavobacterium johnsoniae cysteate-fatty acyl transferase required for capnine synthesis and for efficient gliding motility. Environ Microbiol. 2021;23: 2448–2460. doi:10.1111/1462-2920.15445

3. Evans R, O'Neill M, Pritzel A, Antropova N, Senior A, Green T, et al. Protein complex prediction with AlphaFold-Multimer. bioRxiv. 2022; 2021.10.04.463034. doi:10.1101/2021.10.04.463034

4. Teufel F, Almagro Armenteros JJ, Johansen AR, Gíslason MH, Pihl SI, Tsirigos KD, et al. SignalP 6.0 predicts all five types of signal peptides using protein language models. Nat Biotechnol. 2022;40: 1023–1025. doi:10.1038/s41587-021-01156-3

5. Drula E, Garron ML, Dogan S, Lombard V, Henrissat B, Terrapon N. The carbohydrate-active enzyme database: Functions and literature. Nucleic Acids Res. 2022;50: D571–D577. doi:10.1093/nar/gkab1045

6. Vallenet D, Labarre L, Rouy Z, Barbe V, Bocs S, Cruveiller S, et al. MaGe: a microbial genome annotation system supported by synteny results. Nucleic Acids Res. 2006;34: 53–65. doi:10.1093/nar/gkj406

7. Boratyn GM, Schäffer AA, Agarwala R, Altschul SF, Lipman DJ, Madden TL. Domain enhanced lookup time accelerated BLAST. Biol Direct. 2012;7: 1–14. doi:10.1186/1745-6150-7-12

8. P. A, C. CT. Poly-3-Hydroxybutyrate Degradation in Rhizobium (Sinorhizobium) meliloti: Isolation and Characterization of a Gene Encoding 3-Hydroxybutyrate Dehydrogenase. J Bacteriol. 1999;181: 849–857. doi:10.1128/jb.181.3.849-857.1999

9. McBride MJ, Xie G, Martens EC, Lapidus A, Henrissat B, Rhodes RG, et al. Novel features of the polysaccharide-digesting gliding bacterium Flavobacterium johnsoniae as revealed by genome sequence analysis. Appl Environ Microbiol. 2009;75: 6864–6875. doi:10.1128/AEM.01495-09

10. Zhu Y, Thomas F, Larocque R, Li N, Duffieux D, Cladière L, et al. Genetic analyses unravel the crucial role of a horizontally acquired alginate lyase for brown algal biomass degradation by Zobellia galactanivorans. Environ Microbiol. 2017;19: 2164–2181. doi:10.1111/1462-2920.13699

11. Guzman LM, Weiss DS, Beckwith J. Domain-swapping analysis of FtsI, FtsL, and FtsQ, bitopic membrane proteins essential for cell division in Escherichia coli. J Bacteriol. 1997;179: 5094–5103. doi:10.1128/jb.179.16.5094-5103.1997

12. Agarwal S, Hunnicutt DW, McBride MJ. Cloning and characterization of the Flavobacterium johnsoniae (Cytophaga johnsonae) gliding motility gene, gldA. Proc Natl Acad Sci U S A. 1997;94: 12139–12144. doi:10.1073/pnas.94.22.12139

13. Mally M, Cornelis GR. Genetic tools for studying Capnocytophaga canimorsus. Appl Environ Microbiol. 2008;74: 6369–6377. doi:10.1128/AEM.01218-08

14. Gibson DG, Young L, Chuang RY, Venter JC, Hutchison CA, Smith HO. Enzymatic

assembly of DNA molecules up to several hundred kilobases. Nat Methods. 2009;6: 343–345. doi:10.1038/nmeth.1318

15. Servais C, Vassen V, Verhaeghe A, Küster N, Carlier E, Phégnon L, et al. Lipopolysaccharide biosynthesis and traffic in the envelope of the pathogen Brucella abortus. Nat Commun. 2023;14: 911. doi:10.1038/s41467-023-36442-y
